# Ancestry, admixture, and pathogens in contemporaneous Neolithic farmers and foragers on the Island of Gotland

**DOI:** 10.64898/2025.12.07.692553

**Authors:** Magdalena Fraser, Federico Sanchez-Quinto, Emrah Kırdök, Kristiina Ausmees, Gulsah Merve Kılınç, Maximilian Larena, Leonardo Correa-Mendoza, Adrien Le Meur, Antonio Blanchet, Nora Bergfeldt, Eduardo Arrieta-Donato, Mariana Escobar-Rodríguez, Anders Götherström, Karla Lozano-Gonzalez, Israel Aguilar-Ordoñez, Helena Malmström, Kjel Knutsson, Paul Wallin, Nicolas Rascovan, Jan Storå, Mattias Jakobsson

## Abstract

Two archaeological cultural complexes coexisted on Gotland for over 500 years, between ∼3300 and 2800 calBCE, i.e. the Neolithic Funnelbeaker culture (FBC), and the Pitted ware culture (PWC). The ancestry of the FBC farmers and PWC marine foragers largely aligns with European Neolithic Farmers and European Mesolithic foragers, respectively, but the direct interactions between the groups on Gotland is not understood. We present a Middle Neolithic (MN) high-coverage genome and a Late Neolithic (LN) low-coverage genome from the Ansarve FBC dolmen. We investigate ancestry, admixture, and pathogens among these MN farmers (n =6), foragers (n=19), and a LN individual. We find that recent gene-flow between farmers and foragers could have taken place, although most gene-flow happened prior to their coexistence on the island. We also find evidence of different *Yersinia pestis* strains in the three cultural groups, showing that the pestis was widespread among groups with different subsistence strategies.

## Introduction

Due to the rapid advancement in archaeogenomics, we now know that some of the present-day patterns of genomic variation in Northern Europe were shaped by several major pre-historic demographic events^1^. These include the Palaeolithic and Mesolithic post-glacial dispersal of the pioneer hunter-gatherer groups northwards over the continent, the displacement and assimilation of Mesolithic hunter-gatherers (HGs) during the expansion of Early Neolithic farmer (ENF) groups^2^, followed by the expansion of groups of the Corded Ware complex (CWC) bringing the so-called Steppe-ancestry genetic component into the area^3,4^.

Although the overarching patterns of migration during the European Neolithic are known, the demographic and social development in many local regions of Europe have not been fully explored and may therefore show different and unique patterns of interactions between foragers and farmers. The island of Gotland [Fig. 1] is unique in this way; though situated in the middle of the Baltic Sea, it mirrors the general Neolithic developments seen in southern Scandinavia^2,5–9^, but also displays local developments^10–14^.

**Figure 1.**
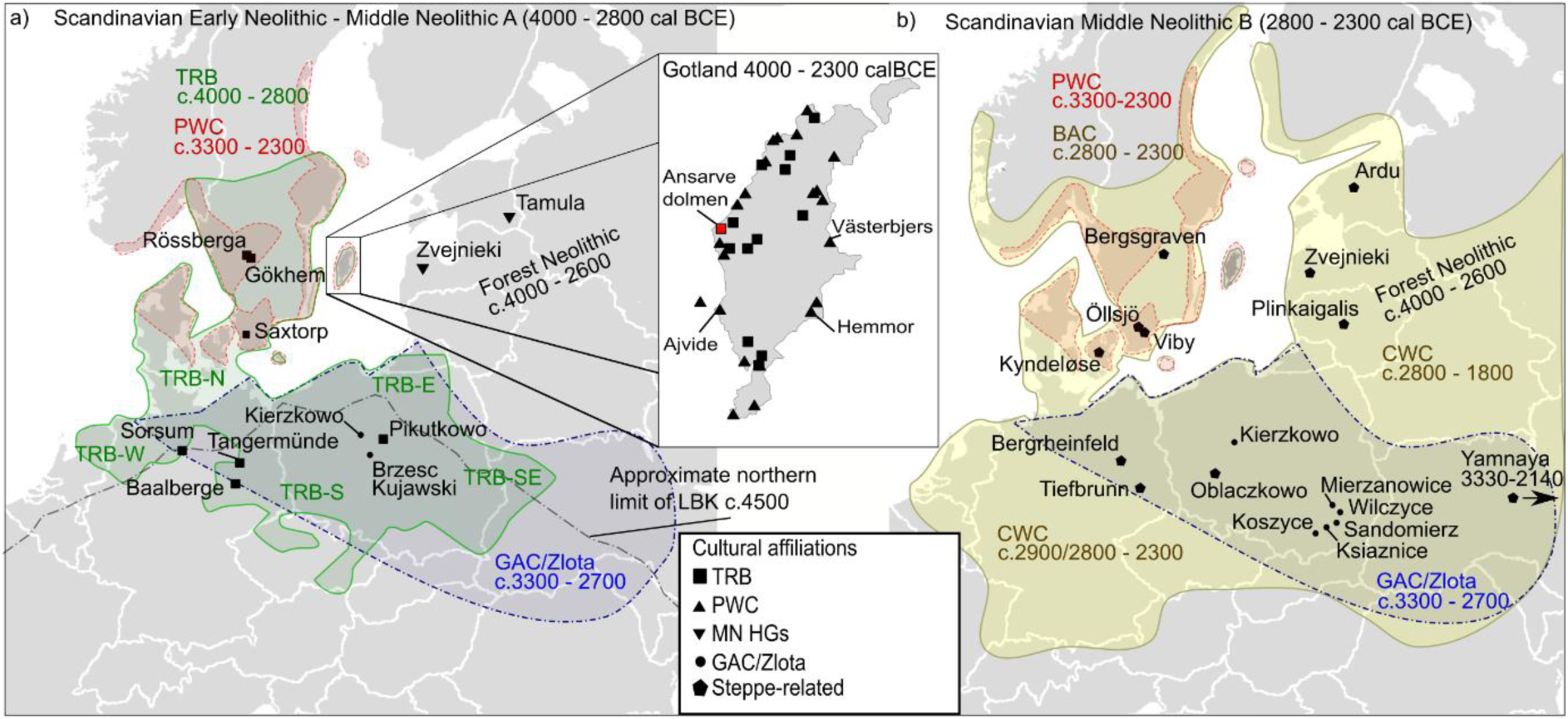
Map of Northern and Central Europe with approximate distribution and timelines for the TRB (Trichterbecher) culture, PWC (Pitted Ware) culture, Forest Neolithic cultures, GAC (Globular Amphorae)/Zlota cultures, and BAC/CWC (Battle Axe/Corded Ware) Steppe-related cultures. a) Distribution during the Scandinavian Early Neolithic to Middle Neolithic A periods, relevant contemporaneous sites with comparative data are shown in black. The dashed grey line shows the northern limit of the LBK (Linearbandkeramik) culture. Cultural distributions are shown in green (TRB), red (PWC), and blue (GAC/Zlota cultures). Insert: Map of Gotland showing the location of TRB-sites (squares) and PWC-sites (pyramids). The Ansarve dolmen from this study is shown in red, sites with comparative data are labeled. b) Distribution during the Scandinavian Middle Neolithic B period. The PWC and GAC/Zlota cultures show the same distribution and color as in a. Distribution of BAC/CWC (Battle Axe/ Corded Ware) cultures are shown in yellow, contemporaneous sites with comparative data are shown in black. Cultural timelines in both a and b are presented as cal BCE, see Extended Dataset 1 for comparative data.

The Scandinavian peninsula was initially populated from the south by Mesolithic HG-groups displaying genetic ancestry from western HGs (WHGs)^8^, but also from northeast by HG-groups exhibiting an eastern genetic ancestry (EHG) resulting in varying degrees of WHG and EHG-admixture into the regional Scandinavian HG-groups (SHGs), which is also seen on Gotland^14,15^. The Neolithization in southern Scandinavia (including Gotland), is associated with the Funnel Beaker/Trichterbecher cultural complex (TRB, 4000 – 2800 calBCE)^e.g. 16–22^. The transition is associated with the appearance of groups exhibiting genetic ancestry mainly from the ENF, but also exhibiting some degrees of ENF and HG-admixture^2,5,7,8,11,23,24^. It has recently been shown that TRB-associated individuals from present-day Denmark, Sweden, and Poland form a genetic subcluster, suggesting a northeastern European origin for the source population of the Scandinavian TRB groups^8^. Around c. 3600 calBCE^25^ a new megalithic burial tradition was manifested in the TRB-complex in form of dolmens and passage-graves^21,26–29^, indicating continued long-range contacts within the widespread TRB-sphere [Fig. 1A].

It has been suggested that the forager admixture components in the Scandinavian TRB-groups was WHG-related and that admixture mainly occurred prior to reaching Scandinavia^8^, though local admixture with SHG-related groups has also been suggested^5^. However, Neolithic period forager groups were also present in much of Scandinavia, i.e. the marine Pitted Ware culture (PWC, 3300-2300 calBCE), that exhibit contacts with local TRB-groups making it possible that there were direct interactions between these contemporaneous groups. The PWC thrived along the coastal areas of present-day northeastern Denmark up to middle Sweden, southern Norway, on Gotland, and western islands of the Baltic Sea [Fig. 1A]^19,30–33^. While the culture was widespread, genetic analyses have previously only been conducted for Gotlandic PWC-associated individuals, revealing mainly SHG-ancestry with a slight degree of ENF-admixture^e.g. 10^. In present-day Denmark, two individuals contemporaneous with the Gotlandic PWC have been found that display only SHG-ancestry (and no ENF-admixture)^8,34^, suggesting differences in the demographic history.

PWC-associated human remains are rare in the archaeological record of mainland Scandinavia. However, the remains of both TRB- and PWC-associated individuals are represented in the archaeological record on the island of Gotland. Until recently, it was suggested that Gotland in the Middle Neolithic was settled by one group with seasonal subsistence practices that adopted farming and animal husbandry^32,35^. However, archaeogenetic studies have shown that individuals from the two archaeological groups -- associated with different subsistence economy and cultural practices -- exhibit distinct genetic variability, and thus, different demographic history and ancestry.^11,24^. Although they coexisted on Gotland for over 500 years^12^ [Supplementary section 1], the social contacts and, thus, possible gene-flow between the TRB and PWC groups on the island, have not been fully investigated.

While at least ten sites with TRB pottery and domestic faunal remains have been found on Gotland^36,37^, only one confirmed megalithic TRB burial has been located to date, the Ansarve dolmen [Fig.1]^12,38–41^. Previous archaeogenetic analyses of individuals buried in the Ansarve dolmen revealed continuity in the male lineage over time, as well as a 2^nd^ degree kinship relation between two males suggesting that the burial was used by a patrilocal society^24^. These results are in accordance with the recent findings for the contemporaneous TRB on the Swedish mainland^7^ and emphasize the uniqueness of the Ansarve burial for the local TRB groups on the island.

At least twenty different PWC-settlement sites have been documented along the former Gotlandic coastline [Fig. 1] and several of them include burial grounds with many flat grave burials^32,35,42–47^. New sites are still being found^48^ which indicates that the PWC culture was abundant. Thus, the TRB and PWC complexes were contemporaneous on Gotland during Scandinavian Middle Neolithic (MN) period between c. 3300 – 2800 calBCE, but after ∼2700 calBCE only the PWC complex remained.

Around c. 2800 calBCE the archaeological record changes in present-day Sweden with the appearance of the Battle Axe (BAC/CWC) cultural complex^19,44,49–53^ [Fig. 1B]. Archaeogenetic research show that the transition coincides with the dispersal of humans with a Steppe-related ancestry component^3,6–8^. The BAC/CWC-phase overlaps with the end of the TRB phase on Gotland. Some BAC stray artefacts have been found across the island^38^, and some artefact types even in some PWC-burials^43,44,54^, although, no typical BAC-burials or settlements have been located. Neither has any Steppe-related admixture been detected in individuals of PWC burials, not even in those with cultural BAC influences^10^.

Interestingly, during this period of high human mobility, interaction, and cultural and social transformation in the region, pathogens, and more specifically *Yersinia pestis,* seem to have widely spread, and several prehistoric *Y. pestis* strains have recently been reported. The earliest basal lineage to date was found in a hunter-fisher-gatherer from Riņņukalns, Latvia (3350 – 3100 calBCE)^55^, prior to the appearance of CWC in the Baltic area. However, repeated outbreaks of plague have been discovered in a TRB community in Falbygden and several other sites across southern Scandinavia, showing that the so called Neolithic plague was common in these regions 5000 years ago^7,56^. Additional lineages have been discovered in two contemporaneous individuals (3340 – 2900 calBCE) from the Warburg Necropolis in western central Germany, of which one lineage seem to fit within the Falbygden *pestis* clade^57^. A few centuries later, and concomitant to the CWC migrations, additional strains from the so-called Late Neolithic/Bronze Age (LNBA) *pestis*-lineage arrived in the Baltic area, found in two individuals from Gyvakarai, Lithuania (2621 – 2472 calBCE) and Kunila, Estonia (2574 – 2340 calBCE), both exhibiting the Steppe-related ancestry component^58^. Thus, it has been shown that this pathogen probably played an important role during the Neolithic decline^7,56^ and its potential presence on Gotland could add new information to the demographic developments on the island during the Scandinavian Neolithic.

In this study we reanalyze the six individuals from the Ansarve dolmen^24^, together with 19 individuals from three PWC sites on the island (Ajvide, Hemmor, and Västerbjers) presented in Coutinho et al.^10^ to elucidate the demographic development on Gotland when the TRB- and the PWC complexes co-existed. We present new high coverage genomic data for one of the Ansarve individuals, as well as new low coverage data for a later dated individual buried in the dolmen. Finally, we investigate the possible presence of ancient strains of *Y. pestis* in the Gotlandic individuals.

## Results

From here on we use the collective label “Neolithic Farmer” (NF) when referring to all archaeological individuals/groups from an “Early – Middle” Neolithic context in the population genetic analyses, as genetically they broadly shared the same ENF-demographic history. For more information on reference panels and labels see Fig. 2A, Table 1, Extended Dataset 1, and Supplementary section 4.1.

**Figure 2.**
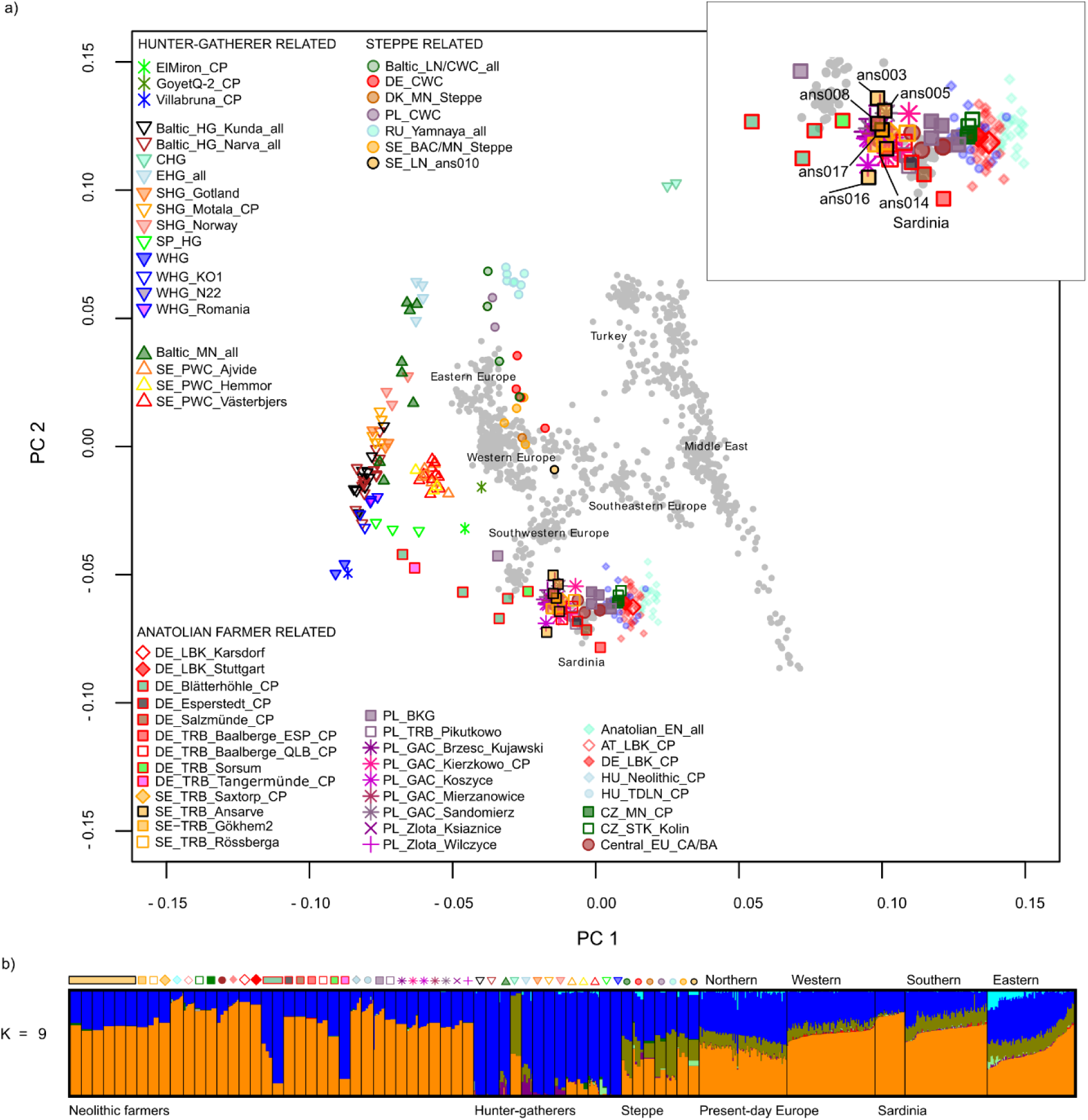
PCA and unsupervised admixture plot. a) PCA of 991 present-day individuals (gray dots) from 67 European, Near Eastern, and Caucasian populations from the Human Origins v2 data, with previously published HGs (top left labels), Neolithic Farmer groups (bottom left labels), and Yamnaya/BAC/CWC culture associated individuals (top right labels) projected onto the first two principal components. The nomenclature is based on a two-letter International Organization for Standardization (ISO) code for country, plus abbreviated cultural name/ or the time-period(s), and site name (for groups)/ or lab names (for individuals). The Scandinavian data is based on whole genome shotgun (SG) sequencing, comparative capture-generated data has an added “CP” tag, and the tag “all” is used when both are combined. [see section S5.1 and Extended Dataset 1 for more information on the cultures and nomenclature]. b) Unsupervised ADMIXTURE modeling results (K = 9) for ancient samples (with > 0.05x genome coverage, except SE_LN_ans010) [Extended Dataset 1], and 2627 individuals from 320 world-wide populations from the HO dataset v2 [all Ks are shown in Extended Data Fig. 1]. The dark blue component is maximized in WHG, the orange component is maximized in Anatolian NFs, the green component is maximized in CHG and Yamnaya. The plum and cyan component is related to Ancient North Eurasian ancestry which is present in EHG and Steppe-related individuals. Labels correspond to [Fig. 2A].

**Table 1.**
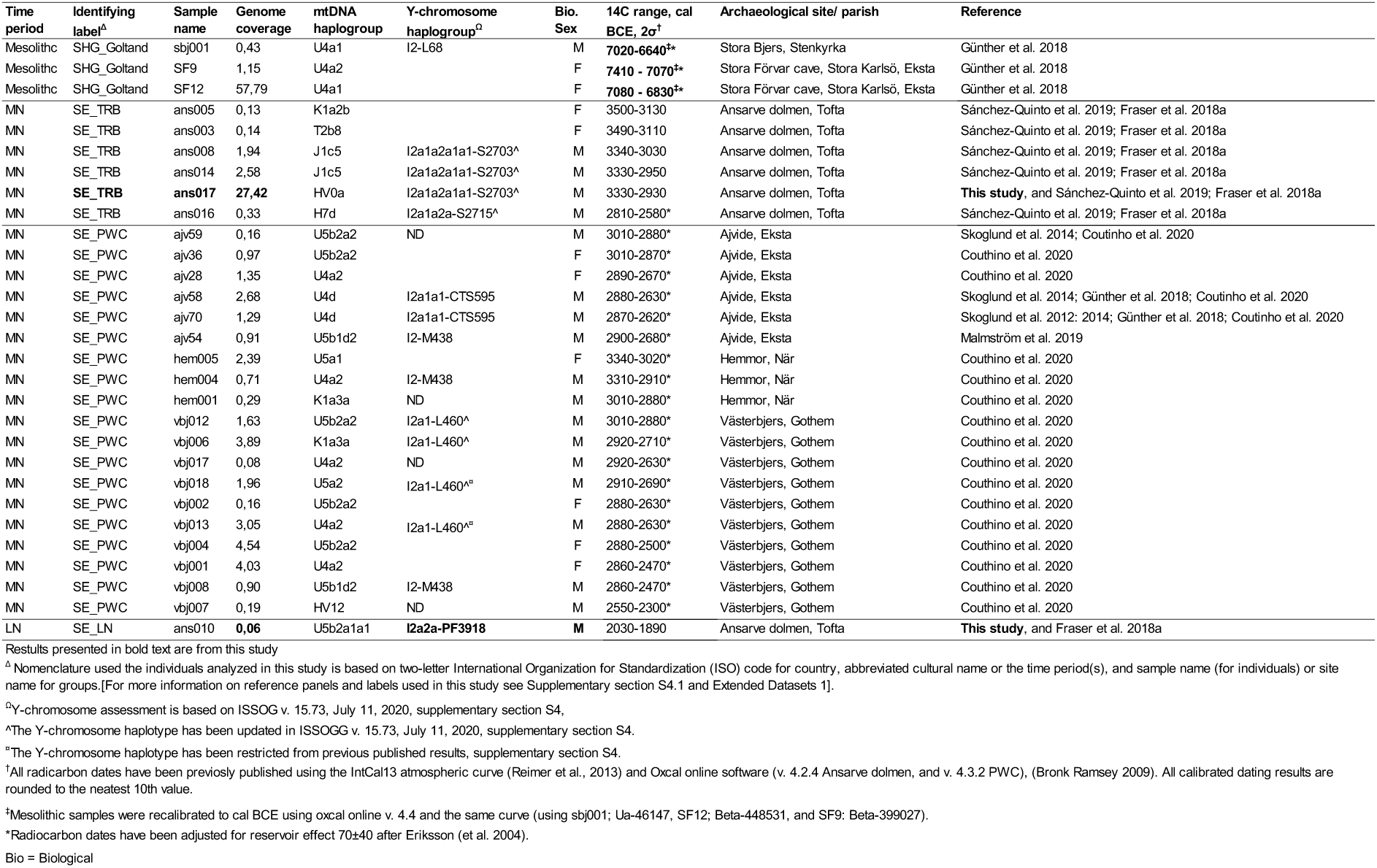
Ancient individuals analyzed from Gotland in this study, showing nomenclature used in this study, genome coverage, mtDNA and Y-chromosome haplogroups, biological sex, 14C range, archaeological site and parish, plus reference to data.

### Genome sequencing and population structure

We generated a 27.42x coverage Uracil-DNA glycosylase (UDG) treated genome for the juvenile male SE_TRB_ans017 (3330 - 2930 calBCE) presented in Sánchez-Quinto et al.^24^. As well as, 0.06x genome coverage for the adult individual SE_LN_ans010 (2030 - 1890 calBCE) from later use of the burial presented in Fraser et al.^12,13^, which we here could determine to be male [Methods, Table 1 and Supplementary Tables S1 and S4]. The sequence data exhibit characteristic properties of ancient DNA: short fragment size, and cytosine deamination at the ends of fragments^59^ between 0.15 - 0.58, and contamination levels were found to be low [Supplementary Fig. 1 and Supplementary Tables S2 – S3].

We projected the Ansarve and the PWC individuals together with 210 published individuals with ancient genome-wide data, dated between the Upper Paleolithic and Early Bronze Age [Extended Dataset 1], on top of the two first principal components using 991 present-day individuals from 67 European, Near Eastern and Caucasian populations from the Human Origin v2 data^60,61^ [Methods]. Specifically, we compared the Gotlandic individuals to individuals from contemporaneous central European cultural groups and find that the Scandinavian TRB cluster together with individuals from the Globular amphorae (GAC) and Zlota cultural contexts in Poland in the PCA space, sharing similar levels of HG admixture. Some contemporaneous NF individuals with elevated HG-admixture (e.g. Blätterhöhle, Tangermünde, and Sorsum) form a cline between the Scandinavian NF cluster and WHG [Fig. 2A and 2B]. As previously shown^10^, the PWC individuals form a separate cluster separated from SHGs slightly in the direction of NFs caused by low levels of NF-admixture [Fig2B]. Furthermore, the LN male individual from the Ansarve dolmen (SE_LN_ans010, 2030-1890 calBCE) fall in a cline together with other individuals that display Steppe-related ancestry, making him the earliest individual with Steppe-related ancestry found on Gotland to date.

Further exploring the genetic structure among the NFs, we observed that the NF-individuals broadly separate into three main clusters of different geographic dispersal: i) Scandinavian, ii) Polish, and iii) EN – MN Central European and Anatolian cluster [Fig. 3A], based on a shared-drift *f3-statistic*^61^ MDS analysis [Methods]. At the group level, we confirm the clustering of the Polish and Scandinavian TRB^62^, however we also see a connection with the DE_TRB_Baalberge_QLB_CP individual [Supplementary Fig. 2]. At the individual level, we observe more genetic drift indicating some degree of isolation among the Ansarve individuals which differentiates them from Scandinavian mainland TRB contexts in Falbygden (Gökhem and Rössberga) [Fig. 3A]. The differentiation between the Ansarve individuals and TRB-associated individuals in the Falbygden area with > 250 megalithic burials can also partly be explained by the many familial relations found in that region^7^.

**Figure 3.**
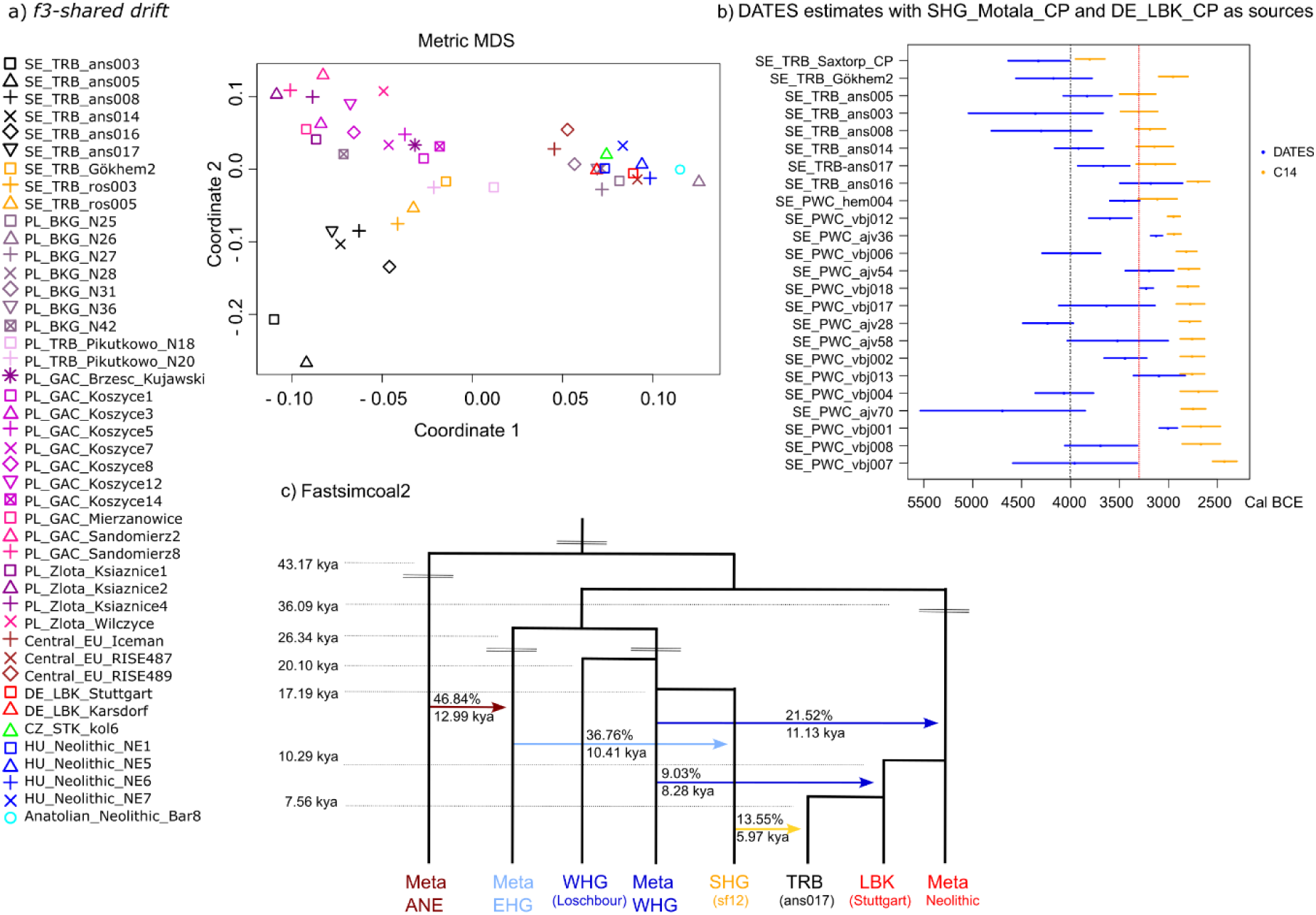
*f3*-shared drift MDS plot, DATES admixture estimation analysis, and demographic inference with *fastsimcoal2*. a) MDS plot of *f3*-shared drift between the MN and later dated individuals from a farming context. b) DATES analysis of timing of HG-, and NF-admixture in the Scandinavian TRB and PWC, respectively. Blue = point estimate admixture time in years (25 * generation) and SE, Yellow = point estimate for radiocarbon dates calBCE, 95.4% CI. Black dashed line = Neolithization of Scandinavia, Red dashed line = arrival on PWC material culture to Scandinavia and Gotland. c) Demographic inference with *fastsimcoal2 -* best-fitting model (# 4): WHG (Loschbour), SHG (sf12), ENF (Stuttgart), and SE_TRB_ans017 representing the Ansarve population. Unsampled meta populations labelled as “Meta ANE” (Ancestral North Eurasian), “Meta WHG”, “Meta EHG”, and “Meta Neolithic” with point estimates for branch split time, admixture event, and the admixture proportions [Extended Table S13].

### Social structure and dynamics in Neolithic Gotland

Previous studies demonstrated close kinship connections up to the 2^nd^ degree in the Gotlandic TRB and PWC individuals^10,24^. Here, we also noticed that the two chronologically older females (ans003 and ans005) shared more drift with each other than with the other Ansarve individuals [*f4*-*statistics*, Supplementary Fig. S3]. We ran ancIBD^63^, which can detect kinship relations up to the 7^th^ degree, and apart from the 2^nd^ degree male relationship at Ansarve^24^, we found a 3^rd^ degree relationship between the above mentioned females [Methods and Extended Table S8]. However, there was not sufficient data to determine the nature of their relatedness, and the coverage for both individuals [Table 1] was below the recommended 0.4x cut-off value.

We found kinship relations also in the three PWC groups and confirm the 2^nd^ degree relatedness between SE_PWC_vbj001 and SE_PWC_vbj008^10^, here predicted to be grandparent-grandchild, and found another 3^rd^ degree (great grandparent-grandchild) relation between SE_PWC_vbj006 and SE_PWC_vbj013. As well as several other kin relations from the 2^nd^ to the 3-4^th^ degree, both within and between all three sites, which suggest a long population continuity for PWC on the island and also contacts between the different sites [Extended Table S8].

We applied several different methods to understand the past population sizes of the different Stone Age groups [Methods and Supplementary sections 5.8 – 5.10]. From a PSMC analysis^64^ of the high-coverage genomes of SE_TRB_ans017, SHG_Gotland_sf12, WHG_Loschbour, and DE_LBK_Stuttgart, we found that the Ansarve TRB had a greater effective population size (Ne) compared to SHGs and WHGs, but smaller compared to ENFs, [Supplementary Fig. S11]. These observations are largely consistent with runs of homozygosity (ROHs)^65^ analyses of imputed data (from >∼1x genomes) [Methods and Supplementary Fig. S7], where the NF-individuals displayed broadly the same proportion of short ROH tracks along the genome with some slight variation on the amount of long ROHs.

The Mesolithic and Neolithic HGs, on the other hand, showed a higher proportion of short (< 1.6 Mb) and long (> 1.6 Mb) ROH tracks than the NF-groups [Supplementary Fig. S7], which implies that the HG-populations lived in smaller groups. The PWC individuals have a smaller proportion of these tracks than the Mesolithic HGs, probably as a consequence of the NF gene-flow. The lower Ne in the Gotland PWC individuals compared to the NFs was further confirmed from analyzing tracks of homozygosity (> 4cM)^66^ [Supplementary Fig. 10], and from conditional nucleotide diversity estimation^5^ [Supplementary Fig. S6].

However, the individual SE_TRB_ans017 showed a greater fraction of long ROH tracks (mostly above 12cM and longer) that also were present in most of the chromosomes [Supplementary Figs. S8 – S9], which is an indication of inbreeding. He was also predicted to be the offspring of closely related parents^65,66^ [Supplementary Fig. S10]. This observation is in accordance with the inbreeding coefficient estimation (F = 0.10) for SE_TRB_ans017 [Methods].

In order to assess the biological impact of the inbreeding in SE_TRB_ans017, we screened the high-coverage genome for deleterious variants [Methods]. Six variants were classified as “high impact” using SNPEff^67^ [Supplementary Table S11], however most of these variants either occurred at previously unknown variable sites, were present as heterozygotes, or its functional consequences are unknown which complicates the interpretation of their biological implications [Supplementary Table S12].

### Admixture patterns analyses

As the Ansarve individuals display some level of gene-flow with HGs, and PWC individuals display some level of gene-flow with NFs, and the two groups coexisted on Gotland for more than 500 years, we investigated if this admixture could have occurred during the period of coexistence on the island. It has previously been shown that NF admixed with WHGs during the continental expansion^4,5,68^. Thus, the ancestors of the Ansarve individuals exhibited evidence of WHG admixture prior to arrival to Scandinavia^8^. However, other studies have also proposed a later admixture between Scandinavian NF and SHG^e.g. 5^.

We used qpGraph^61^ to model different scenarios. As we wanted to investigate HG admixture over time in the Ansarve TRB, we performed the analyses at the individual level for each Ansarve individual but investigated the PWC as one group. First, we tested a scenario with WHG, Anatolian_Neolithic and SHG_Motala admixture into both Ansarve TRB individuals and into the PWC group, but without admixture between them [Methods and Supplementary Fig. S4]. Four out of the six Ansarve individuals (ans005, ans008, ans017 and ans016) are compatible with that model. Furthermore, these results also indicate that the Ansarve individuals exhibit between 9 - 21% gene-flow from a SHG-related population in addition to their Farmer/WHG admixed ancestry. Interestingly, this SHG-admixture seems to have increased with time, when considering the radiocarbon dates of these individuals. The Ansarve individuals (ans003 and ans014) displayed less SHG-admixture (7% and 2%, respectively), but they were poorly fitted to the model.

Next, we assessed if the PWC group could be modeled as a mixture between SHG and the Ansarve TRB. This model would imply gene-flow between an Ansarve TRB-related source and the ancestors of the PWC (occurring either on Gotland or the mainland). Our results indicate that PWC individuals as a group can be modeled as having derived 31-36% Ansarve ancestry [Supplementary Fig. S5], likely acting as a proxy for farmer ancestry.

We then investigated closer genetic affinity (using a cross-coalescence rate approach^64^) between Ansarve_TRB (represented by the high-coverage genome ans017) and WHG_Loschbour, SHG_Gotland_SF12, plus DE_LBK_Stuttgart and found that SE_TRB_ans017 shared more recent ancestry with a SHG-like populations, than with a WHG-like population [Methods and Supplementary Fig. S12].

Based on these results we modeled the above mentioned high-coverage genomes, together with unsampled ghost meta populations labelled as “Meta ANE” (Ancestral North Eurasian), “Meta WHG”, “Meta EHG”, and “Meta Neolithic using fastsimcoal 2.7^69^ to infer the Ansarve demographic population history [Methods]. We tested four models: (1) No pulse admixture into the Ansarve population, (2) A WHG pulse admixture into the LBK+Ansarve ancestor, (3) A SHG pulse admixture into the Ansarve, and (4) A WHG pulse admixture into the LBK+Ansarve ancestor, plus a SHG pulse admixture into the Ansarve. We found that model (4) had the best likelihood [Fig. 3C, Methods, Extended Table S13 and Supplementary Fig. S15]. The total hunter-gatherer (SHG+WHG) ancestry in SE_TRB-ans017 amounts to 38% (95% CI: 11.24 - 44.71) [Methods], consistent with the estimates inferred from qpGraph [Supplementary Figs. S4 – S5].

We then investigated the timing of HG-admixture into the Scandinavian TRB individuals [DATES^70,71^, Methods and Extended Tables S6.1 – S6.2] and show that most of these gene-flow events with foragers occurred in their distant ancestors, either from a WHG-related population in the continent prior to the Neolithization of Scandinavia, and/or in Scandinavia with a SHG-related population prior to the appearance of the PWC [Fig. 3B]. Interestingly, SE_TRB_ans016, who displays the most SHG-ancestry (21%), also shows a more recent admixture event, after the appearance of the PWC-complex on Gotland. Furthermore, although the NF-admixture events seen in most of the PWC individuals happened in their distant ancestors, after the NF dispersal to Scandinavia and prior to the development of the PWC on Gotland, several individuals (SE_PWC_ajv36, SE_PWC_ajv54, SE_PWC_vbj001, SE_PWC_vbj013, and SE_PWC_vbj018) displayed more recent NF-admixture, that could have happened on the island.

### Y. pestis detection in the Ansarve and PWC samples

Lastly, we investigated the possibility of *Y. pestis* being present on Gotland at Ansarve and the PWC-sites^2,5,10,12,14,24,68, and this study^ [Methods and Supplementary section 8], given previous reports of *Y. pestis* in Scandinavia, the Baltic, and central Europe^7,55–58^.

Out of 783 libraries from 11 Ansarve, and 19 PWC individuals, we found evidence for the possible presence of *Y. pestis* sequences in five individuals (SE_TRB_ans003, SE_TRB_ans005, SE_TRB_ans007, SE_PWC_ajv58, and SE_LN_ans010). Since the number of mapping reads was too low to be able to perform robust phylogenetic analyses, we authenticated our results by combining different strategies; (i) competitive mapping against representatives of all known species of the *Yersinia* genus, (ii) edit distance profiles, (iii) number of reads mapping on chromosomes and plasmids, and (iv) the distribution of single nucleotide variants (SNVs) among strains [Supplementary Tables S14 – S15 and Supplementary Figs. S16 and S19].

Our results supported, in all the analyses, the presence of *Y. pestis* in the LN individual (ans010; 2030 - 1890 calBCE), with a strain that most likely belonged to the LNBA-lineage defined in recent publications^58,72^. Membership to this lineage could be expected by the temporality of this sample. We also confirmed the finding of *Y. enterocolitica* previously detected in lower coverage data from this individual^73^, which could imply a co-infection with both species [Extended Table S16 and Supplementary Fig. S17 – S18]. *Y. enterocolitica* causes a possible lethal intestinal infection (yersiniosis) generally spread by infected water and food, commonly obtained from digesting raw or undercooked meat from suids^74^. A co-infection of the earlier Neolithic *Y. pestis strain* and *Y. enterocolitica* has previously been found in a MN individual from Falbygden^7^. Although we found evidence of *Y. enterocolitica* in several of the other individuals analyzed here, including the previously reported SE_PWC_ajv58^73^, there was too little data to confirm these results.

Similarly, authenticating the presence of *Y. pestis* was more challenging for the other four individuals. However, by combining the complementary evidence provided by the different authentication analyses we could verify the presence of *Y. pestis* in the early phase of the Ansarve burial (ans003; 3490 - 3110 calBCE), supported by the presence of reads mapping into the three typical plasmids of *Y. pestis* (pMT1, pPCP1 and pCD1), edit distance profiles closer to *Y. pestis* than to the closely related ancestor *Y. pseudotuberculosis,* and the presence of several SNVs that are typically characteristic of *Y. pestis* [Supplementary Table S14 and Supplementary Figs. S16 and S20]. Although the observed SNVs suggest an affinity to some of the most basal strains of *Y. pestis*, we could not determine which strain was the closest. We were not able to unambiguously validate the presence of *Y. pestis* in the other two MN individuals from the dolmen (ans005 and ans007), thus higher sequencing efforts would be necessary to fully authenticate these results.

We did however find that *Y. pestis* was most likely present in SE_PWC_ajv58 (c. 2880 - 2630) from the Ajvide PWC site, as evidenced by several SNVs that are typically present in *Y. pestis* and not, for instance, in *Y. pseudotuberculosis* [Supplementary Fig. S20]. Moreover, we also identified several SNVs that suggest a closer affinity to the most basal RV2039 strain^55^ than to any other Neolithic and LNBA strain, suggesting that this individual was infected with a strain that is closer to the one infecting HGs in the Baltic.

## Discussion

Our analysis has generated unique insights into the lives and social networks of the descendants of the first Neolithic farmers (TRB) on the island of Gotland and their relationship to the contemporaneous marine HGs (of the PWC complex). These distinct archaeological groups coexisted on the island for more than 500 years and had different demographic backgrounds, material culture expressions, subsistence economy, and lifestyles.

Regarding the genetic affinities among TRB groups, we confirm the connection between Ansarve and other TRB-associated individuals from Scandinavia and Poland^7,62^ but also find a connection with a TRB-associated individual from present day Germany, suggesting a common genetic origin among groups of the TRB cultural sphere. The presence of the Ansarve dolmen confirms that the Gotland TRB group must have had contact with the larger TRB-complex, as they erected the monumental burial at the same time as they appear in Scandinavia and the continent, and more than 500 years after the initial appearance of TRB material culture on the island.

The Ansarve individuals seem to be genetically more similar to each other than to other TRB individuals and groups, likely due to the relative isolation on Gotland [Fig. 3A and Supplementary Fig. S3]. The presence of many TRB settlement sites and the abundance of TRB-associated stray finds across the island, do however suggest that there must have been a larger TRB population on the island, also confirmed by analyses of Ne. Whether or not the Ansarve group is good representatives of this larger population remains unknown, since no other TRB-associated burial has been found on Gotland.

In addition to the previously shown male kin-relation (ans014 and ans017) we also find that the two females (ans003 and ans005) from the earliest use of the burial were likely related to the 3^rd^ degree. These females could possibly be cousins, or great aunt/niece, although a strict matrilineal kinship relation can be excluded based on their mitochondrial haplotypes [Table 1]. Interestingly, even though the contemporaneous males (ans008 and ans014) share both Y chromosomal and mtDNA haplogroup lineages, they are not close kin related. However, as there were ∼30 individuals in the burial, and at least 15 had previously been dated to the MN period^12^ there most likely were additional kin relations, as seen in the contemporaneous megalithic burials in mainland Sweden^7^. As previously shown^65,66^, SE_TRB_ans017 was predicted to be the offspring of 1^st^ degree cousins, and showed signs of inbreeding (F = 0.10). Although rare, a few cases of inbreeding have also been found at other Neolithic farmer sites^7,65,75^. Thus, the kinship relations and close parental relatedness suggest that the Ansarve dolmen was the burial site of an extended family with long continuation over time.

We confirm previous reported kin-relations between PWC individuals^10^, as well as additional kin-relations, both within and between the three sites (Ajvide, Hemmor, and Västerbjers), suggesting close contacts between the different PWC groups in different regions of the island. We do, observe a slightly higher Ne in the earlier individuals from Hemmor, compared to Ajvide and Västerbjers, which may be an indication of bottleneck after the initial settlement, and subsequent isolation of the PWC population. This bottleneck does however not suggest elevated inbreeding in the later PWC individuals.

Contrary to previous suggestion of no local geneflow from SHG into the Scandinavian TRB^8^, we do find evidence of SHG–related gene flow in the Ansarve individuals. They can be modeled with an additional 9 – 21% SHG admixture, in addition to their already NF/WHG admixed ancestry [Supplementary Fig. S4]. We find that these gene flow events mainly happened prior to the appearance of the PWC material culture complex [Fig. 3B], with an SHG-related population either on the Swedish mainland, or possibly on Gotland as there is some Late Mesolithic material culture evidence on the island^32^.

We found evidence of more recent HG gene flow in the latest dated male Ansarve TRB-individual (ans016; 2810 – 2580 calBCE). As he displays non-local childhood Sr-signals^12^ this gene flow event probably happened outside of Gotland. Moreover, recent HG admixture does not seem to have been common within the Scandinavian TRB, as only two females (FRA108 and ROS027) out of 105 investigated individuals from Falbygden in Västergötland, were suggested to have recent HG-admixture^7^. Furthermore, these individuals, from three different megalithic tombs, date to the end of the TRB cultural phase which, as previously suggested for Västergötland^7^, could indicate relaxed socio-cultural boundaries and/or a demographic decline, and that interactions between the TRB and PWC groups had become more common at this time.

Similarly, the majority of NF-admixture seen in the Gotlandic PWC individuals appears to also have happened after the initial dispersal of Neolithic groups to Scandinavia but prior to their appearance on Gotland. We do, however, find evidence of recent NF-admixture in several individuals from Ajvide and Västerbjers, which suggests ongoing gene flow occurring between TRB and PWC groups. We show that the PWC (as a group), in addition to their SHG-ancestry, exhibit ∼30% of NF/WHG admixed ancestry which can be modeled as gene flow from an Ansarve-related population. However, the exact admixture proportions are difficult to assess as the individuals from both groups already were admixed in different ways.

We did not detect any distant kin-relations between individuals from the two archaeological groups. The archaeological record on Gotland show that some long-range trade occurred, indicating contacts between the PWC and TRB. However, there are no obvious indications of material culture hybridity of the two cultural complexes, as seen in the pottery styles at e.g. the Fagervik PWC site and at the Alvastra Pile dwelling on the Swedish mainland^e.g. 19^. Vanhanen et al.^33^ proposed that some mainland PWC-groups adopted cultivation from TRB farmers. However, on Gotland, it is rather in later dated PWC contexts, and after the TRB cultural phase had ended, that new agricultural elements such as grindstones and skeletal remains of domesticated animals are found^42,76^. Further analyses of additional PWC burials at the end of the TRB phase on Gotland are needed to clarify the level of NF gene-flow into the PWC.

Gotland’s placement in the center of the Baltic Sea and pre-historic maritime customs increased the chance of contracting disease from different geographical areas. Here we were able to detect evidence of *Yersinia pestis* in three individuals from different time periods and different cultural affinities. The early TRB female (ans003) carried a lineage which, although not specified further, suggest affinity to a basal *Y. pestis* strain. As seen in Falbygden^7^, the Neolithic plague was widespread during this time.

We also report an infection by a LNBA-strain for the first time in Scandinavia. The Late Neolithic male from the dolmen (ans010; 2030 – 1890 calBCE) was most likely infected with the LNBA-strain from the continent. He also had a coinfection of *Y. enterocolitica*^73 and this study^. Unfortunately, the cultural affinity of this individual is not known as he was interred in an older burial structure, but as he showed Steppe-related ancestry, and a non-local childhood Sr-signal^12^, it could possibly be evidence of a plague transmission route. However, the individual is represented by one tooth only, and due to the low genomic coverage, of both the individual (0.06x) and the pestis genome, further analyses were not possible to conduct.

Interestingly, the third strain, found in a PWC male (ajv58) from Ajvide, suggests closer affinity to the most basal RV2039 lineage found in Latvia^55^. This could indicate contacts to the east which have previously been discussed based on similarities in PWC-artefacts and burials customs with the Late Combed Ware culture in the Baltic^19,77^. Interestingly, *Y. pestis* has also been detected in a contemporaneous dog from the same PWC site^57^. Although our results on *Y. pestis* and *Y. enterocolitica* are well-supported, further sequencing and capture-enrichment strategies will be needed to perform phylogenomic analyses that can provide a clearer picture on the strain diversity and their spread between these populations and regions.

In summary, we have examined the demographic history of two genetically and archaeologically distinct Neolithic groups on Gotland (TRB farmers and PWC marine foragers) by analyzing their genetic variation. While the Gotland TRB share ancestry with contemporaneous TRB groups from present-day Scandinavia, Poland, and Germany, they also show some level of isolation. Furthermore, although we find some evidence of recent admixture between the Ansarve TRB and the Gotlandic PWC that could have happened on Gotland, we show that most of the interaction between the different groups appears to have happened in the past, and prior to PWC appearing on the island. We also find evidence of different *Y. Pestis* strains in individuals from both TRB and PWC on the island, as well as in a Late Neolithic individual buried in the Ansarve dolmen after the TRB phase. Thus, different factors could have aided in the decline of the TRB-complex on the island. However, the extent to which pathogens such as *Y. pestis* might have affected these populations on Gotland needs further investigation.

## Methods

### Sample preparation, DNA extraction, library building, authentication, and contamination control

The ans017 and ans010 samples were prepared in facilities dedicated to analyses of ancient DNA [aDNA] at the DNA Laboratory DBW (De Badande Wännerna) at Campus Gotland, and at Human Evolution, Institute of Organismal Biology (IOB), Uppsala University (UU) as previously presented^12,24^. Illumina multiplex DNA libraries were prepared using blunt-end (BE) ligation with P5 and P7 adapters, and amplification for 9-15 cycles using IS4 and index primers from Meyer and Kircher^78^ [Supplementary section 2 and Supplementary Table 1]. Twelve additional Uracil-DNA glycosylase (UDG) treated damaged repaired Illumina multiplex BE DNA libraries were produced on three previously generated extractions^12,24^ for ans017 following Meyer and Kircher^78^ and Briggs and Heyn^79^, deep sequenced to high coverage, and merged separately. All libraries were quantified on Bioanalyzer 2100 (High Sensitivity DNA chip) or 2200 TapeStation (High sensitivity DNA screentape), Aligent, following manufacturer’s protocol and sequenced on Illumina Hiseq 2500 (v.4 chemistry and 125 bp paired-end reads) or HiSeq Xten (v2.5 chemistry and 150bp paired-end reads) at the SNP & SEQ technology platform, UU. Additional data for ans010 was generated by resequencing previously made libraries [Supplementary Table S1]. Sequencing data was processed following Günther et al.^14^ as previously reported^12,24^ [Supplementary Table S2].

Authentication and contamination was estimated from read length [Supplementary Table S2] and deamination patterns [Supplementary Fig. S1], as well as from different contamination-estimation approximations applied to three data sources: [i] the mitochondrial genome^80^, [ii] the X chromosome if the individual was male^81,82^, and [iii] the autosomes^83,84^ as described^14,24^ [Supplementary Table S3].

### Biological sexing and uniparental markers

Sex determination for the Late Neolithic ans010 was conducted using the *Rx*^85^ method [Table 1]. The mtDNA haplotypes for SE_TRB_ans017 and SE_LN_ans010 were classified using Halplogrep 3 (v.3.2.1) and coincide with previously reported findings^12,24^ [Table 1]. The Y-chromosome haplogroups (Y-chr hg’s) for all male individuals in Table 1 were classified using all single base substitutions from the International Society of Genetic Genealogy (ISOGG; http://isogg.org) version 15.73, 11 July 2020. Published Y-chr hg’s classified from earlier ISSOGG versions were updated, and the Y-chr hg’s for SE_PWC_vbj013 and SE_PWC_vbj018^10^ have been restricted due to more stringent reporting in this study [See supplementary section 3 for more information].

### Panels and labels for the population genetic analyses

Ancient DNA data was merged with various published datasets (Panels 1 - 4), depending on the nature of the analyses, by generating pseudo-haploid data [Supplementary section 4]. For each SNP site, a random read covering that site with minimum mapping and base quality of 30 was drawn (using Samtools 1.3v mpileup) and its allele was assumed to be homozygous in the ancient individual (pseudo-haploid)^14^. Non-biallelic SNPs in the ancient individuals were excluded from the data. The labels used for the ancient individuals in the demographic analyses are described in detail in Supplementary section 4.2 and Extended Dataset1.

### Principal Component Analysis

PCAs were performed using smartpca from the EIGENSOFT package, with the “numoutlieriter: 0 and “r2thresh: 0.2” parameters. For each ancient individual, a PCA was conducted together with 991 individuals from 67 European (EU), Near Eastern (NE) and Caucasian (Cau) populations [Human Origins (HO) Panel 1, Supplementary section 4.1] extracted from the Human Origins panel v2^60,61^ and 210 published ancient genomes [Extended Dataset 1] dated between the Upper Paleolithic and Early BA were plotted using Procrustes transformation^2^, using all SNPs as described in^86^. The result was plotted using an in-house R script from the vegan library.

### Unsupervised ADMIXTURE

Ancestry components were inferred using ADMIXTURE v1.3^87^ and was based on 2392 individuals from 213 world-wide populations from the Human Origins Panel 1 [Supplementary section 4.1]. We subset Yoruba individuals to represent African genetic variation. This subset of individuals was merged with the ancient individuals described in Extended Dataset1. Only transversion SNPs were used in order to avoid biases due to cytosine deamination [Fig. 2B and Supplementary section 4.3]. The results for all K’s are shown in Extended Data Fig. 1.

#### f-statistics

To investigate the genetic structure among the ancient individuals from a farming context we performed S*hared-drift f3-statistics* computation with the qp3Pop program in the ADMIXTOOLS package^61^ and using SGDP Mbuti individuals^86^ as an outgroup [Supplementary section 4.4.1], using Panel 2 (for individuals) and Panel 4 (for groups) [Supplementary section 4.1]. We then summarized information from each distance matrix on two dimensions using multidimensional scaling (MDS) with the ‘cmdscale’ function in the R ‘stats’ package (https://www.r-project.org) [Figure 3A and Supplementary Fig. S2]. *f4-statistics* were computed with the same set up using only SG sequenced genome-wide data overlapped with Panel 2 and different topologies dependent on the analysis [Supplementary section 4.4.2 and Supplementary Fig. S3].

### qpGraph

To further characterize the demographic history of Ansarve and the Gotlandic PWC, as well as the proportion of gene-flow from different sources into these populations we used qpGraph (version 6100)^61^ and overlapped genetic data from all relevant samples across all sites with Panel 4 and Mbuti as outgroup. The individuals used in the source groups are displayed in Extended Table S5. We accepted models with a |Zscore| < 3 values and excluded models with less than 15 000 SNPs (due to low power for rejection), or if models had inner zero-drift branches; however, final zero-drift branches were permitted [Supplementary section 4.5 and Supplementary Figs. S4 – S5].

### Admixture DATES estimate

To investigate the timing of the Admixture in the Scandinavian TRB and PWC we used DATES^70,71^ and genetic data overlapped with all sites from Panel 4. We used default parameters as suggested in the github default web page (https://github.com/priyamoorjani/DATES). SHG_Motala_CP and DE_LBK_CP were used as proxy populations [Extended Table S5] as we wanted to have data generated from the same methodology with a large sample size that would allow us to measure the time to admixture for both Scandinavian farmers and PWC simultaneously. We assumed 25 years per generation, and disregarded results when a negative number of generations was observed or when the estimated standard error was superior to the estimated number of generations. [Fig 3B and Extended Tables S6.1 – S6.2]. We also investigated to date the time of admixture using WHG_CP and DE_LBK_CP as proxy populations [Extended Table S5], however the results generally suggested much older dates [Extended Table S6.2].

### Conditional Nucleotide Diversity

We investigated genetic diversity for specific populations as previously described^5,68^. To avoid ascertainment bias, post-mortem damage, as well as to increase the number of sites we only used SNPs from Panel 3^68,86^ [Supplementary section 4.6 and Supplementary Fig. S6].

### ancIBD

We used ancIBD^63^ to infer kinship relatedness between 3^rd^ and 7^th^ degree. Samples were imputed using GLIMPSE^88^ following author recommendations (https://odelaneau.github.io/GLIMPSE/ docs/tutorials). Each sample was imputed separately to avoid batch effects. As a reference panel for imputation, we used the phased haplotypes from the 1000 Genomes dataset (http://ftp.1000genomes.ebi.ac.uk/vol1/ftp/release/20130502/). Imputed data and phased was used as input for ancIBD (v.0.2a; https://pypi.org/project/ancIBD). As suggested by authors, we down sampled the imputed data to 1,240,000 SNPs and screened all pairs of Ansarve and PWC individuals with at least 0.4x coverage (n = 16) using ancIBD recommended default settings, to avoid spurious results between the lower coverage individuals. Results for each pair of individuals with IBD detected are displayed in Extended Table S8. However, we did include the lower coverage individuals SE_TRB_ans003 (0.14x) and SE_TRB_ans005 (0.13x) as they shared more drift with each other in the *f-statistics* [Supplementary Fig. S3].

### Imputation for analysis of Runs of Homozygosity (ROHs)

Imputation was carried out using Beagle 4.0^89^ for the ancient genomes of 55 individuals with depth coverage of >0.7x [Extended Table S9], following previously published methods^90^ [Supplementary section 5]. Genotype likelihoods for biallelic autosomal SNPs in the 1000 Genomes phase 3 dataset (ftp.1000genomes.ebi.ac.uk/vol1/ftp/release/20130502) were called using the “UnifiedGenotyper” tool in “Genome Analysis Toolkit” (GATK) v3.5.0^91^. Non-biallelic sites with missing data, as well as genotypes probably derived from post-mortem damage, were excluded. Samples were merged by chromosome and imputed in 15,000 marker windows using the 1000 Genomes phase 3 haplotypic reference panel and genetic map files provided by the BEAGLE website (http://bochet.gcc.biostat.washington.edu/ beagle). To assess accuracy, imputed genotypes for downsampled 1x genomes from high-coverage samples were compared to their diploid genotype calls [Supplementary Fig. S13].

### Runs of Homozygosity (ROHs)

To further understand the demographic history of the Scandinavian farmers, in contrast to reference HGs and farmers, we estimated the ROHs from these individuals using the imputation data described above. Following recommendations from Cassidy et al. ^65^ we estimated the fraction of the total genome under ROH for both track length categories, and visually displayed the differences for each individual between each category of ROH [Supplementary section 4.7.1 and Supplementary Fig. S7].

Inbreeding coefficient was calculated following recommendations fromGazal et al.^92^ by estimating the fraction of the genome that is within homozygous-by-descent (HBD) in the high-coverage diploid data of SE_TRB_ans017 using ROH as a proxy^65^.

### HapROH

We also investigated ROHs in low coverage data (> 0.3x) from using hapROH^66^ version 0.1a4 (https://pypi.org/project/hapROH) where we overlapped pseudo-haploid data on the 1240K SNPs (Panel 4). We used default parameters and default genetic map, and 5008 haplotypes from 1000 Genomes^86^ as reference [Supplementary section 4.7.2, Supplementary Figs. S8 – S9, and Extended Tables S7.1 – S7.2].

### Multiple Sequentially Markovian Coalescent method

Effective population size per individual as a function of time “PSMC” was inferred using the MSMC method^64^ v. 0.1.0 [Supplementary section 4.8.1]. Analysis was performed on a dataset consisting of four present-day individuals from the HGDP (French, Han, Yoruba and Karitiana)^93^ together with UDG-treated high-coverage data from this study (SE_TRB_ans017) plus four previously published individuals (SHG_Gotland_sf12, WHG_Loschbour, DE_LBK_Stuttgart, and RU_Ust’_Ishim), following the workflow reported in Günther et al.^14^ assuming a generation time of 30 years and a mutation rate of 1.25×10e-8 [Supplementary Fig. S11].

To investigate the divergence over time between SE_TRB_ans017 and WHG_Loschbour, SHG_Gotland_SF12, DE_LBK_Stuttgart, respectively, we used relative Cross Coalescence rate (rCCR) and ran MSMC for two individuals (four haplotypes) [Supplementary section 4.8.2]. We restricted this analysis to only UDG-treated genomes or down-scaled the first/last ten bases (when using non-UDG-treated libraries) to avoid potential biases from non-corrected deaminations. We considered a recombination rate of 0.88, a mutation rate of 1.25e-8 and a generation time of 30 years. To calculate the relative Cross Coalescence Rate (rCCR) we divided the cross-population coalescence rate by the mean of within-populations coalescence rate, which displayed how close the populations were at a given time point [Supplementary Fig. S12].

### Discovery and annotation of novel variants in the SE_TRB_ans017

We used GATK, v3.5.0 to discover novel variants in the high coverage genome of SE_TRB_ans017 [Supplementary section 6]. Indel realignment was performed using “RealignerTargetCreator” and “IndelRealigner” GATK’s modules. Genotype calling was conducted using GATK’s “UnifiedGenotyper” with expressions “-stand_call_conf 50.0 & -stand_emit_conf 50.0 & -mbq 30 & - contamination 0.02 & --output_mode EMIT_ALL_SITES”. dbSNP version 142 was used as known SNPs with “--dbsnp” option. Variants were filtered following the detailed description in Supplementary section 6. Deleteriousness of the novel variants was evaluated using SIFT [Ng 2001] and Polyphen2^94^ [Supplementary Table S12].

### Fastsimcoal modelling

Demographic inference of the Ansarve population’s history was conducted using fastsimcoal2.7^69^ in accordance with the methods outlined by Marchi et al.^95^ [Supplementary section 7]. This analysis was performed on a panel of high-coverage genomes of ancient western Eurasian individuals, with WHG_Loschbour representing WHG, SHG_Gotland_sf12 representing SHG, DE_LBK_Stuttgart representing ENF, and SE_TRB_ans017 representing the Ansarve population.

The dataset was filtered to exclude any sites with missing data, sites where the reference alleles differ between chimpanzee and gorilla reference genomes, CpG sites or regions, and sites located in genomic regions with a recombination rate below 1 cM/Mb. To focus on neutral sites suitable for demographic inference, only BGC-free A>T and G>C polymorphic sites were retained, resulting to a final panel consisting of 112,366,543 sites.

Four models were tested: (1) no pulse admixture into the Ansarve population, (2) WHG pulse admixture into LBK + the ancestors of Ansarve, (3) SHG pulse admixture into Ansarve, and (4) WHG pulse admixture into LBK+ the ancestors of Ansarve, plus SHG pulse admixture into Ansarve [Extended Table S13 and Supplementary Fig. S15 A]. Unsampled meta populations labelled as “Meta ANE” (Ancestral North Eurasian), “Meta WHG”, “Meta EHG”, and “Meta Neolithic” were also incorporated into the models. For each model, the maximum likelihood (ML) parameters were recorded based on the run with the highest likelihood from the 50 independent runs [Supplementary Fig. S15 B]. To obtain confidence intervals around the ML parameter estimates for the best-fitting model (WHG and SHG pulse admixture into Ansarve), a parametric bootstrap approach was employed. The 95% confidence intervals were determined by calculating the 2.5% and 97.5% quantiles of the parameter distribution across the 100 re-estimated ML parameters. We estimated the total HG ancestry (SHG + WHG) in SE_TRB_ans017 [Supplementary section 7] based on the model with the highest likelihood [Fig 3C and Extended Table S13].

### Metagenomic analysis

We used MALT^96^ to screen the metagenomic data from the Ansarve and PWC individuals^2,5,10,12,14,24,68, and this study^ for microbial sequences and MEGAN^97^ tool to process MALT’s output to create a species absolute abundance table. Our initial analysis showed *Y. pestis* signals in ans003, ans005, ans007, ans010, and ajv58. We then mapped metagenomic reads to relevant reference genomes using bwa, and processed and authenticated endogenous sequences following recommendations by Key et al.^98^ as described in Supplementary section 8. A heatmap plot of shared derived single nucleotide variants (SNVs) between mapped reads from the samples of this study and 79 ancient and modern *Y.pestis* sequence was generated with R using package heatmap2^99^ [Supplementary Fig. S20].

## Data availability

the aligned BAM files are available through the European Nucleotide Archive under accession number XXX. Any other relevant data is available from the corresponding authors upon reasonable request.

## Acknowledgements

We thank Leena Drenzel at the National History Museum of Sweden for supplying the materials, Arielle Munthers for bioinformatic processing and Hanna Edlund for laboratory support. The additional data for ans010 were conducted by SciLife aDNA Unit, IOB, UU. Sequencing was performed at SciLifeLab’s National Genomics Infrastructure (NGI) Uppsala. The computations and data handling were enabled by resources provided by the National Academic Infrastructure for Supercomputing in Sweden (NAISS), partially funded by the Swedish Research Council through grant agreement no. 2022-06725.

This work was supported by Riksbankens Jubileumsfond grant no. P19.0740:1 (MF), and grant no. P21-0266 (HM), Swedish Research Council grant no. 2017-02503 (MF, HM), grant no. 2020-04789 (M.L.), and grant no. 2022-04642 (M.J.), and the Knut and Alice Wallenberg Foundation (M.J.).

## Author information

Magdalena Fraser and Federico Sanchez-Quinto: These authors contributed equally to this work.

## Contributions

M.F., F.S-Q., M.J, A.G., and P.W. conceived the study. M.F. performed the ancient DNA lab work and screening, with additional library prep performed by SciLife Lab aDNA Unit. F.S-Q. performed downstream bioinformatics and population genetic analyses, with additional help by A.B., E. A-D., M.E-R., I. A-O., and K.L-G. The metagenomic analyses were performed by N.R. and E.K., with additional help by A. L.M., and N.B. M.L. performed qpGraph and Serialsimcole2 analyses. Imputation was performed by K.M., and G.M.K. performed the (discovery and annotation of novel variants) analyses. M.F., F.S-Q., and N.R. wrote the paper with additional help by M.J., H.M., J.S., P.W., E.K., and K.K.

