## Supplementary material for "Ancestry, admixture, and pathogens in contemporaneous Neolithic farmers and foragers on the Island of Gotland"

Magdalena Fraser[a, *, #], Federico Sanchez-Quinto[a, b, *, #], Emrah Kırdök[c], Kristiina Ausmees[a, d], Gulsah Merve Kılınç[e], Maximilian Larena[a], Leonardo Correa-Mendoza[b, f], Adrien Le Meur[g], Antonio Blanchet[b, f], Nora Bergfeldt[h, i, j], Eduardo Arrieta-Donato[b, f], Mariana Escobar-Rodríguez[g], Anders Götherström[h, k], Karla Lozano-Gonzalez[l], Israel Aguilar-Ordoñez[m], Helena Malmström[a], Kjel Knutsson[n], Paul Wallin[o], Nicolas Rascovan[g], Jan Storå[p], and Mattias Jakobsson[a, #]

[a]*Human Evolution, Department of Organismal Biology, Uppsala University, SE-752 36 Uppsala, Sweden*
[b]*International Laboratory for Human Genome Research, Universidad Nacional Autónoma de México (UNAM)*
[c]*Mersin University, Faculty of Science, Department of Biotechnology, 33100 Yenişehir Mersin*, Turkey
[d]*Department of Information Technology, Uppsala University, Uppsala 751 05, Sweden*
[e]*Department of Bioinformatics, Graduate School of Health Sciences, Hacettepe University, 06100, Ankara, Turkey*
[f]*Center for Genomic Sciences, National Autonomous University of Mexico, 62209 Cuernavaca, Morelos, Mexico*
[g]*Microbial Paleogenomics Unit, Institut Pasteur, Université de Paris Cité, CNRS UMR 2000, F-75015 Paris, France*
[h]*Centre for Palaeogenetics, Stockholm University, 106 91 Stockholm, Sweden*
[i]*Department of Zoology, Stockholm University, 106 91 Stockholm, Sweden*
[j]*Department of Bioinformatics and Genetics, Swedish Museum of Natural History, 104 05 Stockholm, Sweden*
[k]*Department of Archaeology and Classical Studies, Stockholm University, 106 91 Stockholm, Sweden*
[l] *Centre for Developmental Neurobiology, King's College London*
[m]*Tecnologico de Monterrey, oriGen Project. Monterrey, Nuevo León, 64849 México.*
[n]*Department of Archaeology, Ancient History and Conservation, Uppsala University, Box 626, 751 26 Uppsala, Sweden*
[o]*Department of Archaeology, Ancient History and Conservation, Uppsala University-Campus Gotland, SE-621 67 Visby, Sweden*
[p]*Osteoarchaeological Research Laboratory, Department of Archaeology and Classical Studies, Stockholm University, SE-106 91 Stockholm, Sweden*

*These authors contributed equally to this work
#Corresponding authors

#### Data Availability

Merged BAM files produced for this study have been deposited in the European Nucleotide Archive (ENA) under accession number ENA: XXXXXX

### 1 Gotland archaeological background

#### *1.1 Mesolithic activity on Gotland (c. 7200-4000 cal BCE)*

Gotland was first populated during the Scandinavian Late Middle Mesolithic time period (c. 7200 cal BCE) and a few burials have been found from this time; four on the main island (Stora Bjers and Kambs), and also from the Stora Förvar cave on the smaller Stora Karlsö island outside the west coast of Gotland (e.g. Apel et al., 2018, 2015; Arwidsson, 1979, 1949; Lindqvist and Possnert, 1999; Lithberg, 1914; Österholm, 1989; Pira, 1926; Schnittger, 1913; Schnittger and Rydh, 1940). Recent genomic analyses from Stora Bjers (sbj001) and Stora Förvar (SF12 and SF9) have revealed that these first inhabitants on Gotland, although chronologically older, shared ancestry with other Scandinavian HGs (SHGs) (Günther et al., 2018), and showed admixture with both so called Western Hunter-Gatherers (WHGs), and Eastern Hunter-Gatherers (EHGs). However, at present it is not known whether these populations continued occupation on the island as there are few finds of archaeological remains and radiocarbon dated sites from around 5500-4000 cal BCE (Apel et al., 2018). During the same time period, the southern Baltic Sea area was inhabited by different hunter-gatherer (HG) groups such as the Ertebølle and Comb Ceramic (CCC) cultures (e.g. Fisher and Kristiansen, 2002; Hallgren, 2008; Kriiska, 2003; Price, 2015, 2000). However, some Late Mesolithic (LM) activity is found on Gotland from c. 4500 cal BCE (Apel et al., 2018; Österholm, 1989) but since there are no burials or pottery associated with this time period, the duration, cultural, and genetic affiliation of these LM forager groups on Gotland remains unknown.

#### *1.2 Scandinavian Neolithic period (4000-1750 cal BCE)*

The Neolithic period is a multicultural time in Scandinavia. The Funnel Beaker culture (TRB from German, 4000-2700 cal BCE) are associated with the Neolithization of Scandinavia and the TRB culture complex expanded from the Netherlands to Poland, and from the Czech Republic to southern Scandinavia [Fig. 1A] (e.g. Bakker, 1979; Fisher and Kristiansen, 2002; Hallgren, 2008; Malmer, 2002; Midgley, 1992; Müller, 2011; Persson, 1999; Sjögren, 2003; Sørensen, 2014). TRB cultural remains have been found reaching up to middle central Sweden and southern Norway, including the islands Bornholm, Öland, and Gotland (Hallgren, 2008; Persson, 1999; Sørensen and Karg, 2014) [Fig. 1A]. Around c. 3600 calBCE (Schultz Paulsson, 2017) a new burial tradition was manifested in the TRB-complex in the form of thousands of megalithic tombs which also have been found in southern Scandinavia (Midgley, 2008; Müller, 2011; Sjögren, 2003; Tilley, 1999).

The sub-Neolithic marine hunter-gatherers of the Pitted Ware culture (PWC, 3400-2300 cal BCE) thrived along the Scandinavian coastal areas and western islands of the Baltic Sea including Gotland and Åland [Fig. 1A] (e.g. Iversen, 2016a; Iversen et al., 2021; Larsson, 2009; Malmer, 2002; Österholm, 1989; Vanhanen et al., 2019), and the later expansion of the Battle Axe culture (BAC 2800-2400 cal BCE) in present-day Sweden, was a local development of the wide spread Corded Ware culture (CWC, 3000/2800-2400/1800 cal BCE) phenomenon seen around the whole Baltic Sea area (except the northern parts of the Gulf of Bothnia) [Fig. 1B] (e.g. Edenmo, 2008; Gimbutas, 1956; Iversen, 2016b; Knutsson, 1988; Kriiska, 2003, 2001; Kristiansen, 1989; Malmer, 2002, 1975, 1962; Nordqvist, 2016; von Hackwitz, 2009). Material culture from TRB, PWC, and BAC has also been found on Gotland (e.g. Bägerfeldt, 1992; Österholm, 1989), but the known burials from this time are associated with the TRB and PWC complexes. During the Scandinavian Late Neolithic period (LN, 2350-1750 cal BCE) a new homogenous cultural expression is found in Scandinavia and on Gotland called the Late Neolithic or Dagger culture with new burial customs and economy, and cultural remains from the TRB, PWC and BAC are no longer found.

*1.3 Gotland TRB and the Ansarve dolmen (c. 4000-2600 cal BCE)*

From around c. 4000 cal BCE there is evidence of TRB cultural remains and domesticated animals on Gotland (e.g. Lindqvist, 1997; Lindqvist and Possnert, 1999, 1997; Österholm, 1989). There are ten sites with TRB-pottery spread out across the island dated between c. 4000-3000 cal BCE, but only one confirmed TRB burial, the Ansarve dolmen in Tofta parish (Bägerfeldt, 1992; Lindqvist, 1997; Lithberg, 1914; Wallin, 2010). Another smaller dolmen (Lixarve), also located in Tofta parish in close vicinity (Wallin and Wehlin 2010) was recently excavated but no human remains was found as it most probably had been plundered in historical times (Wallin and Sjöstrand, 2018; Wallin and Wehlin, 2010). The Ansarve dolmen has been excavated twice, a test pit was dug in 1912 where some human remains were collected (n=8) (Lindqvist, 1997; Lithberg, 1914), and a more extensive excavation was performed in 1984 (Bägerfeldt, 1992). The human remains from the dolmen were fragmented and comingled, but it has been estimated that at least 31 individuals: men, women, and children of all ages were present (Bägerfeldt, 1992; Wallin, 2010; Wallin and Martinsson-Wallin, 1997). However, there could have been several more individuals present as the materials from both excavations have not been analyzed together. Seventeen individuals have now been dated which shows that the Ansarve dolmen was used continuously for more than 900 years (3500-2580 cal BCE, 95.4% CI), also with later use in the LN period (Fraser et al., 2018a, 2018b; Lindqvist, 1997). Stable isotope ($^{13}C/^{15}N$) analyses have shown that these individuals mainly showed terrestrial dietary signals although at varying degrees, however some slight marine and freshwater fish input was also noted in a few individuals (Fraser et al., 2018a; Lindqvist, 1997). Strontium isotope ($^{87}Sr/^{86}Sr$) analyses was performed for nine individuals across the main phase showing local Sr-signals, except the later dated individual (ans016) who was not native to the island (Fraser et al., 2018a).

Nine individuals from the main phase gave results for mitochondrial DNA (mtDNA) and was previously presented and discussed in Fraser et al. (2018a). The mt haplogroup composition (n = 9; K1a, T2b, J1c, HV0a, K2 and H7) of these individuals showed maternal continuity with individuals from Neolithic contexts in Scandinavia and central Europe, however some overlap with PWC was also noted. Although, the haplotypes (J1c8a, K2b1a, and H7d), have shown to be more common in later dated individuals associated with the LN & EBA cultures of central and northern Europe (e.g. Allentoft et al., 2015; Lipson et al., 2017; Olalde et al., 2018), the mt H7d haplotype found in the individual (ans016) have also been found in an earlier individual (HQU4, 3950-3400 BCE) from the Baalberge (TRB) culture in central Germany (Haak et al., 2015). However, many of the earlier mitochondrial haplogroups presented from archaeogenetic analyses were produced from PCR-based methodology, and thus are not possible to compare on the more refined level associated with high coverage mt genomes produced from next generation shotgun-generated analyses.

Six of these individuals from three different time periods of the main phase (ans003 and ans005; 3500-3110 cal BCE, ans008, ans014 and ans017; 3340-2930 cal BCE, and ans016; 2810-2580 cal BCE) generated genomic sequences consistent with ancestry from the EN Farmer (ENF) expansion, but also with some varying extent of HG admixture (Fraser, 2018; Sánchez-Quinto et al., 2019), as seen in other contemporaneous European Neolithic farmer individuals (e.g. Günther et al., 2015; Haak et al., 2015; Skoglund et al., 2014). Kinship relationships among the Ansarve burials (a second-degree relationship between SE_TRB_ans14 and SE_TRB_ans017) was previously investigated from using the READ software (Kuhn et al., 2018) and a detailed investigation of the pedigrees of the Ansarve and other megalithic burials was explored and discussed in Sanchez-Quinto et al. (2019).

The Y chromosomal I2a1b haplogroup found in the Ansarve burial (Fraser, 2018; Sánchez-Quinto et al., 2019) [For a discussion on the Y-chromosome haplogroups see Section 3 and Table S4], has also been found in individuals from MN megalithic burials in Scotland (Olalde et al., 2018; Sánchez-Quinto et al., 2019), as well as in WHG and SHG (Günther et al., 2018 e.g. Lazaridis et al., 2014; Mathieson et al., 2015; Mittnik et al.,

2018), and a later dated sub-Neolithic individual (vbj018, 2910-2690 cal BCE) from the Västerbjers PWC burial site on Gotland (Coutinho et al., 2020). Although the Y chromosome I2 lineage is common among individuals from MN megalithic burials (Sánchez-Quinto et al., 2019), the I2a1b1a1 lineage displayed in three of the males in the Ansarve burial has previously only been presented in the contemporaneous TRB associated individual (ESP24, 3360-3090 cal BCE) from present-day central Germany (Mathieson et al., 2015), showing interesting links between these TRB groups also on the uniparental markers.

##### *1.4 Gotland PWC (c. 3300-2300 cal BCE)*

Remains from the Pitted Ware culture (PWC) complex are found on the island from c. 3300 to 2300 cal BCE (Apel et al., 2018; Coutinho et al., 2020; Janzon, 1974; Norderäng, 2008; Österholm, 1989; Rundkvist et al., 2004; Wallin, 2016), in the form of c. 20 settlement sites of which several also contained burial grounds. The first PWC site Gullrum, Näs parish was excavated in 1893 (Lithberg, 1914), and new sites are still being found (Widerström and Norderäng, 2020). At least 200 burials have been excavated on eleven burial grounds on Gotland (Malmer 2002). The PWC performed flat grave inhumations and had a varied and rich ritualistic expression in their burial practices (Wallin, 2015; Wallin and Martinsson-Wallin, 2016). They were mostly buried in supine positions in single graves, but a few multiple burials, as well as package graves of disarticulated remains also occurred (Burenhult, 2002; Janzon, 1974; Molnar, 2008). Their clothes were adorned with tooth pendants (dog, seal, and pig) and bird bone beads and some had elaborate necklaces from large pig tusks (Burenhult, 2002; Janzon, 1974; Orascanin, 2010). Grave goods were commonly fishhooks, harpoons, pottery, stone tools, bone awls, but a few burials also contained artefacts associated with the Battle Axe Culture (Malmer, 1975, 1962). Some burials also include large numbers of pig jaws indicative of feasting at burials. These sub-Neolithic forager groups overlap the use of the Ansarve dolmen and were thus contemporaneous, but mainly lived of a marine economy, as seen in the zooarchaeological record, material culture, and stable isotopes (e.g. Coutinho et al., 2020; Eriksson, 2004; Eriksson et al., 2008; Lindqvist and Possnert, 1997; Martinsson-Wallin, 2008; Storå, 2001).

The PWC on Gotland was one the first Swedish archaeological cultures to be analyzed genetically, first with PCR based methods for mitochondrial DNA (mtDNA) (Malmström et al., 2015, 2010, 2009), and later from genomics through Next Generation Sequencing technology (Coutinho et al., 2020; Günther et al., 2015; Malmström et al., 2019; Skoglund et al., 2014, 2012). They are also part of the first study to show that the Neolithization was driven by migration (Skoglund et al., 2012). The PWC were the latest dated HG in Scandinavia and show ancestry with SHG, but also slight ANF admixture (Coutinho et al., 2020; Günther et al., 2018; Skoglund et al., 2014, 2012), which also is noted in the mitochondrial haplogroup composition; U4, U5, T2b, HV0, K1a, HV, V (Malmström et al., 2015). Recent Strontium isotope analyses of individuals buried at the Västerbjers PWC site only showed Sr-signals local to Gotland (Ahlström and Price, 2021), of which some individuals also were analyzed genetically in Coutinho et al. (2020).

##### *1.5 Presence of BAC artefacts and pottery on Gotland*

Continued usage of the EN megalithic burials are common, and it has previously been found that later usage of TRB tombs in Scandinavia also contain BAC pottery and artefacts (e.g. Iversen, 2016b; Sjögren, 2003). On Gotland there are some stray finds of cultural remains of BAC/CWC (Bägerfeldt, 1992; Rundkvist et al., 2004), and both BAC pottery and hocker-style burials have been found within the PWC (Janzon, 1974; Malmer, 1975, 1962; Palmgren and Martinsson-Wallin, 2015). However, no BAC/CWC settlements sites or burials have been located on the island and no BAC/CWC artefacts located in the Ansarve dolmen. Coutinho et al, (2020) investigated admixture within the PWC based on the BAC pottery and Hocker-style burials from three sites on the island (Västerbjers, Hemmor, and Ajvide). However, all individuals showed close genetic affinity, and no

Steppe-related admixture was found within any of these individuals. However, the possibility of BAC/CWC being present on Gotland in the 3rdML BCE still needs to be investigated.

Here we present new genomic data from two individuals from the Ansarve burial previously presented in (Fraser, 2018; Fraser et al., 2018a, 2018b; Sánchez-Quinto et al., 2019) [Main Table 1]. The materials have been stored at the Swedish History Museum (SHM31173) (1912 material), and at Uppsala university Campus Gotland (formerly, Gotland University, 1984 material). For additional information on the site and the material see (Bägerfeldt, 1992; Fraser et al., 2018a; Lindqvist, 1997; Martinsson-Wallin and Wallin, 2010; Wallin, 2010).

#### 2 DNA extraction, library building, authentication, and contamination control

**Table S1** Extraction and library information for ans010 and ans017

| Sample name | Sample | Yang extraction | BE library | Additional PCRs | DR library | Screaned & resequenced libraries | Reference |
|---|---|---|---|---|---|---|---|
| ans017 | M1 + M2, mand, dx | 13 | 13 | | | 30 | Sanchez-Quinto et al. 2019; Fraser et al 2018a |
| **ans017** | | | | | **12** | **20** | **This study** |
| ans010 | M1, max, dx | 3 | 4 | | | 7 | Fraser et al 2018a |
| **ans010** | | | | **12** | | **12** | **This study** |

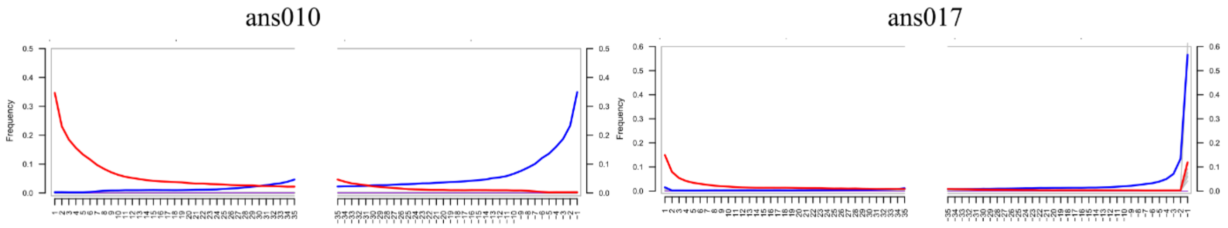

Figure S1. Deamination patterns at the fragment read-ends for blunt-end nuclear data for ans010 and ans017 (nonDR, ans017 DR data is not included in the plot).

**Table S2** Radiocarbon dating in chronological order rounded to the nearest 10th value, biological age, nuclear and mt genome coverage, average read length, biological sex for the seven Ansarve individuals investigated in this study

| Time period | Sample name | 14C, cal BCE (95% CI) | Bio. age | Genome coverage | mtDNA coverage | RL average | Bio. sex | Reference |
|---|---|---|---|---|---|---|---|---|
| MN | ans005 | 3500-3130 | 17-25 | 0,13 | 114,7 | 69 | F | Sanchez-Quinto et al. 2019; Fraser et al. 2018a |
| | ans003 | 3490-3110 | 25-35 | 0,14 | 300,9 | 77 | F | Sanchez-Quinto et al. 2019; Fraser et al. 2018a |
| | ans008 | 3340-3030 | 35-64 | 1,94 | 1462,4 | 56 | M | Sanchez-Quinto et al. 2019; Fraser et al. 2018a |
| | ans014 | 3330-2950 | 35-64 | 2,58 | 431,5 | 82 | M | Sanchez-Quinto et al. 2019; Fraser et al. 2018a |
| | ans017 | 3330-2930 | 13-17 | 6,80 | 491,0 | 94 | M | Sanchez-Quinto et al. 2019; Fraser et al. 2018a |
| | **ans017DR** | | | **27,42** | **1593,1** | **93** | | **This study** |
| | ans016 | 2810-2580* | 35-64 | 0,33 | 23,2 | 94 | M | Sanchez-Quinto et al. 2019; Fraser et al. 2018a |
| LN | ans010 | 2030-1890 | 18-60 | nd | 187,8 | 71 | nd | Fraser et al. 2018a |
| LN | **ans010** | | | **0,06** | **1397,4** | 71 | **M** | **This study** |

*Corrected for radiocarbon age (70 ± 40)

**Table S3.** Mitochondrial coverage and contamination estimates from the "Green Method" and Contamix, and nuclear contamination estimates (transversions only), VerifyBamID (Autosomes) and ANGSD (X-chromosome).

| | | Green Method | | | Contamix Bpielup | | | VerifyBamID | | | ANGSD | | |
|---|---|---|---|---|---|---|---|---|---|---|---|---|---|
| Sample | mtDNA coverage | Cont. est. [%] | #sites | Cont. interval [%] | P.est | C.I. | #Rds | P est. | Average cover at SNPs | # Reads used | Cont. est. [%] | SE | # SNPs |
| ans017 | 491,04 | 0,781 | 4 | 0.321 - 1.241 | 0,06 | 0.01 - 2.06 | 25061 | 1,461 | 7,55 | 29459851 | 0,7165 | 5,50569E-14 | 14184 |
| **ans017DR** | **1593,1** | **0,027** | **2** | **0.000 - 0.080** | **0,03** | **0.00 - 0.68** | **83816** | **0,487** | **18,91** | **56663087** | **0,2041** | **2,49839E-15** | **18815** |
| ans010 | 187,8 | 0,89 | 6 | 0.234 - 1.549 | nd | nd | nd | nd | nd | nd | nd | nd | nd |
| **ans010** | **1397,4** | **0,96** | **7** | **0.71 - 1.21** | **0,04** | **0.01 - 1.22** | **36113** | **nd** | **nd** | **nd** | **nd** | **nd** | **nd** |
| Bold this study | | | | | | | | | | | | | |

#### *3 Y chromosome analyses*

All single base substitutions from the International Society of Genetic Genealogy (ISOGG; http://isogg.org) version 15.73, 11 July 2020 were called using BAM files mapped to hg19 and SAMTools mpileup (version 1.3). Sites with a mapping quality and a base quality of at least 30 were extracted and insertions/deletions, and sites that displayed multiple alleles, were excluded. We investigated both transitions, transversions, plus A>T and G>C sites and report derived alleles within the called haplogroup. There was not enough data to determine the Y haplogroup for SE_LN_ans010.

The marker name, nucleotide position, type of substitution (ancestral>derived state), and coverage, are shown in brackets. When more than 20 haplogroup defining markers display derived allele states, we report three of them in the text.

**SE_TRB_ans017DR belongs to I2a1a2a1a1** and displayed derived allele states for the following: I2a1a2a1a1 (S2703:17361387; C>A, 10), I2a1a2a (S185:22513718, C>T, 19), I2a1a2 (S2768:22905944, G>C, 11; S2632:7317227, G>A, 8; S2621:2785672, A>G, 27; S2638:14074218, A>T, 14; S2679:16233135, A>G, 14; S2687:16594452, A>G, 16; S2702:17359886, A>C, 23; S2715:17893806, A>G, 18; S2722:18049134, C>T, 20; S328:15574052, G>A, 13 and M423:19096091, G>A, 13**),** I2a1 (PF3647:7879415, A>C, 15), I2 (PF3781:18700150, C>T, 22; S31:16638804, A>G, 15 and Z2638:8567995, G>A, 19), and for 166 markers defining haplogroup I (e.g. PF3715:14847792, A>C, 8 and PF3742:16354708, G>A, 9). Moreover, SE_TRB_ans017DR was also ancestral for I2a1b1b (Y6098:9647453, C>T, 13 and SK1254:6742730, T>C, 26), for I2a1b1 (PF3857:7716262, A>C, 24; S152:17570599, C>T, 16; M223:21717307, G>A, 13; L59:7113556, C>T, 32; S24:15517851, T>G, 17; S119:24475669, G>T, 14; PF3858:8353707, C>A, 23; U250:18888200, C>G, 19 and S117:16699334, C>G, 9), I2a1b (S33:18747493, G>C, 8; L181:19077754, G>T, 25; L368:6931594, C>T, 21; S30:13992338, C>G, 16; S23:7628484, C>T, 15 and S32:17493630, T>G, 18), which altogether supports I2a1a2a1a1.

The newly reported low coverage individual (SE_LN_ans010) was determined to belong to I2a2a.

**SE_LN_ans010 likely belongs to I2a2a** and displayed derived allele states for the following: I2a2a (PF3918:18694038; A>G, 1), I2a2 (S6728:16526276, T>C, 1), and two markers defining haplogroup I (PF3728:15536759, T>C, 1, and PF3645:7853028, C>A, 1). SE_LN_ans010 was also ancestral for I2a2c~(BY161750;14285512, A>G, 1) and I2a2b (PH3525:179262020, A>G, 1; M4099:23989924, C>G, 1; SK1274:6788888, A>C, 1; Z26408:22626296, C>A, 1), and I2b (BY32207:17460884, T>C, 1, supporting the I2a2a call despite the low coverage data.

The new high coverage ans017DR Y-chromosome data coincides with the previously reported substitutions and haplotype calls (Sánchez-Quinto et al., 2019) [Table S4]. However, in this analysis we used all single base substitutions from the International Society of Genetic Genealogy (ISOGG; http://isogg.org) version 15.73, 11 July 2020. The previously published Y-chromosome haplotypes in the four MN Ansarve males (ans008, ans014, ans017: I2a1b1a1a, and ans016: I2a1b) based on the derived substitutions (S2703:17361387, C>A and S2715:17893806, A>G, respectively), were determined from using the ISOGG version 11.329, 22 Dec 2017 [Table S4]. Moreover, as of ISOGG version 13.307, Dec 2018 these previously reported derived substitutions have has now been updated to belong to the I2a1a2a1a1 and I2a1a2 haplogroup lineages, respectively [Main Table 1 and Table S4]. These lineage updates do not change our conclusions for the Y-chromosome analyses in Sánchez-Quinto et al. (2019).

Furthermore, the haplotypes for SE_PWC_vbj013 (I2a1a-CTS595, C>T) and SE_PWC_vbj018 (I2a1b1-L161, C>T) have been restricted to the previous SNP call (I2a-L460, A>C) [Table S4] (from Table S.7 in Coutinho et al., 2020) based on that the previously published calls were made on only one C>T SNP. The I2a-L460 SNP has also been updated in the newer version to I2a1a [Main Table 1 and Table S4].

**Table S4.** Differences in reported Y-Chromosome haplogroup estimations for the Ansarve and Västerbjers individuals based on updates in ISSOGG 2020.

| **Use of dolmen** | **Sample** | Marker | **Ychr ISSOG 2017**[#] | **Ychr ISSOG 2020** |
|---|---|---|---|---|
| Main phase | ans008 | S2703 | I2a1b1a1a | I2a1a2a1a1 |
| | ans014 | S2703 | I2a1b1a1a | I2a1a2a1a1 |
| | ans017 | S2703 | I2a1b1a1a | I2a1a2a1a1 |
| | **ans017DR** | **S2703** | | **I2a1a2a1a1** |
| | ans016 | S2715 | I2a1b | I2a1a2a |
| Later use | **ans010** | **PF3918** | | **I2a2a** |
| PWC | vbj012 | L460 | I2a | I2a1a |
| | Vbj006 | L460 | I2a | I2a1a |
| | vbj018* | L460 | I2a | I2a1a |
| | vbj013* | L460 | I2a | I2a1a |

Bold result from this study

[#]Y Chromosome Haplogroups reported in Sánches-Quinto et al. 2019 and Coutinho et al. 2020.

*vbj018 was previously determined to I2a1b1-L161, and vbj013 to I2a1a-CTS595 in Coutinho et al. 2020, based on one C>T SNP and has been reduced to the I2a-L460 SNP marker.

### 4 Population genetic analyses

#### 4.1 Reference panels

Genetic data from the ancient individuals from the present study and previous publications [Extended Dataset1] were overlapped with four reference datasets as previously presented in (Günther et al., 2018). We only include ancient individuals with 5% or higher genome coverage, except for SE_LN_ans010 (~3%) as he is relevant to this study [see below].

Panel 1) We merged non-overlapping populations from Human Origins (Lazaridis et al., 2016) data sets comprising 616,938 SNPs genotyped in 2,582 modern individuals from 252 populations. Were used subsets of this dataset to perform PCA analyses and unsupervised ADMIXTURE v1.3 (Alexander et al., 2009) [see Main Methods for more information].

Panel 2) 3,862,510 biallelic transversion SNPs with a minor allele frequency of at least 1% in worldwide populations from the 1000 genomes project (Auton et al., 2015). This panel was used to perform *f-statistics* analyses (for SG-generated data only): *f3-statistics* for MDS and *f4-statistics* to further investigate genetic affinities within the Ansarve individuals as well as to other farmer groups.

Panel 3) 1,938,919 biallelic transversion SNPs with a minor allele frequency of at least 10% in Yorubans (YRI) from the 1000 genomes project (Auton et al., 2015) was used to perform CND analyses as described previously (Günther et al., 2018), and also ROHs on imputed data (Cassidy et al., 2020, 2016), and inbreeding coefficient (Gazal et al., 2014).

Panel 4) Autosomal SNPs from the 1.2M SNP captured array (Mathieson et al., 2015). Genetic data from all ancient individuals were screened for the 1,150,639 autosomal coordinates from this capture array, (and used for both SG- and CP-generated data). Ancient DNA data overlapped with this panel were used for qpGraph analyses (Patterson et al., 2012), DATES (Chintalapati et al., 2022; Narasimhan et al., 2019), and HapRoh (Ringbauer et al., 2021). For sex-bias analyses 49,704 SNPs on the X-chromosome were selected from the 1.2M SNP captured array.

##### *4.2 Group labels*

The labels used for demographic analyses are described in detail below and in Extended Dataset1. When performing demographic analyses, we refer to the ancient individuals from the same site as a group (in general) using a two-letter International Organization for Standardization (ISO) code for country, abbreviated cultural name or the time period(s), and site name; i.e. the individuals from the TRB associated Ansarve burial in present-day Sweden are labeled "SE_TRB_Ansarve" as a group, and when analyzed individually the site name is switched to the lab name (e.g. SE_TRB_ans003).

Additionally, to differentiate Genome-wide shotgun (SG) and capture (CP) generated data in the analyses the group labels for capture generated data also include a two-letter code "_CP", at the end of the label, SG generated data does not have any two-letter code ending. In cases where both CP and SG data is merged, i.e., for some of the groups in the PCA and ADMIXTURE analyses [Fig. 2], we add "_all" at the end of the label. In the case where grouped individuals are genetically similar but comes from different time periods and/or sites or cultural affinity and also are more distantly related to the individuals in question of this study, they are labeled after geography and culture or time period, (e.g., "DE_LBK_CP" and "HU_Neolithic_CP"). We also use the notation "/", e.g. "Central_EU_LN/CA" as an indication that the chronology extends through two time periods; however, the genetic profile of these individuals is broadly similar in qualitative terms.

In a few cases there is only one individual in a group in which case we have given them specific names in line with the other groups to further explain their affinity, i.e. "SE_LN_ans010" (ans010), "SE_TRB_Gökhem2" (Gok2), "SE_TRB_Saxtorp_CP" (Saxtorp5164), "DE_TRB_Baalberge_ESP_CP (ESP30), DE_TRB_Sorsum (Sorsum), DE_Esperstedt_CP (ESP24), DE_Salzmünde_CP (SALZ3B), and DE_TRB_Tangermünde_CP (TGM009).

When possible, already established nomenclature is used (e.g., the Mesolithic hunter-gatherers belonging to "WHG", "EHG" etc), whereas, new groups, and/or areas with several different groups, are labeled as explained above. E.g., for this study the Scandinavian hunter-gatherers are divided into three groups based on the geographic barriers ("SHG_Gotland", "SHG_Motala" and "SHG_Norway"), whereas the hunter-gatherers from Estonia, Latvia and Lithuania are grouped together geographically due to their similarity in genetic ancestry and small geographic distribution, but separated culturally and temporally (i.e. "Baltic_HG_Kunda_all" & "Baltic_HG_Narva_all"). The genetically analyzed Middle Neolithic hunter-gatherers of the present-day Baltic States ("Baltic_MN_all") were not further defined due to lack of information regarding cultural affiliation in some of these individuals, even though some show differences in genetic ancestry (EHG and/or WHG).

##### *4.3 Unsupervised ADMIXTURE*

Ancestry components were inferred using ADMIXTURE v1.3 (Alexander et al., 2009) and was based on 2392 individuals from 213 world-wide populations from the Human Origins Panel 1 [Supplementary section 4.1]. We subset Yoruba individuals to represent African genetic variation. This subset of individuals was merged with the ancient individuals described in Extended Dataset1. Only transversion SNPs were used in order to avoid biases due to cytosine deaminationThe dataset was filtered for linkage disequilibrium using PLINK v1.90b4.988,89 with parameters (--indep-pairwise 200 25 0.4), this retained 75,789 SNPs. ADMIXTURE v1.3 was run in 20 replicates with different random seeds for ancestral clusters from K=3 to K=16. Common signals between independent runs for each K were identified using the LargeKGreedy algorithm of CLUMPP90. CLUMPP predicted that K = 9 was the highest K at which >80% of the runs were consistent in their ancestral component prediction [Fig. 2B]. Clustering was visualized using pong 91.  Starting from K=7, the Western-European Mesolithic genetic component is first defined. From that K onwards, the Middle Neolithic individuals are modelled as a mixture of Mesolithic HG ancestry and the ANF ancestry component maximized in ancient

Anatolian agriculturalist. The Caucasian Hunter-gatherer (CHG) component is first defined at K=8 and can from that point onwards be found in the other individuals with Steppe-related ancestry, as well as the present-day European populations. . The results for all K's are shown in Extended Data Fig. 1.

#### 4.4 *f-statistics*

We used *f-statistics* (Patterson et al., 2012; Reich et al., 2009) to formally test the qualitative patterns we observed in the PCA and ADMIXTURE analyses.

##### *4.4.1 Affinity with other farmers: f3-statistics and Multi-Dimensional Scaling (MDS)*

Shared drift patterns were investigated both at the group and the individual level. At the group level we included CP-generated data and used Panel 4 [Fig. S2], while the individual level was investigated based on SG-generated data only using Panel 2 [Fig 3A]. In both cases individuals with low coverage per site were removed to avoid distortion of the MDS space. In the former analysis, DE_TRB_Baalberge_ESP_CP and DE_Salzmünde_CP were removed due to low coverage, while the group HU_Neolithic_CP was eliminated as data from same individuals, but shotgun-generated, was incorporated in the analysis. At the individual level we removed pairwise kin-related connections among the Ansarves (SE_TRB_ans014 and SE_TRB_ans017), and the PL_GAC_Koszyce and PL_Zlota_Ksiaznice groups (Schroeder et al., 2019).

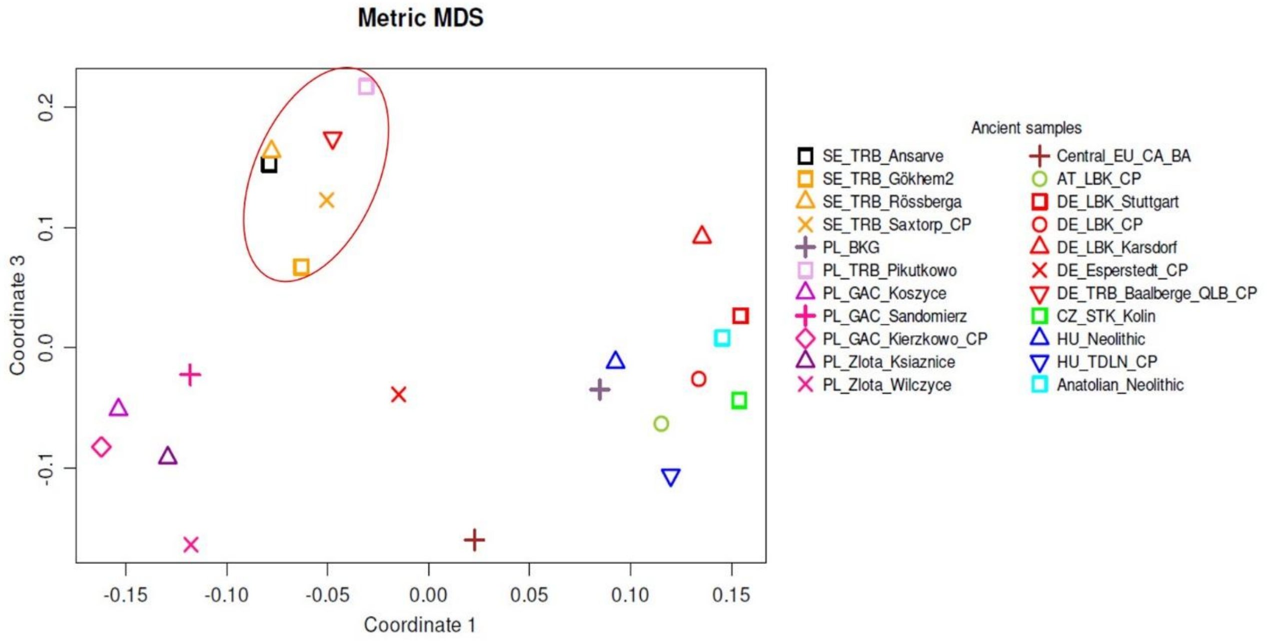

Figure S2. MDS plot of the 1st and 3rd coordinates display the connection of the different TRB groups from present-day Sweden, Germany, and Poland.

##### *4.4.2 Relationships among the Ansarve individuals: f4-statistics*

In order to further investigate the genetic affinities of the Ansarve individuals to other samples at the individual level, we performed *f4*-statistics analyses using SG sequenced genome-wide data overlapped with Panel 2. We investigated a topology of the form *f4* (O, X; Anatolian Neolithic, SE_TRB_Ansarve individual) employing Mbuti as an outgroup, and X a Neolithic farmer [Fig. S3].

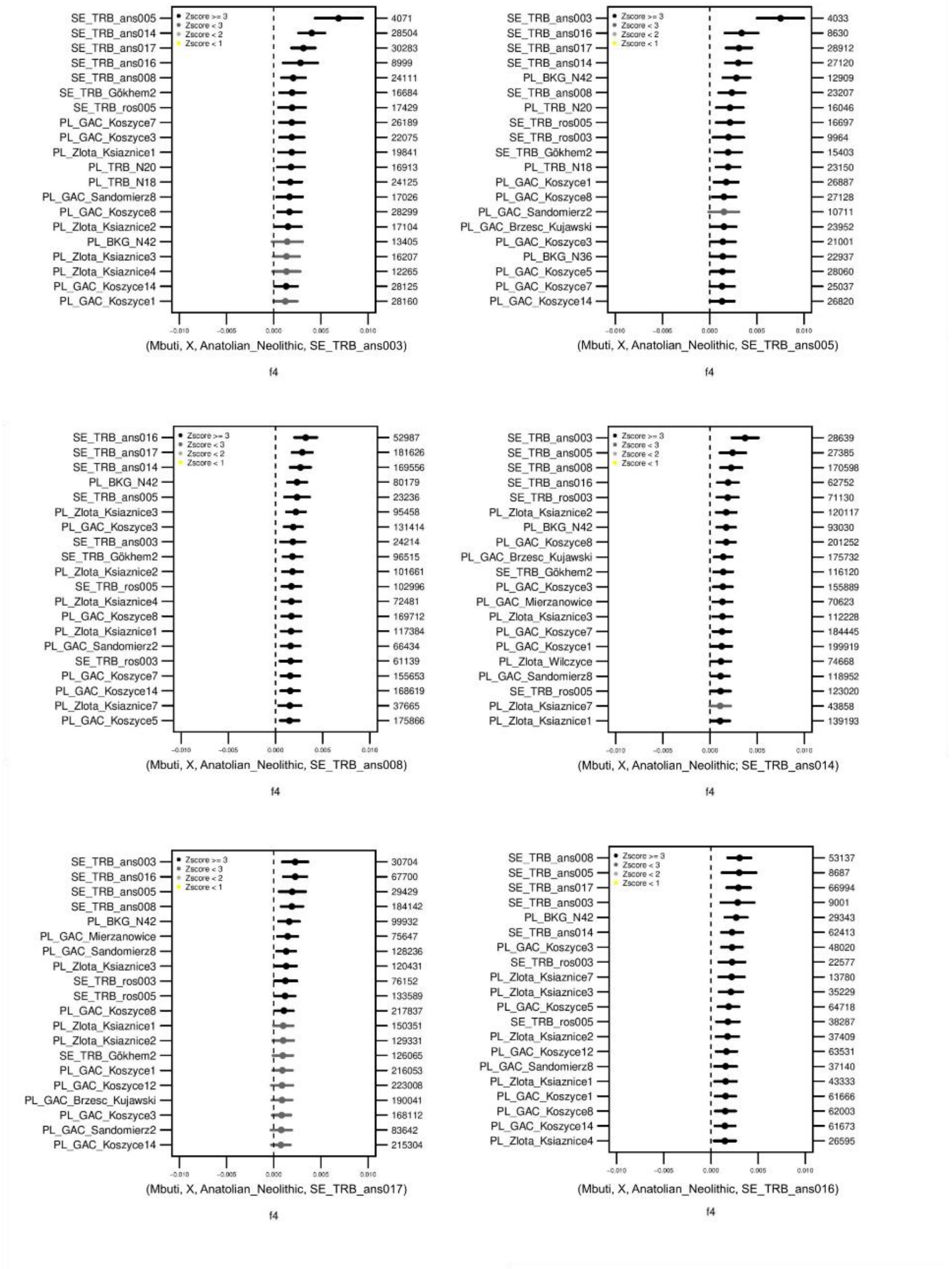

Figure S3. Top twenty results for each Ansarve individual in *f4*-statistics tests for degree of shared drift between the Ansarve and Neolithic farmer test individuals *f4*(Mbuti, X, Anatolian_Neolithic, SE_TRB_Ansarve individual). Error bars show two block-jackknife standard errors. The color represents the Z-score >= 3 (black), <= 3 (dark grey), <= 2 (light grey), < 0 (yellow). The number of SNPs included for each individual is reported to the right of each plot.

#### 4.5 Admixture graphs qpGraph

The Ansarve individuals received between 9 - 21% gene-flow from a SHG-related population in addition to their Farmer/WHG admixed ancestry [Fig. S4]. The PWC group displayed, in addition to their SHG-related ancestry, between 28 - 33% of Farmer/WHG admixed ancestry.

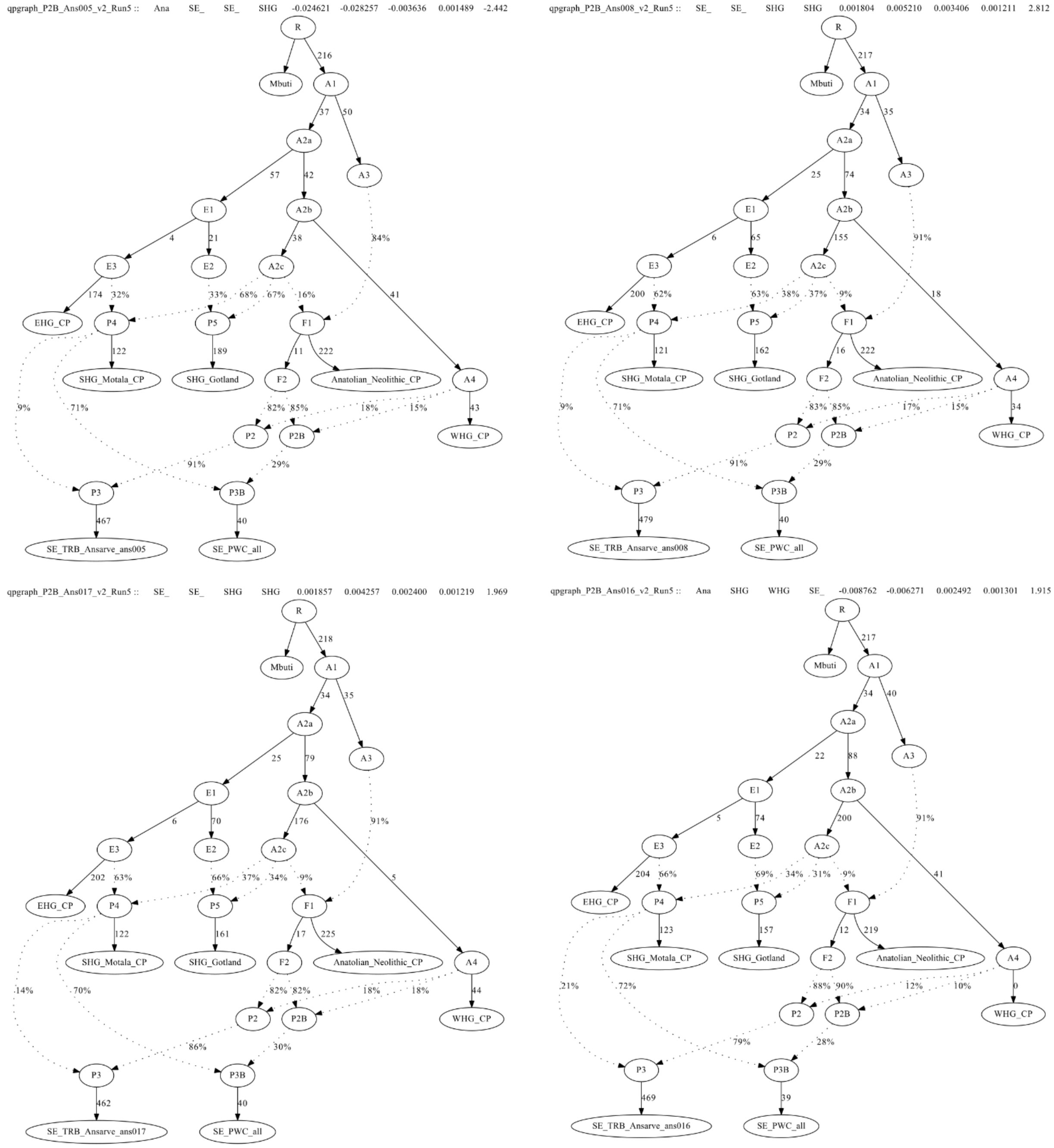

Figure S4. QpGraph results for the model (WHG, Anatolian_Neolithic and SHG_Motala) for both SE_TRB_Ansarve individuals (ans005, ans008, ans017, and ans016) and SE_PWC_all, with no admixture between the TRB and PWC. The models for SE-TRB-ans003 and SE_TRB-ans014 were rejected.

As in the previous model, we observe a 7 - 21% of SHG-related ancestry present in the Ansarve individuals, which seems to increase with time [Fig. S5]. The Ansarve individuals (ans008 and ans014) displayed similar values, although these models were rejected due to inner zero-drift branch results. The PWC individuals as a group can be modeled as deriving 31-36% Ansarve ancestry.

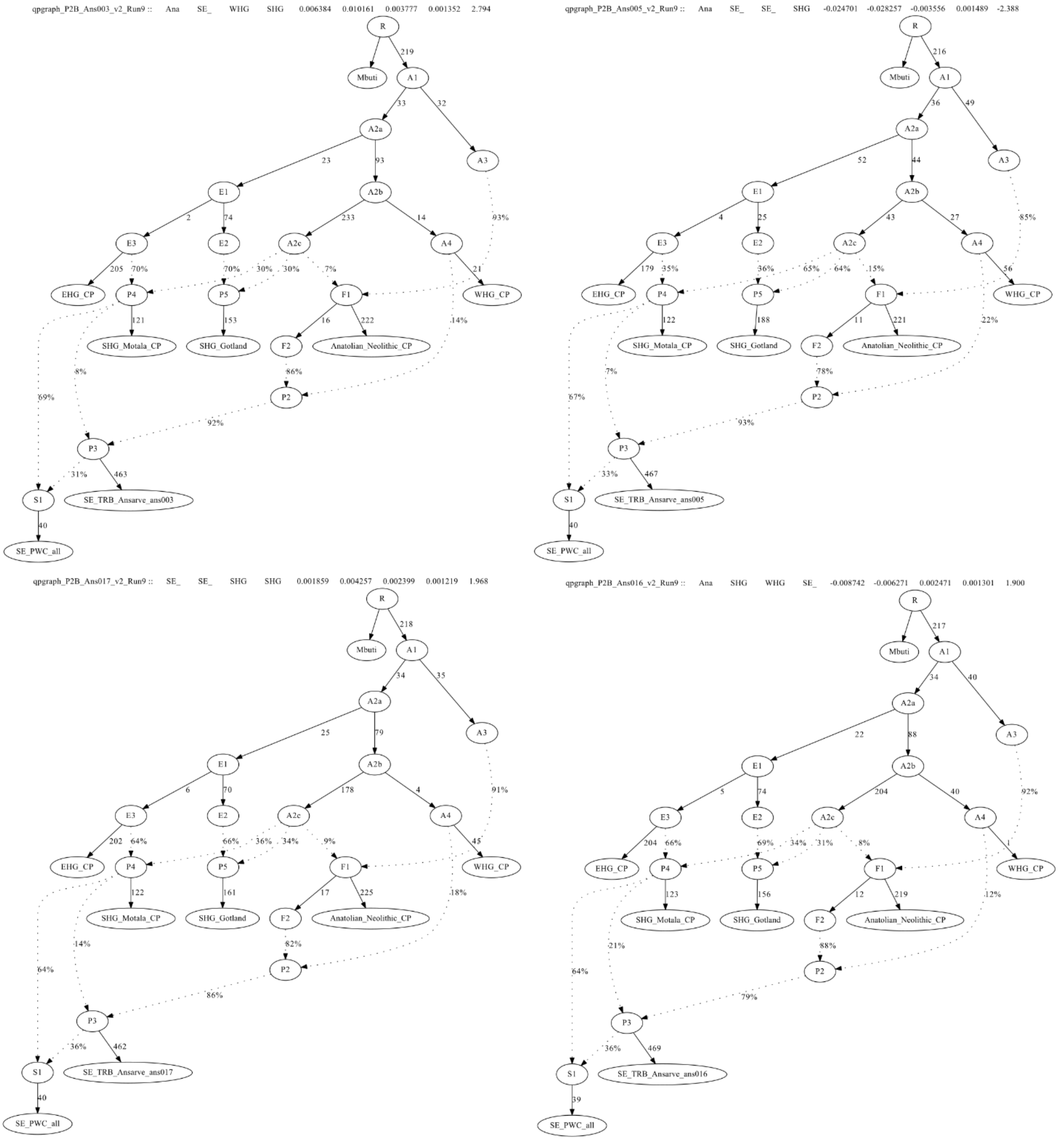

Figure S5. QpGraph results for the model (WHG, Anatolian_Neolithic and SHG_Motala) for SE_TRB_Ansarve individuals (ans005, ans008, ans017, and ans016), plus SHG_CP and a Ansarve TRB-related source into SE_PWC_all. The models for SE-TRB-ans008 and SE_TRB-ans014 were rejected.

#### 4.6 Conditional Nucleotide Diversity (CND)

We investigated genetic diversity for specific populations as computed in (Günther et al., 2015; Skoglund et al., 2014). To obtain an estimate of scaled diversity in ancient individuals we used at least two contemporaneous unrelated individuals from each archaeological site to account for low coverage sequencing, and kin-related individuals. To avoid ascertainment bias, post-mortem damage, as well as to increase the number of sites we only used SNPs from Panel 3 (Auton et al., 2015; Günther et al., 2015). Our diversity estimate is the average number of mismatches across all sites with coverage in both individuals. Standard error confidence interval was estimated from the distribution of mismatches for all pairs within a population using R [Fig. S6].

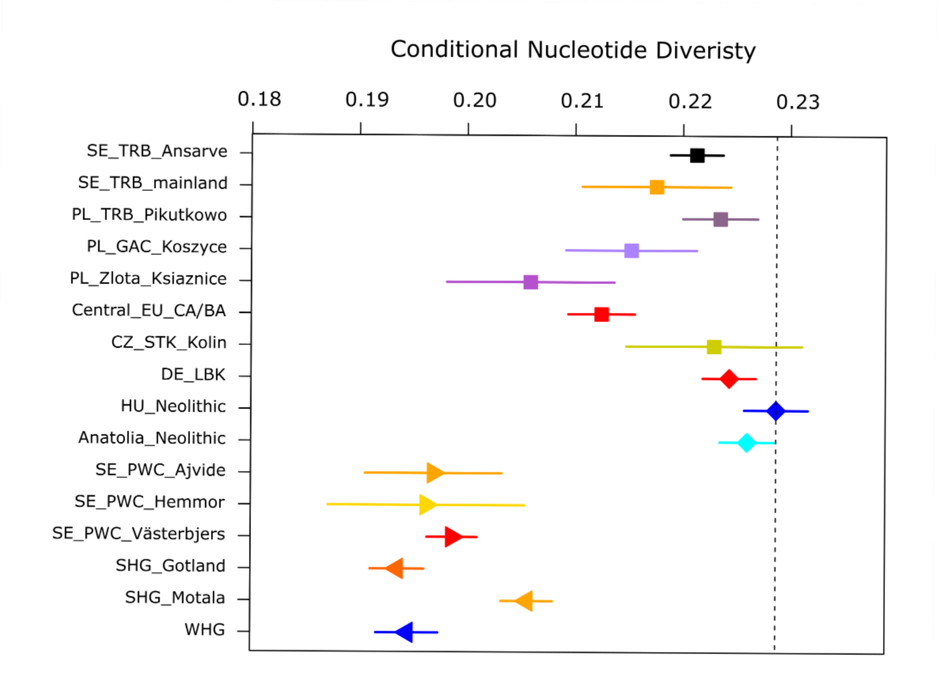

Figure S6. CND plot of 16 ancient farmer- and HG groups. Diversity was estimated using only individuals produced from SG-generated data.

#### 4.7 Runs of Homozygozity (ROH)

Genome-wide distributions of ROH are influenced by population history, as well as cultural practices such as endogamy (Cassidy et al., 2016). Therefore, analyzing the extent of homozygous segments across the genome can be informative on past events and effective population sizes. Many short segments of homozygous SNPs can be connected to historically small population sizes, while an excess of long runs of homozygosity suggests recent inbreeding.

##### *4.7.1 ROH using imputed data and high-coverage genomes*

ROHs was first investigated from imputed data in 55 ancient individuals following Cassidy et al.65 [Supplementary section 4.7.1]. Imputed data was overlapped with a set of 1.9 M transversions from panel 3 in order to maximize the information; monomorphic alleles had been previously removed using PLINKv1.90b4.9 92,93 using the a –maf flag. The length and number of runs of homozygosity were estimated using PLINK v1.90b4.9, using the following parameters: --homozyg-density 50 --homozyg-gap 100 --homozyg-kb 500 --homozyg-snp 50 --homozyg-window-het 1 --homozyg-window-snp 50 --homozyg-window-threshold 0.05 94. Individual ROH tracks were separated into two size bins over and under 1.6 Mb. Long ROH (>1.6 Mb) are informative about recent patterns of inbreeding, while short ROH (<1.6 Mb) shed light on ancient population constrictions. Following recommendations from Cassidy et al. 65 we estimated the fraction of the total genome under ROH for both track length categories, and visually displayed the differences for each individual between each category of ROH [Supplementary Fig. S7]. Moreover, genotype data for the five high-coverage ancient reference samples were included and compared to 1x down-sampled imputed data from the same individuals to highlight the high accuracy of the imputation process from Section 5 and Fig. S13.

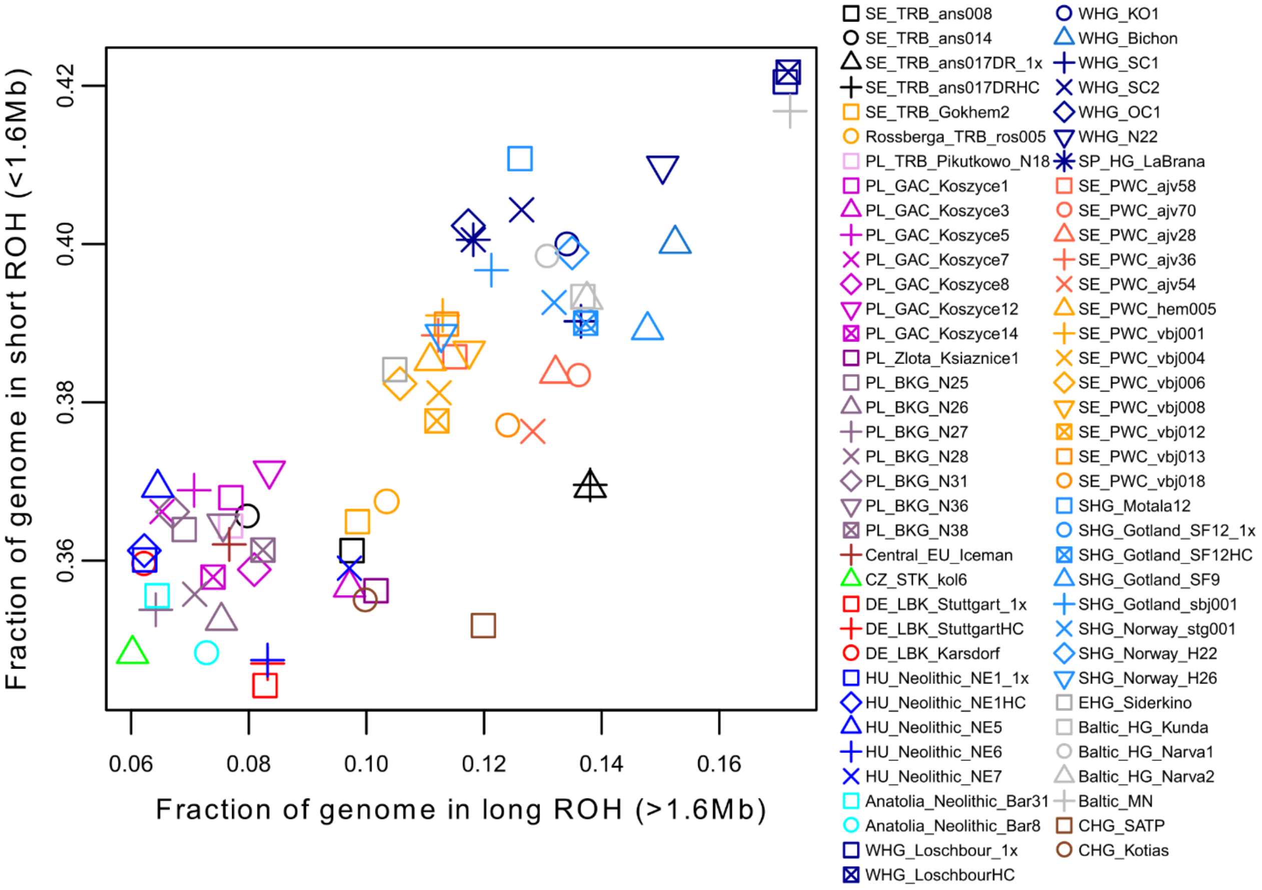

Figure S7. Short and long runs of homozygosity (ROH) spectra in 66 ancient individuals. Genotype data from the five high-coverage ancient reference samples were included as a reference to highlight the high accuracy of the imputation process.

##### *4.7.2 Estimating tracks ROH from low-coverage data HapROH*

Effective population sizes (Ne) were inferred both at the group and individual levels [Extended Tables S6.1 - S6.2], including only individuals without inbreeding. We report the total sum of ROH >4, >8, >12, >20 centiMorgan (cM) for the Scandinavian TRB and PWC [Fig S8].

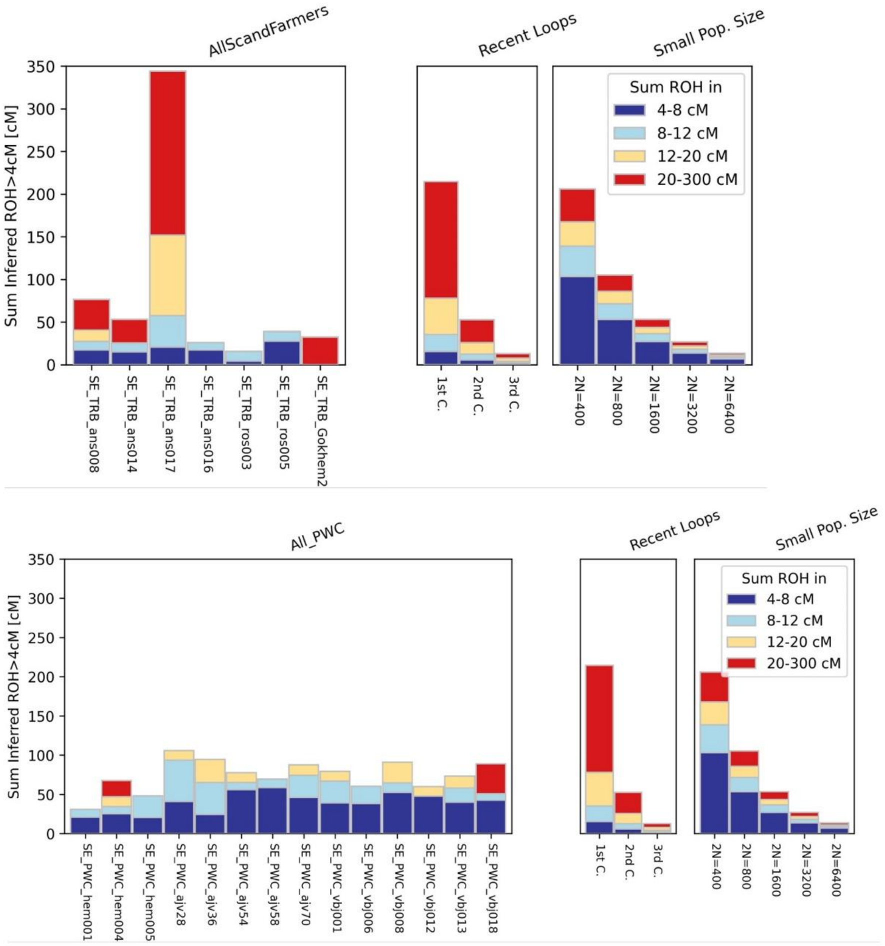

Figure S8. HapROH results of parental relatedness and effective populations sizes of Scandinavian TRB (Top) and PWC (Bottom) with >0.3x coverage.

In the case of the inbreed individual SE_TRB_Ansarve017 we provide a graphic display of ROH tracks across all chromosomes generated with *hapROH* [Fig. S9]. As expected for an inbreed individual who most likely is the product of 1st degree cousin admixture, ROH tracks (of at least 4cM in size) are long and present in several chromosomes.

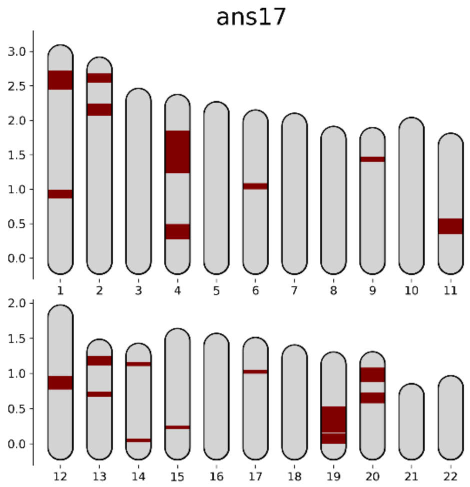

Figure S9. Karyotype ROH whole genome results for SE_TRB_ and017 from HapROH.

Although several pedigree scenarios for the 2$^{nd}$ degree relatedness between SE_TRB_ans017 and SE_TRB_ans014 are possible, we provide two versions where SE_TRB_ans017 parents also are 1$^{st}$ degree cousins [Fig. S10].

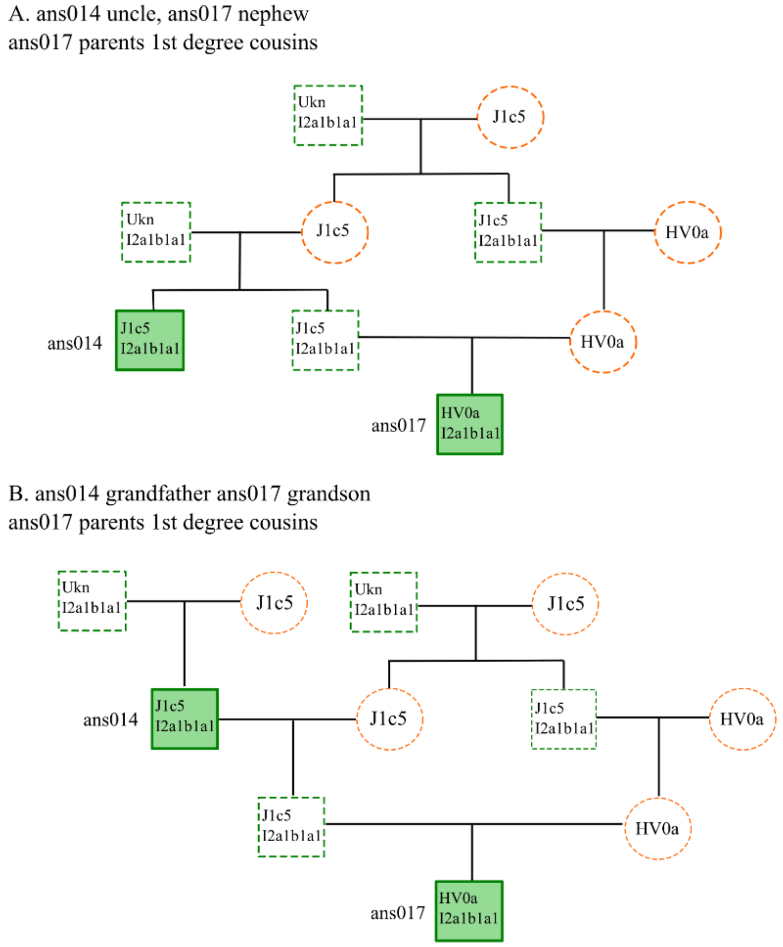

Figure S10. Possible pedigree scenarios for the 2$^{nd}$ degree relatedness between SE_TRB_ans014 and SE_TRB_ans017, where SE_TRB-ans017 parents are 1$^{st}$ degree cousins. A: Uncle/nephew pedigree. B: Grandfather/grandson pedigree. Dotted lines show unsampled individuals with inferred haplogroups, Ukn=unknown.

#### 4.8 Multiple Sequentially Markovian Coalescent (MSMC)

##### *4.8.1 Effective population size per individual as a function of time (PSMC)*

Effective population size per individual as a function of time "PSMC" was inferred using the MSMC method (Schiffels and Durbin, 2014) v. 0.1.0 [Supplementary section 4.8.1]. Analysis was performed on a dataset consisting of four present-day individuals from the HGDP (French, Han, Yoruba and Karitiana) (Prüfer et al., 2013) together with UDG-treated high-coverage data from this study (SE_TRB_ans017) plus four previously published individuals (SHG_Gotland_sf12, WHG_Loschbour, DE_LBK_Stuttgart, and RU_Ust'_Ishim), following the workflow reported in Günther et al. (2018). Input files were prepared using the scripts provided in msmc-tools (https://github.com/stschiff/msmc-tools). Briefly, using (i) "bamCaller.py" script, mask files were generated and using (ii) generate_multihetsep.py script, input files were prepared. MSMC was run with "--fixedRecombination -r 0.88" parameters and effective population size change was plotted with R using a mutation rate of 1.25x10$^{-8}$ and generation time of 30 years [Fig. S11].

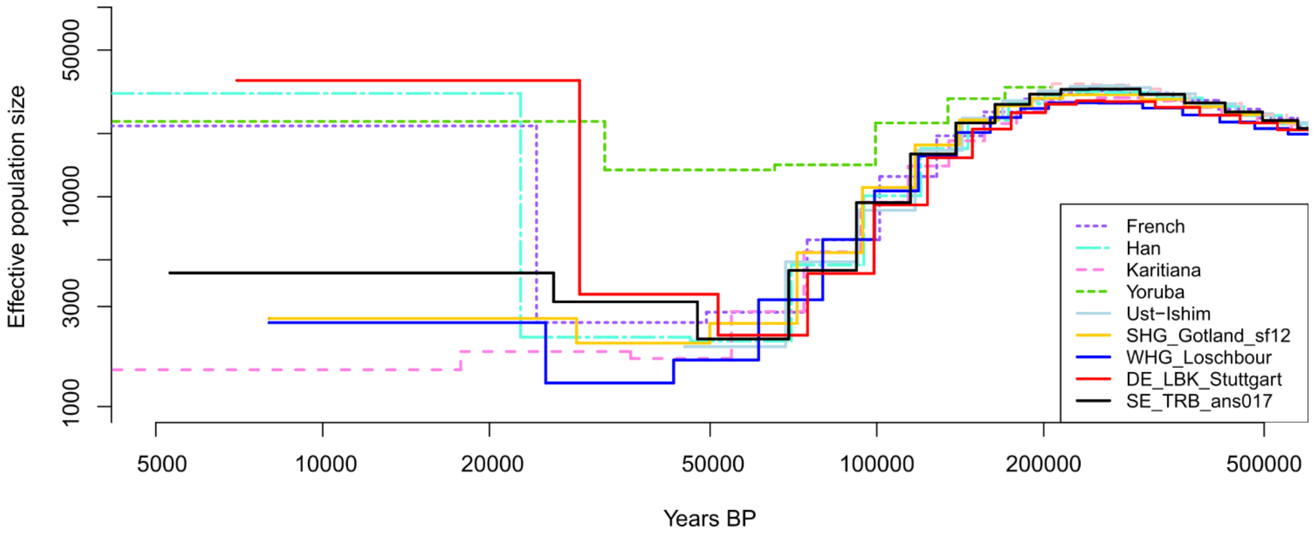

Figure S11**.** Effective population size changes over time for ancient and present-day populations. Effective population sizes were plotted assuming a generation time of 30 years and a mutation rate of 1.25x10e$^{-8}$. Radiocarbon dating results of the ancient individuals were used to shift the curves.

##### *4.8.2 Divergence over time using relative Cross Coalescence rate (rCCR)*

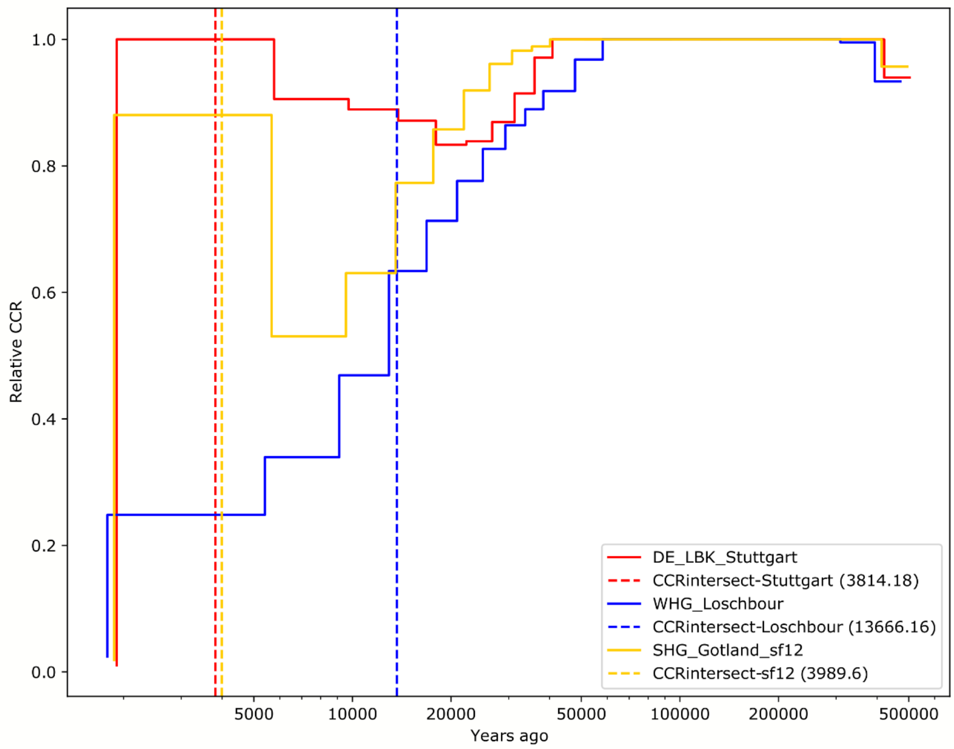

Figure S12. Relative Cross Coalescence Rate (rCCR) values of SE_TRB_ans017 when compared to WHG_Loschbour (blue), SHG_Gotland_SF12 (yellow), DE_LBK_Stuttgart (red). Values near to 1 suggest that the populations are panmitic or that there is a considerable amount of gene flow between them, while values near to 0 indicate that the populations are completely separated from each other. The dotted vertical lines refer to the intersect on time when rCCR drops to 0.5, which could be interpreted as time when the populations start to diverge from each other.

### 5 Imputation for analysis of ROHs

Imputation for the ROH analysis [section 4.7.1] was performed largely as described by Martiniano et al (2017). We selected a set of 55 prehistoric European individuals from hunter/gatherer and farmer contexts that have been produced by whole-genome shotgun sequencing with >0.7X coverage [Extended Table S9]. Such a strategy was performed in order to improve the imputation process as much as possible for the ancient DNA data, as this analysis uses both present-day and ancient data to assess genotype data for all ancient samples. Therefore, some of the ancient samples displayed in Extended Table 10 were only used as reference for the imputation process and were not further analyzed.

Therefore, prior to imputation genotype likelihoods were generated for each ancient sample separately using the Genome Analysis Toolkit (GATK), v3.5.0, tool UnifiedGenotyper, with arguments -mbq 30 --output_mode EMIT_ALL_SITES --genotyping_mode GENOTYPE_GIVEN_ALLELES. Genotypes were called using the 1000 Genomes phase 3 reference dataset (ftp.1000genomes.ebi.ac.uk/vol1/ftp/release/20130502/), filtered to exclude all non-biallelic SNPs, as well as chromosomes X and Y, resulting in a set of 77,818,182 markers. The resulting VCFs were filtered to exclude sites with missing genotypes, or where the SNP is a transition, and the most likely genotype could be derived from a possibly deaminated allele. This was done by removing sites where the reference allele was C and the alternative allele T, and the most likely genotype was 0/1 or 1/1, or correspondingly, if the reference allele was T and the alternative allele C, and the most likely genotype 0/0 or 0/1. The case of G>A deaminations was handled analogously. Finally, the individual files were merged and then split by chromosome using bcftools v1.6.

Imputation was performed using Beagle 4.0, with the 1000 Genomes Phase 3 v5a imputation panel of reference haplotypes and the GRCh37 genetic maps provided for Beagle at (http://bochet.gcc.biostat.washington.edu/beagle/). For more information regarding filtering criteria [see http://bochet.gcc.biostat.washington.edu/beagle/1000_Genomes_phase3_v5a/READ_ME_beagle_ref .] The reference panel was filtered to exclude all markers that are not biallelic SNPs. Imputation was performed based on genotype likelihoods (argument gl=<input>). For reasons of computational efficiency, Beagle was run separately on segments of 50,000 markers with an overlap of 25,000 markers, and the results were subsequently merged. The tools splitvcf.jar and mergevcf.jar provided at (https://faculty.washington.edu/browning/beagle_utilities/utilities.html) were used for this process. The chromosome-wise VCF files were finally concatenated using bcftools v1.6.

The performance of the imputation was evaluated by comparing the imputed genotypes of the five samples sequenced to high-coverage >20x (DE_LBK_Stuttgart, HU_Neolithic_NE1, WHG_Loschbour, SHG_Gotland_sf12 and the high-coverage damage-repaired data generated for SE_TRB_ans017), which had been downscaled to 1x with the corresponding genotypes called from high-coverage data (denoted the high-coverage VCF files here), using genotype concordance and PCA as described in Ausmees et al (2022). The high-coverage VCF files were filtered to keep variants with a minimum depth of 15 and a QUAL score of at least 50. Heterozygous genotypes were further filtered to keep only sites where both the depth of coverage of the reference allele and the alternative allele was at least 25% of the total depth. The imputed genotypes were filtered to only include SNPs with a minimum genotype probability of 0.99.

In order to assess genotype concordance, the set of compared markers for each sample (the intersection of the imputed and high-coverage VCFs) was split into two disjunct subsets: markers that were present in the downsampled VCF file (from the original high-coverage sample), and markers that were not. This was done to evaluate the difference in imputation quality on markers where the imputation algorithm had genotype likelihoods for, and markers that were missing in the sample that was imputed. For each sample, genotype concordance was evaluated for these two disjunct subsets of data using bcftools stats.

Regarding the assessment of the imputation process using PCA, the spatial location on a bidimensional space between the imputed genotypes to their corresponding high-coverage genotypes was compared, in the context of present-day European genetic variation from the Human Origins Panel v2 [see section 5.2]. For each of the 5 high-coverage samples, Human Origin v2 autosomic SNPs used to display present-day genetic structure were intersected with imputed markers, and markers from such sample's high-coverage VCF. PLINK v1.9 was used to intersect and merge ancient samples with the present-day genotypes. The PCA analysis was performed using the SMARTPCA software v16000, with diploid genotypes projecting all ancient DNA data, on top of the present-day European genetic variation [Fig. S13].

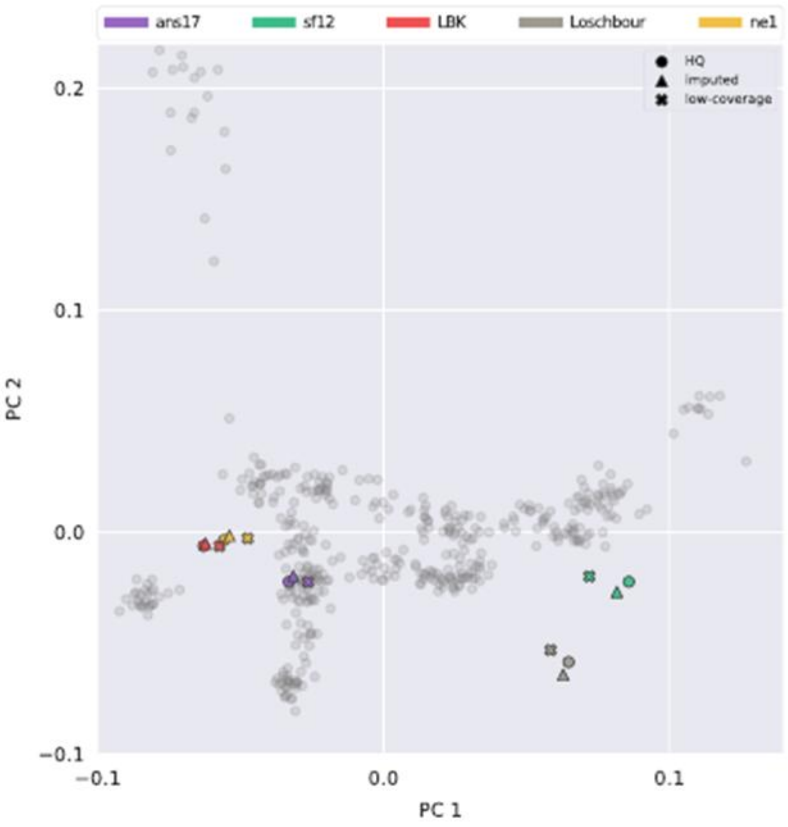

Figure S13. PCA comparing High quality coverage (HQ), and down sampled 1x imputed genomes for the five evaluation individuals. A reference PCA was defined based on genotypes of modern European samples from the Human Origins (Patterson et al., 2012) data set, after which the KDR method was used to estimate scores of ancient samples. Modern individuals are indicated by gray dots and ancient samples colored according to the legend. The imputed data points correspond to the genotypes that have been down sampled to 1x, filtered, and used as input to imputation, with a posterior filter of minimum genotype probability of 0.99 applied.

### 6 Discovery and annotation of novel variants in the SE_TRB_ans017DR genome

To discover novel variants in high coverage genome of SE_TRB_ans017 Genome Analysis Toolkit (GATK) (McKenna et al., 2010) version 3.5.0 was used. Prior to variant calling, base quality of "T" nucleotides at first five and and "A" nucleotides at last five positions of the sequencing reads was set to zero in its BAM file using a custom script from (Günther et al., 2018) to prevent incorrect genotype calling due to residual deamination. "AddOrReplaceReadGroups" function of Picard (Broad Institute. Picard tools *https://broadinstitute.github.io/picard/*, 2016) was used to add read group. Indel realignment was performed using "RealignerTargetCreator" and "IndelRealigner" modules of GATK. Indels reported in 1000 genomes project phase 1 (McVean, 2012) were used as reference set for indel realignment in both “RealignerTargetCreator” and “IndelRealigner” modules with “--known” parameter. Genotype calling was conducted using GATK’s “UnifiedGenotyper” with expressions “-stand_call_conf 50.0 & -stand_emit_conf 50.0 & -mbq 30 & -contamination 0.02 & --output_mode EMIT_ALL_SITES”. dbSNP version 142 was used as known SNPs with “--dbsnp” option. To filter the variants, “VariantFiltration” module of GATK was used with expressions “--filterExpression "QD < 3.0 || FS > 60.0 || MQ < 35.0 || MQRankSum < -12.5 || ReadPosRankSum < -8.0 || MQ0 >=5" and --genotypeFilterExpression "DP < 10 || GQ < ${1} || DP > 120"”. Using bedtools (Quinlan and Hall 2010), sites that are overlapping with 35 bp reads in Altai Neanderthal (Prüfer et al., 2013) were extracted. Transition to transversion ratio was calculated using “VariantEval“ module of GATK with parameters [Table S10].

**Table S10.** Transition to transversion ratio calculated for all variants using GATK's VariantEval.

| Novelty | nTi | nTv | tiTvRatio | TiTvRatioStandard |
|---|---|---|---|---|
| All | 1,359,778 | 644,616 | 2,11 | 2,01 |
| Known | 1,353,711 | 641,471 | 2,11 | 2,12 |
| Novel | 6,067 | 3,145 | 1,93 | 2,01 |

Finally, variants with sequencing depth smaller than 25x were filtered using vcftools (Danecek et al., 2011). This retained a total of 1,672,348 variants. Of these, 6,267 are novel and 1,667,081 are reported in dbSNP142. Functional annotation for all variants was performed using snpEff (v. 4.2) (Cingolani et al., 2012). The proportion of both dbSNP reported and novel variants in each functional annotation category is shown in Fig. S14.

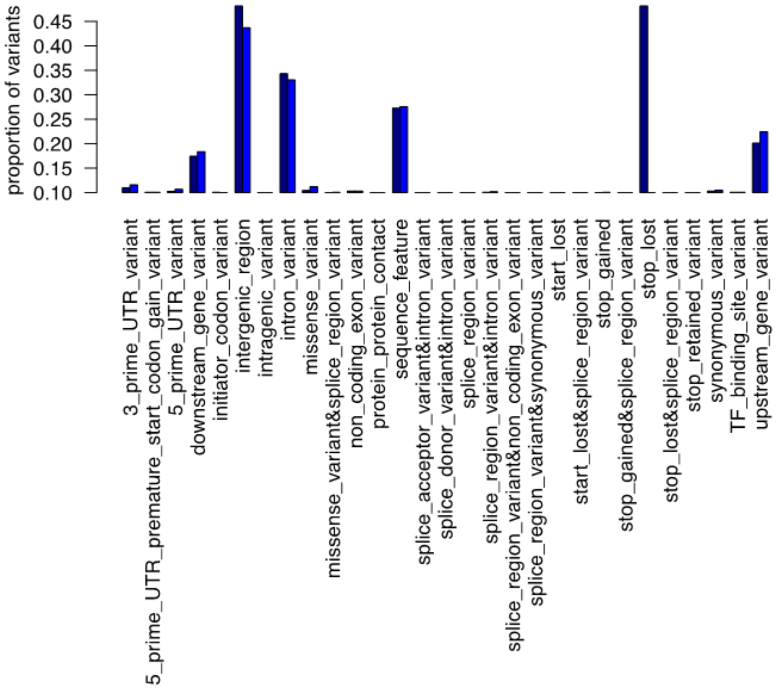

Figure S14. Proportion of each functional annotation category for dbSNP-reported variants (navy) and novel variants (light blue).

Six of the novel variants were classified as “high impact” based on SnpEff [Table S11].

**Table S11.** Novel variants found in SE_TRB_ans017 classified as "high impact" based on snpEff annotation

| Functional annotation | Gene | Chromosome | Position | Ref | Alt |
|---|---|---|---|---|---|
| Stop_gained | *SPATA16* | 3 | 172634172 | G | A |
| Splice_acceptor_variant&intron_variant | *HLA-DRA* | 6 | 32411035 | A | C |
| Stop_gained&splice_region_variant | *PPIL1* | 6 | 36824428 | G | A |
| Stop_gained | *MSRB2* | 10 | 23408037 | C | A |
| Splice_donor_variant&intron_variant | *UROS* | 10 | 127483447 | A | T |
| Start_lost&splice_region_variant | *KIAA1755* | 20 | 36888901 | A | C |

Deleteriousness of the novel variants were further evaluated using SIFT (**Ng 2001)** and Polyphen2 (Adzhubei et al., 2010). 17 of the 6,267 novel variants were predicted to be "damaging" with SIFT scores between 0 and 0.05. Of these, 13 were also predicted to be "probably- or possibly- damaging" by Polyphen2 [Table S12**]**.

**Table S12.** List of novel variants in SE_TRB_ans017 calssified as damaging by SIFT.

| Chr | Pos | Substitution | Gene | SIFT | SIFT | Polyphen2 | Polyphen2_Prediction |
|---|---|---|---|---|---|---|---|
| 1 | 27697206 | C/T | *FCN3* | 0,01 | Damaging | 0,984 | probably damaging |
| 3 | 193385022 | A/T | *OPA1* | 0 | Damaging | 1 | probably damaging |
| 5 | 112155011 | G/C | *APC* | 0,04 | Damaging | 0,081 | benign |
| 6 | 18122719 | C/G | *NHLRC1* | 0,01 | Damaging | 1 | probably damaging |
| 6 | 52957534 | G/A | *FBXO9* | 0,01 | Damaging | N/A | N/A |
| 8 | 11710891 | G/T | *CTSB* | 0,01 | Damaging | 0,315 | benign |
| 9 | 140087091 | G/A | *TPRN* | 0 | Damaging | 1 | probably damaging |
| 12 | 71972610 | C/T | *LGR5* | 0,02 | Damaging | 0,984 | probably damaging |
| 12 | 73015432 | G/A | *TRHDE* | 0 | Damaging | 0,994 | probably damaging |
| 12 | 132313113 | T/C | *MMP17* | 0,02 | Damaging | 0,953 | possibly damaging |
| 16 | 12798730 | C/T | *CPPED1* | 0 | Damaging | 0,996 | probably damaging |
| 17 | 27286451 | A/G | *SEZ6* | 0 | Damaging | 0,937 | possibly damaging |
| 19 | 14863154 | T/G | *EMR2* | 0,02 | Damaging | 0,481 | possibly damaging |
| 19 | 50304801 | C/T | *AP2A1* | 0,05 | Damaging | 1 | probably damaging |
| 19 | 54756736 | G/A | *LILRB5* | 0,01 | Damaging | 0,004 | benign |
| 20 | 36888901 | A/C | *KIAA1755* | 0 | Damaging | 0,995 | probably damaging |
| 21 | 34143706 | G/A | *PAXBP1* | 0,05 | Damaging | 0,978 | probably damaging |

### 7 Fastsimcoal modelling

Demographic inference of the Ansarve population's history was conducted using fastsimcoal2.7 (Excoffier et al., 2021) in accordance with the methods outlined by Marchi et al. (2022) [Supplementary section 7]. This analysis was performed on a panel of high-coverage genomes of ancient western Eurasian individuals, with WHG_Loschbour representing WHG, SHG_Gotland_sf12 representing SHG, DE_LBK_Stuttgart representing ENF, and SE_TRB_ans017 representing the Ansarve population.

The dataset was filtered to exclude any sites with missing data, sites where the reference alleles differ between chimpanzee and gorilla reference genomes, CpG sites or regions, and sites located in genomic regions with a recombination rate below 1 cM/Mb. To focus on neutral sites suitable for demographic inference, only BGC-free A>T and G>C polymorphic sites were retained, resulting to a final panel consisting of 112,366,543 sites.

Four models were tested: (1) no pulse admixture into the Ansarve population, (2) WHG pulse admixture into LBK + the ancestors of Ansarve, (3) SHG pulse admixture into Ansarve, and (4) WHG pulse admixture into LBK+ the ancestors of Ansarve, plus SHG pulse admixture into Ansarve [Extended Table S13 and Fig. S15 A].

Unsampled meta populations labelled as “Meta ANE” (Ancestral North Eurasian), “Meta WHG”, “Meta EHG”, and “Meta Neolithic” were also incorporated into the models. For each model, parameter estimates were obtained by maximizing the model's likelihood across 50 independent runs of fastsimcoal, each involving 50 Expectation Conditional Maximization (ECM) cycles and 500,000 coalescent simulations to estimate the expected Site Frequency Spectrum (SFS) (command line used for this step was: fsc -t Panel.tpl -n500000 -d -e Panel.est -M -L50 -q -C5 --multiSFS --logprecision 18 -c16 -B16).

For each model, the maximum likelihood (ML) parameters were recorded based on the run with the highest likelihood from the 50 independent runs [Fig. S15 B]. The relative likelihood and Akaike Information Criterion (AIC) were calculated for model comparison. We further validated the results by comparing likelihood distributions using 10 million coalescent simulations, repeated 100 times (command line used for validation: fsc -i Panel_maxL.par -R100 -n10000000 -d -u -C5 --logprecision 18 -q -c16 -B16).

To obtain confidence intervals around the ML parameter estimates for the best-fitting model (WHG and SHG pulse admixture into Ansarve), a parametric bootstrap approach was employed. Using the estimated ML parameters, we generated 100 SFS (command line used was: fsc -i Panel.par -n100 -j -d -s0 -x –I -q -u -c16 -B16). For each of the 100 bootstrapped SFS, parameters were re-estimated with 20 independent runs, starting from the ML parameter values, and each run consisted of 60 ECM cycles with 500,000 simulations to estimate the SFS and model likelihood (command line used was: fsc -t Panel.tpl -n500000 -d -e Panel.est --nitvalues Panel.pv -M -L60 -q -C5 --multiSFS --logprecision 18 -c16 -B16). The 95% confidence intervals were determined by calculating the 2.5% and 97.5% quantiles of the parameter distribution across the 100 re-estimated ML parameters.

We estimated the total HG ancestry (SHG + WHG) in SE_TRB_ans017 based on the model with the highest likelihood. Initially, the Meta Neolithic population received an influx of WHG ancestry at 21.52% [95% CI: 3.44% – 25.98%]. Later, the common ancestor of Ansarve and LBK experienced an additional WHG ancestry influx of 9.03% [95% CI: 3.73% – 13.16%]. This resulted in a total WHG ancestry of **28.61%** [95% CI: 7.04% – 35.72%], calculated as: 9.03 + (100−9.03) × 0.2152 = 28.61%. Furthermore, Ansarve received an additional SHG admixture of 13.55% [95% CI: 4.52% – 13.99%]. This led to a total hunter-gatherer ancestry of **38.27%** [95% CI: 11.24% – 44.71%], calculated as: 13.55 + (100−13.55) × 0.286 = 38.27%

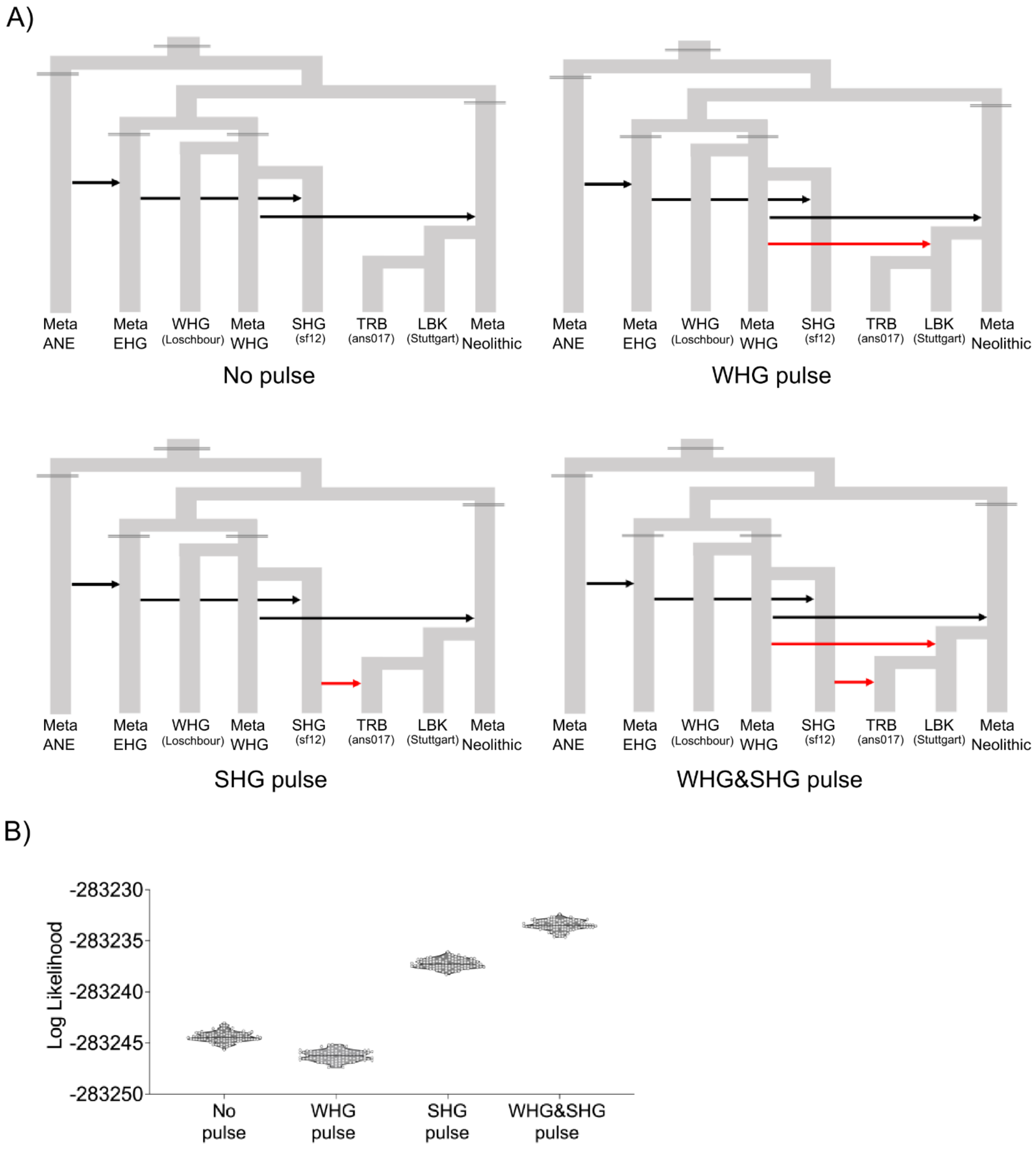

Figure S15. FastSimcole2 analysis of a high-coverage panel with WHG_Loschbour representing Western Hunter-Gatherers, SHG_Gotland_sf12 representing Scandinavian Hunter-Gatherers, DE_LBK_Stuttgart representing Early European Farmers, and SE_TRB_ans017 representing the Ansarve population, and unsampled meta populations labelled as “Meta ANE” (Ancestral North Eurasian), “Meta WHG”, “Meta EHG”, and “Meta Neolithic”. A: The four different models: (1) no pulse admixture into the Ansarve population, (2) WHG pulse admixture into LBK+Ansarve ancestor, (3) SHG pulse admixture into Ansarve, and (4) WHG pulse admixture into LBK+Ansarve ancestor, plus SHG, pulse admixture into Ansarve [Extended Table S13]. B: Relative likelihood for best fitting model showing that Model 4 had the best fit.

#### 8 Metagenomic screening for Yersinia pestis

We screened 180 aDNA libraries from 11 individuals from the Ansarve burial, (Fraser et al., 2018a; Sánchez-Quinto et al., 2019; and this study) and 603 libraries from 19 PWC individuals, in order to assess if we could identify this pathogen in the metagenomic data for Gotland's farmer and foragers communities.

##### *8.1 Metagenomic screening strategy*

Metagenomic data was screened using *malt* (Herbig et al., 2016) to look for potential pathogen sequences by using a reference genome collection that contains archaeal, bacterial, fungal, and viral species (n = 6, 133) from reference and representative genomes of the RefSeq database (O'Leary et al., 2016). Before indexing this reference genome collection, *Dustmasker* (Morgulis et al., 2006) was used to mask repetitive regions of the full-length reference genomes to avoid false positive classifications. Then we aligned DNA reads to the indexed reference genome collection using *malt* with the *semi-global* flag on. This tool aligns DNA reads to the reference genome collection, and using a "lowest common ancestor" method, assigns sequences to a specific taxonomical node.

An initial analysis using MEGAN (Huson et al., 2016) on SE_TRB_ans003 and SE_LN_ans010 showed presence of *Y. pestis* and *Y. enterocolitica*. We found 697 reads that mapped to *Y. pestis* in SE_TRB_ans003, and we identified 7,918 reads aligned to *Y. pestis*, and 10,455 reads aligned to *Y. enterocolitica* in SE_LN_ans010. In light of these results, we decided to map the 783 screened sequencing libraries from the 11 Ansarve and 21 PWC individuals to the reference genomes of *Y. pestis*, *Y. enterocolitica* and *Y. pseudotuberculosis* (see below) in order to authenticate these observations.

Out of 32 individuals we observed the presence of reads mapping to the three investigated microorganisms in five individuals: SE_TRB_ans003, SE_TRB_ans005, SE_TRB_ans007, SE_LN_ans010, and SE_PWC_ajv58 [Table S14]. Mapping was done using *bwa*, forcing reads to map on their entire length (-l 1024), with a mapping quality > 30 (-q 30) and eliminating reads shorter than 30bp. Duplicate reads were then removed using picard tools *MarkDuplicates*. Importantly, for the case of *Y. pestis*, in three of the individuals, we also found reads mapping into the three plasmids that are characteristic of this species. These plasmids have been so far only found in *Y. pestis*, are involved in pathogenicity and their detection has been considered as strong evidence supporting the presence of this pathogen in ancient samples.

**Table S14.** Number of unique reads (after removing duplicates) mapping on the genomes of the two potential pathogens identified during the screening. The mapping against Y. pseudotuberculosis was performed to authenticate reads mapped on the Y. pestis genome.

| NCBI Accession | SE_PWC_ajv58 | SE_TRB_ans003 | SE_TRB_ans005 | SE_TRB_ans007 | SE_LN_ans010 | Ref. Sequence Annotation |
|---|---|---|---|---|---|---|
| ***Y. enterocolitica strain NW56*** | | | | | | |
| CP107102.1 | 518 | 257 | 66 | 62 | 7338 | Chromosome |
| CP107103.1 | | | | | 1 | Plasmid unnamed 1 |
| CP107104.1 | | | | | | Plasmid unnamed 2 |
| ***Y. pseudotuberculosis strain IP32953*** | | | | | | |
| NC_006155.1 | 924 | 466 | 131 | 96 | 4462 | Chromosome |
| NC_006153.2 | 15 | 4 | 1 | 4 | 133 | pYV plasmid |
| NC_006154.1 | | | | | | Cryptic plasmid |
| ***Y. pestis strain CO92*** | | | | | | |
| NC_003143.1 | 972 | 483 | 138 | 94 | 4677 | Chromosome |
| NC_003131.1 | 15 | 4 | 1 | 4 | 139 | Plasmid pCD1 |
| NC_003134.1 | 16 | 3 | | 2 | 93 | Plasmid pMT1 |
| NC_003132.1 | 16 | 3 | | | 45 | Plasmid pPCP1 |

To further authenticate *Y. pestis* hits, we also included *Y. pseudotuberculosis* in the analyses because it is a closely related species from which *Y. pestis* diverged 30-50 kya, so mapping reads must be much closer to *Y.*

*pestis* than to *Y. pseudotuberculosis* to be considered valid. To evaluate the relatedness to each species, we estimated the edit distance (ED) of the reads mapping to *Y. pestis* against the genomes of both species [Fig. S16]. This value indicates the number of single nucleotide substitutions per mapped read. Edit distances were retrieved from the bam files corresponding to the mapping to each reference genome (NM tag) using samtools, and the distribution of read counts mapping at different edit distances to the reference were plotted in a histogram. These analyses revealed the authentic taxonomic classification of *Y. pestis* reads from the SE_LN_ans010, as evidenced by a larger number of reads mapping with ED = 0, followed by ED = 1, while more reads mapping with ED >2 on *Y. pseudotuberculosis*. However, the results of the other four individuals were less conclusive, as evidenced by a higher proportion of reads with ED ≥ 1 in relation to those with ED = 0. Nonetheless, these results could also be explained by the much lower number of mapping reads, a scenario in which the presence of just a few environmental contaminants reads can have a strong effect on the observed ED distribution. On the other hand, the authentication of *Y. enterocolitica* reads showed that while there were reads mapping into the reference genome of this species (strain NW56) in all samples, the ED results revealed that only SE_LN_ans010 showed a very strong affinity to this pathogen, with 7338 unique reads mapping into the genome, 83% of which with ED ≤ 1. Altogether, these results could indicate that the individual SE_LN_ans010 was co-infected with *Y. pestis* and *Y. enterocolitica*. However, whole genome data in addition to phylogenetic analyses to identify the strain and its relationship to other *Y. enterocolitica* samples, would be needed in order to rule out potential contamination with unknown environmental *Yersinia* species.

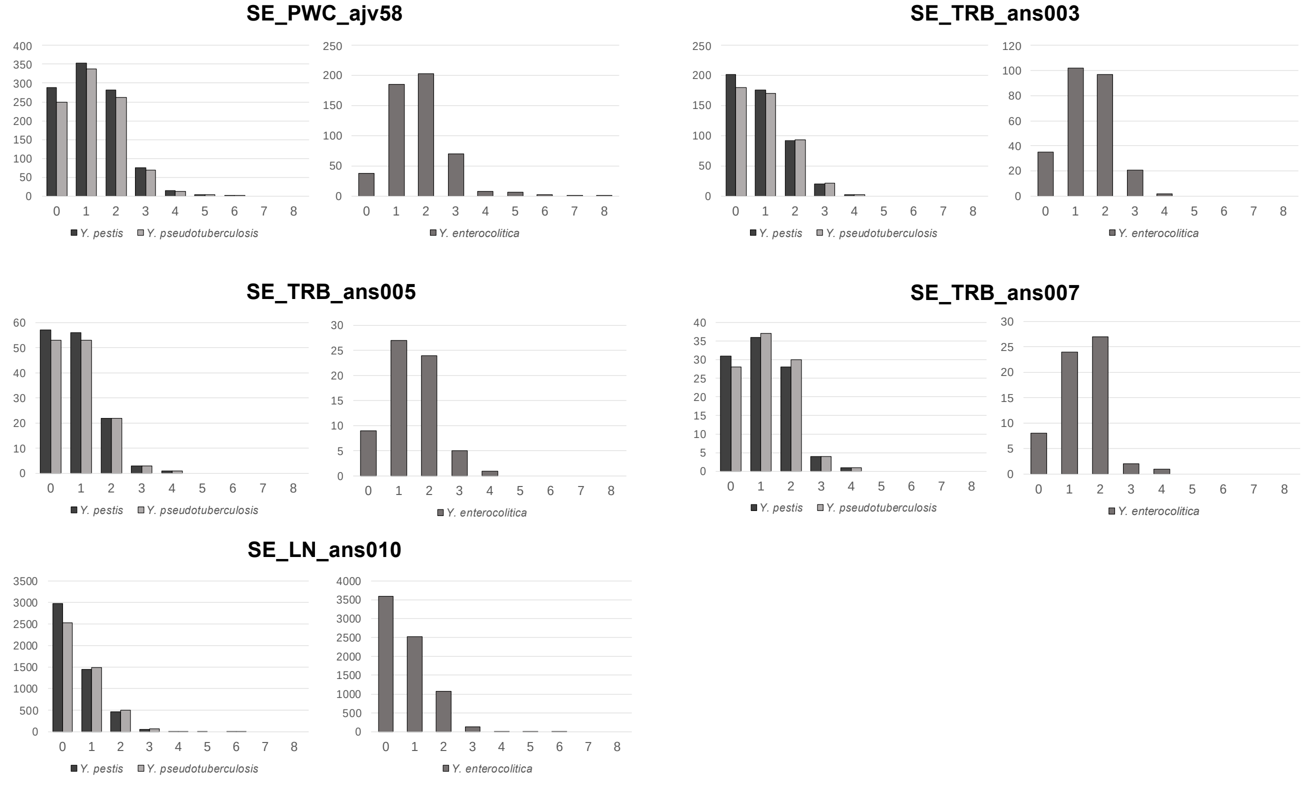

Figure S16. Distribution of edit-distances of the uniquely mapping reads to the reference genomes of *Y. pestis CO92* and *Y. enterocolitica NW56*, the two potential pathogen species detected at the pathogen screening stage. Results against the *Y. pseudotuberculosis* genome were also computed to better authenticate *Y. pestis* hits (rationale explained in the text).

While the ED patterns against *Y. enterocolitica* clearly showed a non-specific mapping of reads in four of the samples (SE_PWC_ajv58, SE_TRB_ans003, SE_TRB_ans005, SE_TRB_ans007), likely explained by sequences originating from environmental contaminants, the authenticity of *Y. pestis* reads in these samples was inconclusive. To further investigate the origins of reads mapping into this species, we conducted a competitive mapping of all reads in the samples, against the reference genomes of 27 species of the *Yersinia* genus (all the species listed in the NCBI taxonomy DB), using the same strategy and parameters as described above for the previous mappings. As expected, the number of uniquely and specifically mapping reads got drastically reduced [Table S15], turning two of the samples impossible to authenticate with this strategy (SE_TRB_ans005 and SE_TRB_ans007 with 1 and 3 mapping reads respectively), while results for SE_LN_ans010 were further authenticated for both *Y. enterocolitica* and *Y. pestis*.

**Table S15.** Read counts of the competitive mapping to the *Yersinia strains: enterocolitica*, *pestis*, and *pseudotuberculosis* to SE_TRB_ans003, SE_TRB_ans005, SE_TRB_ans007, SE_PWC_ajv58, SE_LN_ans010

| Species | SE_TRB_ans003 | SE_TRB_ans005 | SE_TRB_ans007 | SE_PWC_ajv58 | SE_LN_ans010 |
|---|---|---|---|---|---|
| *Y. enterocolitica* | 16 | 1 | 1 | 134 | 2771 |
| *Y. pestis* | 28 | 1 | 3 | 69 | 317 |
| *Y. pseudotuberculosis* | 10 | 0 | 2 | 4 | 24 |

We cannot rule out the possibility that there is *Y. enterocolitica* in SE_LN_ans010, as both the observed breath of coverage and the observed depth of coverage are almost identical to what would be expected for this number of reads [Extended Table S16 and Fig. S17].

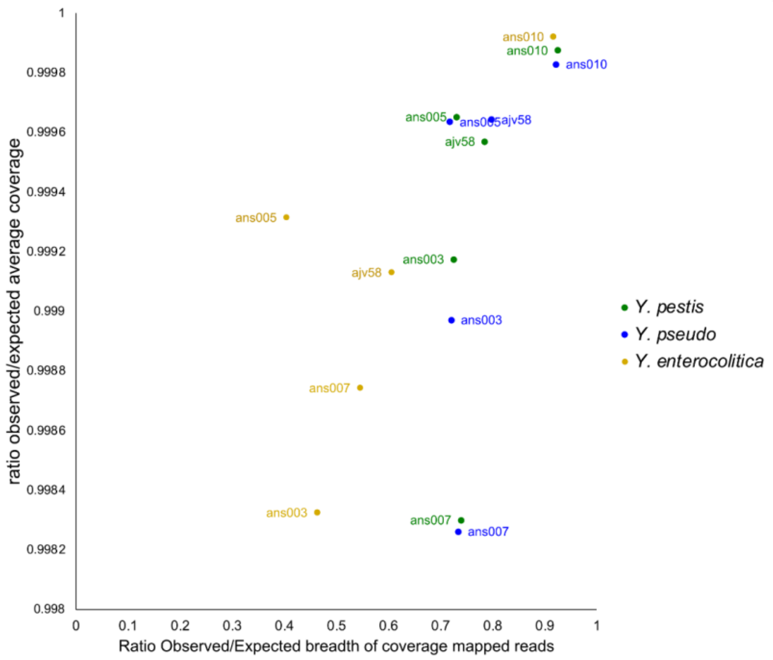

Figure S17. “Ratio Observed/Expected breadth of coverage” plotted against “Ratio observed/expected average depth of coverage” for mapped reads for each of the three species of Yersinia (*Y. enterocolitica*, *Y. pestis*, and *Y. pseudotuberculosis*) from Extended Table S16.

It also appears as the reads mapping on *Y. enterocolitica* from SE_LN_ans010 seem to be quite homogeneously distributed (the peaks are likely highly conserved genes, such as the 16 rRNA or similar [Fig. S18]. However, as mentioned before whole genome data and a proper phylogenetic analysis would be needed to confirm the presence of this pathogen and rule out potential contaminations with yet another unknown environmental Yersinia species.

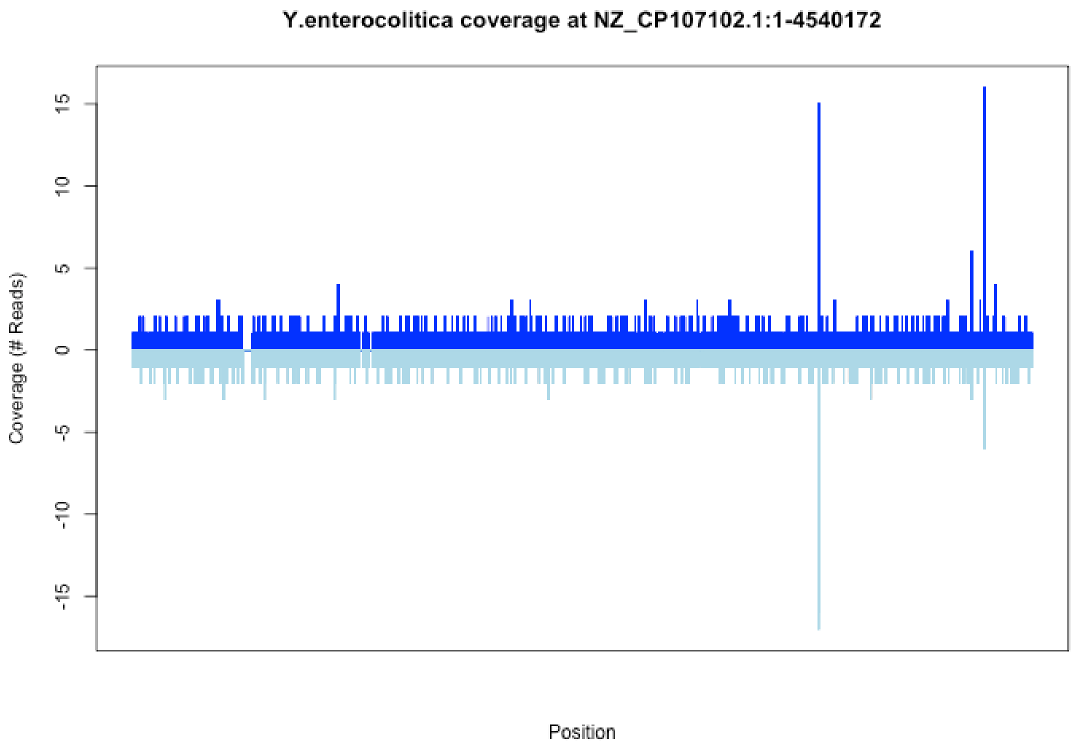

Figure S18. Distribution of mapped reads to *Y. enterocolitica* in SE_LN_ans010.The colors correspond to the coverage on the positive (dark blue) and negative (light blue) strands of the genome.

Regarding SE_PWC_ajv58, in the ED profiles for the 69 reads mapped to *Y. pestis* [Table S15] a non-specific pattern was observed, indicating that either *i*) this individual was not infected with *Y. pestis*, which remains hard to explain by the reads specifically mapping into the plasmids (from the 69 reads, 14 mapped to pPCP1 and 13 to pMT1), *ii)* it was highly DNA damaged or *iii)* contaminated with a related environmental species. Interestingly, sample SE_TRB_ans003 did show an ED profile highly consistent with *Y. pestis* [Fig. S19] and therefore we conclude that this individual was likely infected with this pathogen as well. Results for SE_TRB_ans005 and SE_TRB_ans007 remain in our opinion inconclusive and would require further sequencing to be able to determine if these individuals were infected with plague.

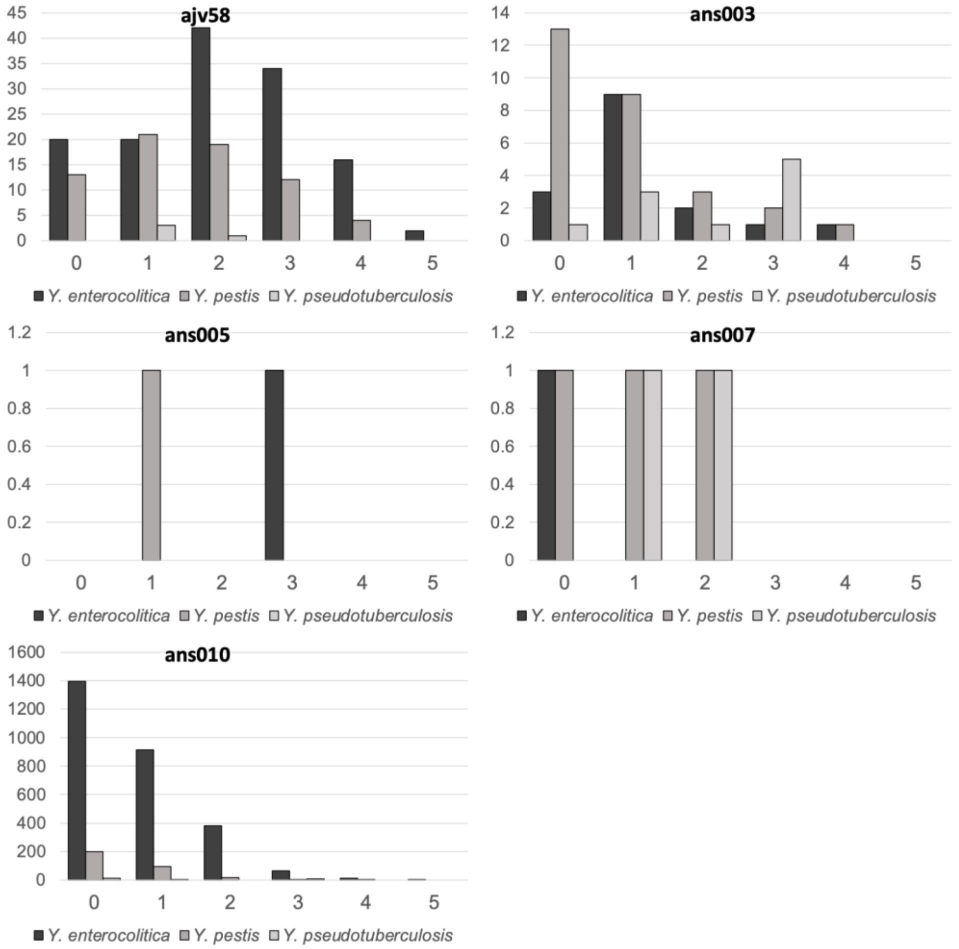

Figure S19. The observed edit-distance for each read map to each *Yersinia* reference genome, identifying the fragments with 0, 1, 2, 3, 4 and 5 mismatches for SE_PWC_ajv58, SE_TRB_ans003, SE_TRB_ans005, SE_TRB_ans007, and SE_LN_ans010.

Finally, we decided to explore the single nucleotide variants (SNVs) identified in the total reads mapping to *Y. pestis* in the five samples, to evaluate if they could provide some additional information regarding the relatedness of these strains to other ones from this species [Fig. S20]. We built a heat map plot with SNVs observed in all strains, excluding singletons (i.e., SNVs that were identified in a single sample), which not only reduced the plot size, but most importantly it reduced the noise from reads and variants originated from environmental contaminants. This analysis provided further evidence of the presence of *Y. pestis* in these samples by finding variants that are specifically characteristic of this species.

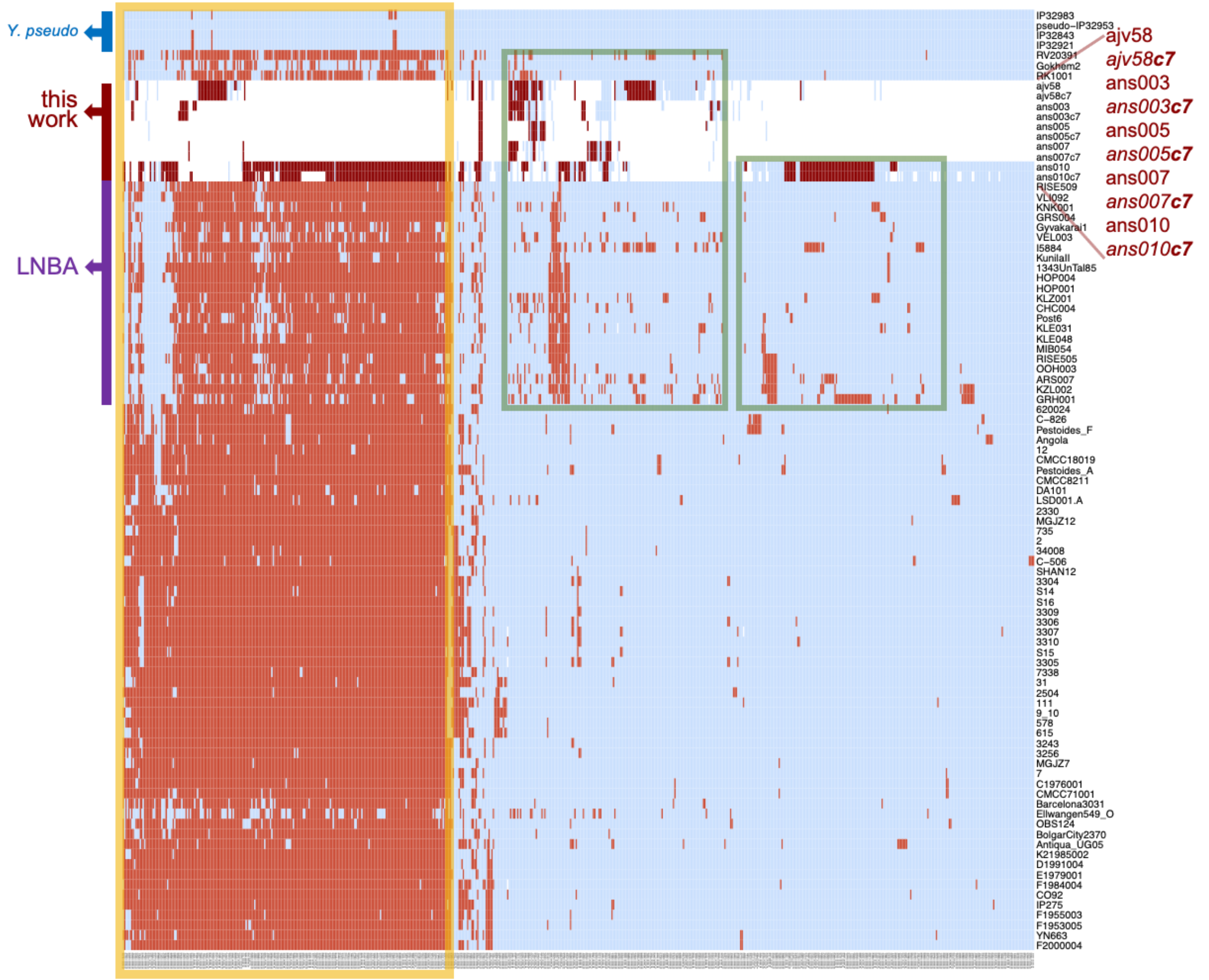

Figure S20. Heatmap showing informative genome wide single nucleotide variant sites from the five strains that were potentially positive for *Y. pestis*. The heatmap shows selected variant sites (columns) when genotyping against *Y. pseudotuberculosis IP32953* reference genome distributed among 79 publicly available genomes from *Y. pestis* (rows) and 4 *Y. pseudotuberculosis* (the four upper rows, marked in blue). The choice of this species as reference in this analysis is to get a better representation of genomic positions that may have been lost, to the more recent *Y. pestis* strains that are normally used as reference. Variants were ordered by clustering using the Euclidean distance and strains were ordered according to their phylogenetic positioning. These informative variant sites were selected by filtering only positions that were variant in at least two strains (i.e., singletons were excluded) and then keeping only positions covered by at least one of the mapping reads of the five potentially positive samples in this study. Variants uniquely found in the four *Y. pseudotuberculosis* strains were also excluded. Variant positions are indicated in red (darker red is used to highlight lines corresponding to the five samples of this study), non-variant in light blue (i.e., same genotype as in *Y. pseudotuberculosis* reference) and missing data in white. Variant sites that are characteristic of *Y. pestis* and not found in Y. pseudotuberculosis strains are marked with a yellow square. Variant positions that seem to be associated with the LNBA lineage are marked with green squares. All five positive samples are shown twice in the plot, including (***c7***) or not a clipping of 7bp at the end of reads, with highly similar results in in both cases, indicating that the identified variants are most likely not produced by DNA damage.

In agreement with previous results, the distribution of SNVs solidly indicated the presence of *Y. pestis* in the SE_LN_ans010 sample, as evidenced by the presence of numerous SNVs typically found in this species and not in *Y. pseudotuberculosis* [Fig. S20, yellow square]. Interestingly, this strain also shared several SNVs with LNBA strains that are not found in the RK1001, Gökhem2 and RV2039 strains (more basal in the phylogeny than LNBA), suggesting that the infecting strain belonged to the LNBA lineage, as it could be expected also by the temporality of this sample (2030-1890 calBCE, contemporaneous to other individuals infected with LNBA strains). While there were several SNVs supporting a *Y. pestis* identity for SE_TRB_ans003 [Fig. S20, yellow square], this was not the case for SE_TRB_ans005 and SE_TRB_ans007, for which results remained inconclusive in this analysis as well. Surprisingly, for the case of SE_PWC_ajv58, which had shown doubtful results for *Y. pestis* in the edit distance profiles, several SNVs were supportive for the presence of *Y. pestis* [Figure S20, yellow square], while many others were shared with the most ancestral and oldest strains (Gökhem2 and RV2039, [Fig. S20, central green square], suggesting that it may have been related to these basal lineages.

Among the four most basal strains (RV2039, Gok2, RK1001 and RISE509), the SE_PWC_ajv58 shared the highest number of SNVs with RV2039 (34 SNVs), followed by RK1001 (31 SNVs), markedly above the SNVs shared with Gökhem2 and RISE509 (19 and 21 SNVs respectively), suggesting that it could have been a strain related or closer to the lineage of RV2039. Interestingly, the SE_PWC_ajv58 was a hunter gatherer from the PWC culture dated in 3300-2300 cal BCE, which overlaps in time with the individual RV2039, a hunter-gatherer from Latvia dated at 3350-3100 cal BCE excavated in a site not far from the Gotland Island across the Baltic Sea (Susat et al., 2020). Although the information collected from SE_PWC_ajv58 is not sufficient to drive any solid conclusion, our results provide sufficient evidence to hypothesize that this individual was indeed infected with *Y. pestis*, with a strain likely pertaining to the most basal lineages of this species, which may have circulated among hunter-gatherers by the end of Neolithic.

### *References*

Adzhubei, I.A., Schmidt, S., Peshkin, L., Ramensky, V.E., Gerasimova, A., Bork, P., Kondrashov, A.S., Sunyaev, S.R., 2010. A method and server for predicting damaging missense mutations. Nat. Methods 7, 248–249. https://doi.org/10.1038/nmeth0410-248

Ahlström, T., Price, T.D., 2021. Mobile or stationary? An analysis of strontium and carbon isotopes from Västerbjers, Gotland, Sweden. J. Archaeol. Sci. Rep. 36, 102902. https://doi.org/10.1016/j.jasrep.2021.102902

Alexander, D.H., Novembre, J., Lange, K., 2009. Fast model-based estimation of ancestry in unrelated individuals. Genome Res. https://doi.org/10.1101/gr.094052.109

Allentoft, M.E., Sikora, M., Sjögren, K.-G., Rasmussen, S., Rasmussen, M., Stenderup, J., Damgaard, P.B., Schroeder, H., Ahlström, T., Vinner, L., Malaspinas, A.-S., Margaryan, A., Higham, T., Chivall, D., Lynnerup, N., Harvig, L., Baron, J., Casa, P.D., Dąbrowski, P., Duffy, P.R., Ebel, A.V., Epimakhov, A., Frei, K., Furmanek, M., Gralak, T., Gromov, A., Gronkiewicz, S., Grupe, G., Hajdu, T., Jarysz, R., Khartanovich, V., Khokhlov, A., Kiss, V., Kolář, J., Kriiska, A., Lasak, I., Longhi, C., McGlynn, G., Merkevicius, A., Merkyte, I., Metspalu, M., Mkrtchyan, R., Moiseyev, V., Paja, L., Pálfi, G., Pokutta, D., Pospieszny, Ł., Price, T.D., Saag, L., Sablin, M., Shishlina, N., Smrčka, V., Soenov, V.I., Szeverényi, V., Tóth, G., Trifanova, S.V., Varul, L., Vicze, M., Yepiskoposyan, L., Zhitenev, V., Orlando, L., Sicheritz-Pontén, T., Brunak, S., Nielsen, R., Kristiansen, K., Willerslev, E., 2015. Population genomics of Bronze Age Eurasia. Nature 522, 167–172. https://doi.org/10.1038/nature14507

Apel, J., Storå, J., Landeschi, G., 2015. Ett återbesök i Stora Förvar. Efterundersökning av stenålderslokalen Stora Förvar, Stora Karlsö, Eksta sn, RAÄ 138:1, Gotland (No. De gotländska pionjärbosättningarna: rapport nr 2). Institutionen för arkeologi och antikens historia Lunds universitet.

Apel, J., Wallin, P., Storå, J., Possnert, G., 2018. Early Holocene human population events on the island of Gotland in the Baltic Sea (9200-3800 cal. BP). Quat. Int. 465, Part B, 276–286. http://dx.doi.org/10.1016/j.quaint.2017.03.044

Arwidsson, G., 1979. Stenåldersmannen från Stora Bjärs i Stenkyrka. Arkeol. På Gotland, Gotlandica 14, Visby.

Arwidsson, G., 1949. Stenåldersfynden från Kambs i Lummelunda. Gotländskt Ark. 1948–1949, 147–167.

Ausmees, K., Sanchez-Quinto, F., Jakobsson, M., Nettelblad, C., 2022. An empirical evaluation of genotype imputation of ancient DNA. G3 GenesGenomesGenetics 12, jkac089. https://doi.org/10.1093/g3journal/jkac089

Auton, A., Brooks, L.D., Durbin, R.M., Garrison, E.P., Kang, H.M., Korbel, J.O., Marchini, J.L., McCarthy, S., McVean, G.A., Abecasis, G.R., 2015. A global reference for human genetic variation. Nature 526, 68–74. https://doi.org/10.1038/nature15393

Bägerfeldt, L., 1992. Neolitikum på Gotland. Problem och konsekvenser. ARKEO-Förlaget. Gamleby.

Bakker, J.A., 1979. The TRB West Group: studies in the chronology and geography of the makers of Hunebeds and Tiefstich pottery, 1979, Cingula. [Albert Egges van Giffen Instituut voor Prae- en Protohistorie, Amsterdam.

Burenhult, G., 2002. The grave-field at Ajvide, in: Burenhult, G. (Ed.), Remote Senisng, Vol. II, Thesis and Papers in North-European Archaeology. Department of Archaeology, Stockholm University, pp. 31–167.

Cassidy, L.M., Maoldúin, R.Ó., Kador, T., Lynch, A., Jones, C., Woodman, P.C., Murphy, E., Ramsey, G., Dowd, M., Noonan, A., Campbell, C., Jones, E.R., Mattiangeli, V., Bradley, D.G., 2020. A dynastic elite in monumental Neolithic society. Nature 582, 384–388. https://doi.org/10.1038/s41586-020-2378-6

Cassidy, L.M., Martiniano, R., Murphy, E.M., Teasdale, M.D., Mallory, J., Hartwell, B., Bradley, D.G., 2016. Neolithic and Bronze Age migration to Ireland and establishment of the insular Atlantic genome. Proc. Natl. Acad. Sci. 113, 368–373. https://doi.org/10.1073/pnas.1518445113

Chintalapati, M., Patterson, N., Moorjani, P., 2022. The spatiotemporal patterns of major human admixture events during the European Holocene. eLife 11, e77625. https://doi.org/10.7554/eLife.77625

Cingolani, P., Platts, A., Wang, L.L., Coon, M., Nguyen, T., Wang, L., Land, S.J., Lu, X., Ruden, D.M., 2012. A program for annotating and predicting the effects of single nucleotide polymorphisms, SnpEff. Fly (Austin) 6, 80–92. https://doi.org/10.4161/fly.19695

Coutinho, A., Günther, T., Munters, A.R., Svensson, E.M., Götherström, A., Storå, J., Malmström, H., Jakobsson, M., 2020. The Neolithic Pitted Ware culture foragers were culturally but not genetically influenced by the Battle Axe culture herders. Am. J. Phys. Anthropol. n/a. https://doi.org/10.1002/ajpa.24079

Danecek, P., Auton, A., Abecasis, G., Albers, C.A., Banks, E., DePristo, M.A., Handsaker, R.E., Lunter, G., Marth, G.T., Sherry, S.T., McVean, G., Durbin, R., 1000 Genomes Project Analysis Group, 2011. The variant call format and VCFtools. Bioinforma. Oxf. Engl. 27, 2156–2158. https://doi.org/10.1093/bioinformatics/btr330

Edenmo, R., 2008. Prestigeekonomi under yngre stenåldern: gåvoutbyten och regionala identiteter i den svenska båtyxekulturen, Occasional papers in archaeology. Inst. för Arkeologi och Antik Historia, University, dissertation, Uppsala.

Eriksson, G., 2004. Part-time farmers or hard-core sealers? Västerbjers studied by means of stable isotope analysis. J. Antropol. Archaeol. 23, 135–162.

Eriksson, G., Linderholm, A., Fornander, E., Kanstrup, M., Schoultz, P., Olofsson, H., Lidén, K., 2008. Same island, different diet: Cultural evolution of food practice on Öland, Sweden, from the Mesolithic to the Roman Period. J. Anthropol. Archaeol. 27, 520–543. https://doi.org/10.1016/j.jaa.2008.08.004

Excoffier, L., Marchi, N., Marques, D.A., Matthey-Doret, R., Gouy, A., Sousa, V.C., 2021. fastsimcoal2: demographic inference under complex evolutionary scenarios. Bioinforma. Oxf. Engl. 37, 4882–4885. https://doi.org/10.1093/bioinformatics/btab468

Fisher, A., Kristiansen, K., 2002. The Neolithisation of Denmark: 150 years of debate. J.R. Collis, Sheffield.

Fraser, M., 2018. People of the Dolmens and Stone Cists: An archaeogenetic Investigation of Megalithic Graves from the Neolithic Period on Gotland. AUN 47. (Doctoral thesis). Uppsala University-Campus Gotland, Department of Archaeology and Ancient History, Uppsala.

Fraser, M., Sanchez-Quinto, F., Evans, J., Storå, J., Götherström, A., Wallin, P., Knutsson, K., Jakobsson, M., 2018a. New insights on cultural dualism and population structure in the Middle Neolithic Funnel Beaker culture on the island of Gotland. J. Archaeol. Sci. Rep. 17, 325–334. https://doi.org/10.1016/j.jasrep.2017.09.002

Fraser, M., Sjödin, P., Sánchez-Quinto, F., Evans, J., Svedjemo, G., Knutsson, K., Götherström, A., Jakobsson, M., Wallin, P., Storå, J., 2018b. The Stone Cist Conundrum: A Multidisciplinary Approach to

Investigate Late Neolithic/ Early Bronze Age Population Demography on the Island of Gotland. J. Archaeol. Sci. Rep. 20 324–337.
Gazal, S., Sahbatou, M., Perdry, H., Letort, S., Génin, E., Leutenegger, A.-L., 2014. Inbreeding coefficient estimation with dense SNP data: comparison of strategies and application to HapMap III. Hum. Hered. 77, 49–62. https://doi.org/10.1159/000358224
Gimbutas, M., 1956. The prehistory of eastern Europe. Cambridge.
Günther, T., Malmström, H., Svensson, E.M., Omrak, A., Sánchez-Quinto, F., Kılınç, G.M., Krzewińska, M., Eriksson, G., Fraser, M., Edlund, H., Munters, A.R., Coutinho, A., Simões, L.G., Vicente, M., Sjölander, A., Sellevold, B.J., Jørgensen, R., Claes, P., Shriver, M.D., Valdiosera, C., Netea, M.G., Apel, J., Lidén, K., Skar, B., Storå, J., Götherström, A., Jakobsson, M., 2018. Population genomics of Mesolithic Scandinavia: Investigating early postglacial migration routes and high-latitude adaptation. PLOS Biol. 16, e2003703. https://doi.org/10.1371/journal.pbio.2003703
Günther, T., Valdiosera, C., Malmström, H., Ureña, I., Rodriguez-Varela, R., Sverrisdóttir, Ó.O., Daskalaki, E.A., Skoglund, P., Naidoo, T., Svensson, E.M., Bermúdez de Castro, J.M., Carbonell, E., Dunn, M., Storå, J., Iriarte, E., Arsuaga, J.L., Carretero, J.-M., Götherström, A., Jakobsson, M., 2015. Ancient genomes link early farmers from Atapuerca in Spain to modern-day Basques. Proc. Natl. Acad. Sci. 112, 11917–11922. https://doi.org/10.1073/pnas.1509851112
Haak, W., Lazaridis, I., Patterson, N., Rohland, N., Mallick, S., Llamas, B., Brandt, G., Nordenfelt, S., Harney, E., Stewardson, K., Fu, Q., Mittnik, A., Bánffy, E., Economou, C., Francken, M., Friederich, S., Pena, R.G., Hallgren, F., Khartanovich, V., Khokhlov, A., Kunst, M., Kuznetsov, P., Meller, H., Mochalov, O., Moiseyev, V., Nicklisch, N., Pichler, S.L., Risch, R., Rojo Guerra, M.A., Roth, C., Szécsényi-Nagy, A., Wahl, J., Meyer, M., Krause, J., Brown, D., Anthony, D., Cooper, A., Alt, K.W., Reich, D., 2015. Massive migration from the steppe was a source for Indo-European languages in Europe. Nature 522, 207–211. https://doi.org/10.1038/nature14317
Hallgren, F., 2008. Identitet i praktik: lokala, regionala och överregionala sociala sammanhang inom nordlig trattbägarkultur, Coast to coast books. Uppsala Univ., Department of Archaeology and Ancient History, Uppsala.
Herbig, A., Maixner, F., Bos, K.I., Zink, A., Krause, J., Huson, D.H., 2016. MALT: Fast alignment and analysis of metagenomic DNA sequence data applied to the Tyrolean Iceman. bioRxiv 050559. https://doi.org/10.1101/050559
Huson, D.H., Beier, S., Flade, I., Górska, A., El-Hadidi, M., Mitra, S., Ruscheweyh, H.-J., Tappu, R., 2016. MEGAN Community Edition - Interactive Exploration and Analysis of Large-Scale Microbiome Sequencing Data. PLOS Comput. Biol. 12, e1004957. https://doi.org/10.1371/journal.pcbi.1004957
Iversen, R., 2016a. Arrowheads as indicators of interpersonal violence and group identity among the Neolithic Pitted Ware hunters of southwestern Scandinavia. J. Anthropol. Archaeol. 44, 69–86. https://doi.org/10.1016/j.jaa.2016.09.004
Iversen, R., 2016b. Was There Ever a Single Grave Culture in East Denmark? Traditions and Transformations in the 3rd Millennium BC., in: Proceedings of the International Workshop “Socio-Environmental Dynamics over the Last 12,000 Years: The Creation of Landscapes III (15th – 18th April 2013)” in Kiel. Eds. Martin Furholt, Ralph Großmann, Marzena Szmyt. In Kommission bei Verlag Dr. Rudolf Habelt GmbH, Bonn, pp. 159–170.
Iversen, R., Philippsen, B., Persson, P., 2021. Reconsidering the Pitted Ware chronology: A temporal fixation of the Scandinavian Neolithic hunters, fishers and gatherers. Praehistorische Z. 96, 44–88. https://doi.org/10.1515/pz-2020-0033
Janzon, G.O., 1974. Gotlands mellanneolitiska gravar, Acta Universitatis Stockholmiensis. Studies in North-European Archaeology, 6. Stockh. : Almqvist & Wiksell.
Knutsson, K., 1988. Making and using stone tools : the analysis of the lithic assemblages from Middle Neolithic sites with flint in Västerbotten, northern Sweden, Aun, 0284-1347 ; 11. Societas archaeologica Upsaliensis :, Uppsala.
Kriiska, A., 2003. From hunter-fisher-gatherer to farmer - Changes in the Neolithic economy and settlement on Estonian territory. Archaeol. Lith. 4.

Kriiska, A., 2001. Stone Age Settlement and Economic Processes in the Estonian Coastal Area and Islands.

Kristiansen, K., 1989. Prehistoric Migrations—the Case of the Single Grave and Corded Ware Cultures. J. Dan. Archaeol. 8, 211–225. https://doi.org/10.1080/0108464X.1989.10590029

Kuhn, J.M.M., Jakobsson, M., Günther, T., 2018. Estimating genetic kin relationships in prehistoric populations. PLOS ONE 13, e0195491. https://doi.org/10.1371/journal.pone.0195491

Larsson, Å.M., 2009. Breaking and making bodies and pots: material and ritual practices in Sweden in the third millenium BC, Aun. Uppsala universitet, Uppsala.

Lazaridis, I., Nadel, D., Rollefson, G., Merrett, D.C., Rohland, N., Mallick, S., Fernandes, D., Novak, M., Gamarra, B., Sirak, K., Connell, S., Stewardson, K., Harney, E., Fu, Q., Gonzalez-Fortes, G., Jones, E.R., Roodenberg, S.A., Lengyel, G., Bocquentin, F., Gasparian, B., Monge, J.M., Gregg, M., Eshed, V., Mizrahi, A.-S., Meiklejohn, C., Gerritsen, F., Bejenaru, L., Blüher, M., Campbell, A., Cavalleri, G., Comas, D., Froguel, P., Gilbert, E., Kerr, S.M., Kovacs, P., Krause, J., McGettigan, D., Merrigan, M., Merriwether, D.A., O'Reilly, S., Richards, M.B., Semino, O., Shamoon-Pour, M., Stefanescu, G., Stumvoll, M., Tönjes, A., Torroni, A., Wilson, J.F., Yengo, L., Hovhannisyan, N.A., Patterson, N., Pinhasi, R., Reich, D., 2016. Genomic insights into the origin of farming in the ancient Near East. Nature 536, 419–424. https://doi.org/10.1038/nature19310

Lazaridis, I., Patterson, N., Mittnik, A., Renaud, G., Mallick, S., Kirsanow, K., Sudmant, P.H., Schraiber, J.G., Castellano, S., Lipson, M., Berger, B., Economou, C., Bollongino, R., Fu, Q., Bos, K.I., Nordenfelt, S., Li, H., de Filippo, C., Prüfer, K., Sawyer, S., Posth, C., Haak, W., Hallgren, F., Fornander, E., Rohland, N., Delsate, D., Francken, M., Guinet, J.-M., Wahl, J., Ayodo, G., Babiker, H.A., Bailliet, G., Balanovska, E., Balanovsky, O., Barrantes, R., Bedoya, G., Ben-Ami, H., Bene, J., Berrada, F., Bravi, C.M., Brisighelli, F., Busby, G.B.J., Cali, F., Churnosov, M., Cole, D.E.C., Corach, D., Damba, L., van Driem, G., Dryomov, S., Dugoujon, J.-M., Fedorova, S.A., Gallego Romero, I., Gubina, M., Hammer, M., Henn, B.M., Hervig, T., Hodoglugil, U., Jha, A.R., Karachanak-Yankova, S., Khusainova, R., Khusnutdinova, E., Kittles, R., Kivisild, T., Klitz, W., Kučinskas, V., Kushniarevich, A., Laredj, L., Litvinov, S., Loukidis, T., Mahley, R.W., Melegh, B., Metspalu, E., Molina, J., Mountain, J., Näkkäläjärvi, K., Nesheva, D., Nyambo, T., Osipova, L., Parik, J., Platonov, F., Posukh, O., Romano, V., Rothhammer, F., Rudan, I., Ruizbakiev, R., Sahakyan, H., Sajantila, A., Salas, A., Starikovskaya, E.B., Tarekegn, A., Toncheva, D., Turdikulova, S., Uktveryte, I., Utevska, O., Vasquez, R., Villena, M., Voevoda, M., Winkler, C.A., Yepiskoposyan, L., Zalloua, P., Zemunik, T., Cooper, A., Capelli, C., Thomas, M.G., Ruiz-Linares, A., Tishkoff, S.A., Singh, L., Thangaraj, K., Villems, R., Comas, D., Sukernik, R., Metspalu, M., Meyer, M., Eichler, E.E., Burger, J., Slatkin, M., Pääbo, S., Kelso, J., Reich, D., Krause, J., 2014. Ancient human genomes suggest three ancestral populations for present-day Europeans. Nature 513, 409–413. https://doi.org/10.1038/nature13673

Lindqvist, C., 1997. Ansarve hage-dösen. Tvärvetenskapliga aspekter på kontext och den neolitiska förändringen på Gotland, in: In A. Åkerlund, S. Bergh, J. Nordbladh, J.Taffinder (eds.). Till Gunborg. Arkeologiska samtal. Stockholm Archaeological Reports 33, pp. 361–378.

Lindqvist, C., Possnert, G., 1999. The first seal hunter families on Gotland. On the Mesolithic occupation of the Stora Förvar Cave. Current Swedish archaeology, 65–88.

Lindqvist, C., Possnert, G., 1997. The subsistence economy and diet at Jakobs/Ajvide and Stora Förvar, Eksta parish and other prehistoric dwelling and burial sites on Gotland in long-term perspective., in: In: Burenhult, G. (Ed.), Remote Sensing, Vol. I. Department of Archaeology, Stockholm Universit, Stockholm, pp. 29–90.

Lipson, M., Szécsényi-Nagy, A., Mallick, S., Pósa, A., Stégmár, B., Keerl, V., Rohland, N., Stewardson, K., Ferry, M., Michel, M., Oppenheimer, J., Broomandkhoshbacht, N., Harney, E., Nordenfelt, S., Llamas, B., Mende, B.G., Köhler, K., Oross, K., Bondár, M., Marton, T., Osztás, A., Jakucs, J., Paluch, T., Horváth, F., Csengeri, P., Koós, J., Sebők, K., Anders, A., Raczky, P., Regenye, J., Barna, J.P., Fábián, S., Serlegi, G., Toldi, Z., Nagy, E.G., Dani, J., Molnár, E., Pálfi, G., Márk, L., Melegh, B., Bánfai, Z., Domboróczki, L., Fernández-Eraso, J., Mujika-Alustiza, J.A., Fernández, C.A., Echevarría, J.J., Bollongino, R., Orschiedt, J., Schierhold, K., Meller, H., Cooper, A., Burger, J., Bánffy, E., Alt, K.W., Lalueza-Fox, C., Haak, W.,

Reich, D., 2017. Parallel palaeogenomic transects reveal complex genetic history of early European farmers. Nature 551, 368–372. https://doi.org/10.1038/nature24476

Lithberg, N., 1914. Gotlands stenålder. Bagge, Stockholm.

Malmer, M.P., 2002. The Neolithic of south Sweden: TRB, GRK, and STR. Royal Swedish Academy of Letters, History, and Antiquities : Distributed by Almquist & Wiksell International, Stockholm.

Malmer, M.P., 1975. Stridsyxekulturen i Sverige och Norge. LiberĽäromedel, Lund.

Malmer, M.P., 1962. Junge Neolitischen Studien, Acta archaeologica Lundensia in 8°, 2. Lund.

Malmström, H., Gilbert, M.T.P., Thomas, M.G., Brandström, M., Storå, J., Molnar, P., Andersen, P.K., Bendixen, C., Holmlund, G., Götherström, A., Willerslev, E., 2009. Ancient DNA Reveals Lack of Continuity between Neolithic Hunter-Gatherers and Contemporary Scandinavians. Curr. Biol. 19, 1758–1762. https://doi.org/10.1016/j.cub.2009.09.017

Malmström, H., Günther, T., Svensson, E.M., Juras, A., Fraser, M., Munters, A.R., Pospieszny, Ł., Tõrv, M., Lindström, J., Götherström, A., Storå, J., Jakobsson, M., 2019. The genomic ancestry of the Scandinavian Battle Axe Culture people and their relation to the broader Corded Ware horizon. Proc. R. Soc. B Biol. Sci. 286, 20191528. https://doi.org/10.1098/rspb.2019.1528

Malmström, H., Linderholm, A., Lidén, K., Stor\a a, J., Molnar, P., Holmlund, G., Jakobsson, M., Götherström, A., 2010. High frequency of lactose intolerance in a prehistoric hunter-gatherer population in northern Europe. BMC Evol. Biol. 10, 1.

Malmström, H., Linderholm, A., Skoglund, P., Stora, J., Sjodin, P., Gilbert, M.T.P., Holmlund, G., Willerslev, E., Jakobsson, M., Liden, K., Gotherstrom, A., 2015. Ancient mitochondrial DNA from the northern fringe of the Neolithic farming expansion in Europe sheds light on the dispersion process. Philos. Trans. R. Soc. B Biol. Sci. 370, 20130373–20130373. https://doi.org/10.1098/rstb.2013.0373

Marchi, N., Winkelbach, L., Schulz, I., Brami, M., Hofmanová, Z., Blöcher, J., Reyna-Blanco, C.S., Diekmann, Y., Thiéry, A., Kapopoulou, A., Link, V., Piuz, V., Kreutzer, S., Figarska, S.M., Ganiatsou, E., Pukaj, A., Struck, T.J., Gutenkunst, R.N., Karul, N., Gerritsen, F., Pechtl, J., Peters, J., Zeeb-Lanz, A., Lenneis, E., Teschler-Nicola, M., Triantaphyllou, S., Stefanović, S., Papageorgopoulou, C., Wegmann, D., Burger, J., Excoffier, L., 2022. The genomic origins of the world's first farmers. Cell 185, 1842-1859.e18. https://doi.org/10.1016/j.cell.2022.04.008

Martiniano, R., Cassidy, L.M., Ó'Maoldúin, R., McLaughlin, R., Silva, N.M., Manco, L., Fidalgo, D., Pereira, T., Coelho, M.J., Serra, M., Burger, J., Parreira, R., Moran, E., Valera, A.C., Porfirio, E., Boaventura, R., Silva, A.M., Bradley, D.G., 2017. The population genomics of archaeological transition in west Iberia: Investigation of ancient substructure using imputation and haplotype-based methods. PLOS Genet. 13, e1006852. https://doi.org/10.1371/journal.pgen.1006852

Martinsson-Wallin, H., 2008. Land and sea animal remains from Middle Neolithic Pitted Ware sites on Gotland Island in the Baltic Sea, Sweden. Isl. Inq. Colon. Seafar. Archaeol. Marit. Landsc. 29, 171.

Martinsson-Wallin, H., Wallin, P., 2010. The story of the only (?) megalith grave on Gotland Island., in: Budja, M. (Ed.), Dokumenta Praehistorica XXXVII. Ljubljana, pp. 77–84.

Mathieson, I., Lazaridis, I., Rohland, N., Mallick, S., Patterson, N., Roodenberg, S.A., Harney, E., Stewardson, K., Fernandes, D., Novak, M., Sirak, K., Gamba, C., Jones, E.R., Llamas, B., Dryomov, S., Pickrell, J., Arsuaga, J.L., de Castro, J.M.B., Carbonell, E., Gerritsen, F., Khokhlov, A., Kuznetsov, P., Lozano, M., Meller, H., Mochalov, O., Moiseyev, V., Guerra, M.A.R., Roodenberg, J., Vergès, J.M., Krause, J., Cooper, A., Alt, K.W., Brown, D., Anthony, D., Lalueza-Fox, C., Haak, W., Pinhasi, R., Reich, D., 2015. Genome-wide patterns of selection in 230 ancient Eurasians. Nature 528, 499–503. https://doi.org/10.1038/nature16152

McKenna, A., Hanna, M., Banks, E., Sivachenko, A., Cibulskis, K., Kernytsky, A., Garimella, K., Altshuler, D., Gabriel, S., Daly, M., DePristo, M.A., 2010. The Genome Analysis Toolkit: A MapReduce framework for analyzing next-generation DNA sequencing data. Genome Res. 20, 1297–1303. https://doi.org/10.1101/gr.107524.110

McVean, G.A., 2012. An integrated map of genetic variation from 1,092 human genomes. The 1000 genomes consortium. Nature 491, 56–65. https://doi.org/10.1038/nature11632

Midgley, M., 2008. The Megaliths of Northern Europe. Routhledge, London and New York.

Midgley, M.S., 1992. TRB culture: the first farmers of the North European plain. Edinburgh University Press, Edinburgh.

Mittnik, A., Wang, C.-C., Pfrengle, S., Daubaras, M., Zariņa, G., Hallgren, F., Allmäe, R., Khartanovich, V., Moiseyev, V., Tõrv, M., Furtwängler, A., Valtueña, A.A., Feldman, M., Economou, C., Oinonen, M., Vasks, A., Balanovska, E., Reich, D., Jankauskas, R., Haak, W., Schiffels, S., Krause, J., 2018. The genetic prehistory of the Baltic Sea region. Nat. Commun. 9, 442. https://doi.org/10.1038/s41467-018-02825-9

Molnar, P., 2008. Tracing prehistoric activitities: life ways, habitual behaviour and health of hunter-gatherers on Gotland, Theses and papers in osteoarchaeology. Dep. of archaeology and classical studies, Stockholm Univ, Stockholm.

Morgulis, A., Gertz, E.M., Schäffer, A.A., Agarwala, R., 2006. A fast and symmetric DUST implementation to mask low-complexity DNA sequences. J. Comput. Biol. J. Comput. Mol. Cell Biol. 13, 1028–1040. https://doi.org/10.1089/cmb.2006.13.1028

Müller, J., 2011. Megaliths and Funnel Beakers: Societies in Change 4100-2700 BC. Drieendertigste Kroon-Voordracht (Amsterdam 2011).

Narasimhan, V.M., Patterson, N., Moorjani, P., Rohland, N., Bernardos, R., Mallick, S., Lazaridis, I., Nakatsuka, N., Olalde, I., Lipson, M., Kim, A.M., Olivieri, L.M., Coppa, A., Vidale, M., Mallory, J., Moiseyev, V., Kitov, E., Monge, J., Adamski, N., Alex, N., Broomandkhoshbacht, N., Candilio, F., Callan, K., Cheronet, O., Culleton, B.J., Ferry, M., Fernandes, D., Freilich, S., Gamarra, B., Gaudio, D., Hajdinjak, M., Harney, É., Harper, T.K., Keating, D., Lawson, A.M., Mah, M., Mandl, K., Michel, M., Novak, M., Oppenheimer, J., Rai, N., Sirak, K., Slon, V., Stewardson, K., Zalzala, F., Zhang, Z., Akhatov, G., Bagashev, A.N., Bagnera, A., Baitanayev, B., Bendezu-Sarmiento, J., Bissembaev, A.A., Bonora, G.L., Chargynov, T.T., Chikisheva, T., Dashkovskiy, P.K., Derevianko, A., Dobeš, M., Douka, K., Dubova, N., Duisengali, M.N., Enshin, D., Epimakhov, A., Fribus, A.V., Fuller, D., Goryachev, A., Gromov, A., Grushin, S.P., Hanks, B., Judd, M., Kazizov, E., Khokhlov, A., Krygin, A.P., Kupriyanova, E., Kuznetsov, P., Luiselli, D., Maksudov, F., Mamedov, A.M., Mamirov, T.B., Meiklejohn, C., Merrett, D.C., Micheli, R., Mochalov, O., Mustafokulov, S., Nayak, A., Pettener, D., Potts, R., Razhev, D., Rykun, M., Sarno, S., Savenkova, T.M., Sikhymbaeva, K., Slepchenko, S.M., Soltobaev, O.A., Stepanova, N., Svyatko, S., Tabaldiev, K., Teschler-Nicola, M., Tishkin, A.A., Tkachev, V.V., Vasilyev, S., Velemínský, P., Voyakin, D., Yermolayeva, A., Zahir, M., Zubkov, V.S., Zubova, A., Shinde, V.S., Lalueza-Fox, C., Meyer, M., Anthony, D., Boivin, N., Thangaraj, K., Kennett, D.J., Frachetti, M., Pinhasi, R., Reich, D., 2019. The formation of human populations in South and Central Asia. Science 365. https://doi.org/10.1126/science.aat7487

Norderäng, J., 2008. 14C-dateringar från Ajvide. In Österholm (ed.), in: Jakobs/Ajvide: Undersökningar På En Gotländsk Boplatsudde Från Stenåldern. Gotland University Press, Visby, pp. 296–297.

Nordqvist, K., 2016. From Separation to Interaction: Corded Ware in the Eastern Gulf of Finland. Acta Archaeol. 87, 49–84. https://doi.org/10.1111/j.1600-0390.2016.12167.x

Olalde, I., Brace, S., Allentoft, M.E., Armit, I., Kristiansen, K., Booth, T., Rohland, N., Mallick, S., Szécsényi-Nagy, A., Mittnik, A., Altena, E., Lipson, M., Lazaridis, I., Harper, T.K., Patterson, N., Broomandkhoshbacht, N., Diekmann, Y., Faltyskova, Z., Fernandes, D., Ferry, M., Harney, E., De Knijff, P., Michel, M., Oppenheimer, J., Stewardson, K., Barclay, A., Alt, K.W., Liesau, C., Rios, P., Blasco, C., Miguel, J.V., Garcia, R.M., Fernandez, A.A., Banffy, E., Bernabo-Brea, M., Billoin, D., Bonsall, C., Bonsall, L., Allen, T., Buster, L., Carver, S., Navarro, L.C., Craig, O.E., Cook, G.T., Cunliffe, B., Denaire, A., Dinwiddy, K.E., Dodwell, N., Ernee, M., Evans, C., Kucharik, M., Farre, J.F., Fowler, C., Gazenbeek, M., Pena, R.G., Haber-Uriarte, M., Haduch, E., Hey, G., Jowett, N., Knowles, T., Massy, K., Pfrengle, S., Lefranc, P., Lemercier, O., Lefebvre, A., Martinez, C.H., Olmo, V.G., Ramirez, A.B., Maurandi, J.L., Majo, T., McKinley, J.I., McSweeney, K., Mende, B.G., Mod, A., Kulcsar, G., Kiss, V., Czene, A., Patay, R., Endrodi, A., Kohler, K., Hajdu, T., Szeniczey, T., Dani, J., Bernert, Z., Hoole, M., Cheronet, O., Keating, D., Veleminsky, P., Dobe, M., Candilio, F., Brown, F., Fernandez, R.F., Herrero-Corral, A.M., Tusa, S., Carnieri, E., Lentini, L., Valenti, A., Zanini, A., Waddington, C., Delibes, G., Guerra-Doce, E., Neil, B., Brittain, M., Luke, M., Mortimer, R., Desideri, J., Besse, M., Brucken, G.,

Furmanek, M., Hauszko, A., Mackiewicz, M., Rapinski, A., Leach, S., Soriano, I., Lillios, K.T., Cardoso, J.L., Pearson, M.P., Wodarczak, P., Price, T.D., Prieto, P., Rey, P.J., Risch, R., Guerra, M.A.R., Schmitt, A., Serralongue, J., Silva, A.M., Smrcka, V., Vergnaud, L., Zilhao, J., Caramelli, D., Higham, T., Thomas, M.G., Kennett, D.J., Fokkens, H., Heyd, V., Sheridan, A., Sjogren, K.G., Stockhammer, P.W., Krause, J., Pinhasi, R., Haak, W., Barnes, I., Lalueza-Fox, C., Reich, D., 2018. The Beaker phenomenon and the genomic transformation of northwest Europe. Nature 555, 190–196. https://doi.org/10.1038/nature25738

O'Leary, N.A., Wright, M.W., Brister, J.R., Ciufo, S., Haddad, D., McVeigh, R., Rajput, B., Robbertse, B., Smith-White, B., Ako-Adjei, D., Astashyn, A., Badretdin, A., Bao, Y., Blinkova, O., Brover, V., Chetvernin, V., Choi, J., Cox, E., Ermolaeva, O., Farrell, C.M., Goldfarb, T., Gupta, T., Haft, D., Hatcher, E., Hlavina, W., Joardar, V.S., Kodali, V.K., Li, W., Maglott, D., Masterson, P., McGarvey, K.M., Murphy, M.R., O'Neill, K., Pujar, S., Rangwala, S.H., Rausch, D., Riddick, L.D., Schoch, C., Shkeda, A., Storz, S.S., Sun, H., Thibaud-Nissen, F., Tolstoy, I., Tully, R.E., Vatsan, A.R., Wallin, C., Webb, D., Wu, W., Landrum, M.J., Kimchi, A., Tatusova, T., DiCuccio, M., Kitts, P., Murphy, T.D., Pruitt, K.D., 2016. Reference sequence (RefSeq) database at NCBI: current status, taxonomic expansion, and functional annotation. Nucleic Acids Res. 44, D733-745. https://doi.org/10.1093/nar/gkv1189

Orascanin, N., 2010. Den uppklädda människan: en diskussion kring den gropkeramiska klädesstilen (The dressed human : a discussion regarding the Pitted Ware clothing) (Archaeology, Bachelors degree). Högskolan på Gotland, Institutionen för kultur, energi och miljö, Visby.

Österholm, I., 1989. Bosättningsmönstret på Gotland under stenåldern. En analys av fysisk miljö, ekonomi och social miljö. Theses and papers in Archaeology 3. Stockholms universitet.

Palmgren, E., Martinsson-Wallin, H., 2015. Analysis of late mid-Neolithic pottery illuminates the presenceof a Corded Ware Culture on the Baltic Island of Gotland. Doc. Praehist. 42, 297-310-297–310. https://doi.org/10.4312/dp.42.21

Patterson, N., Moorjani, P., Luo, Y., Mallick, S., Rohland, N., Zhan, Y., Genschoreck, T., Webster, T., Reich, D., 2012. Ancient Admixture in Human History. Genetics 192, 1065. https://doi.org/10.1534/genetics.112.145037

Persson, P., 1999. Neolitikums början: undersökningar kring jordbrukets introduktion i Nordeuropa, Kust till kust-böcker. Dept. of Archeology, University of Gothenburg ; Dept. of Archaeology and Ancient History, University of Uppsala, Göteborg : Uppsala.

Pira, A., 1926. On bone deposits in the cave "Stora Förvar" in the Isle of Stora Karlsö, Sweden. Acta Zool. 7, 123–217. https://doi.org/10.1111/j.1463-6395.1926.tb00927.x

Price, T.D., 2015. Ancient Scandinavia: an archaeological history from the first humans to the Vikings. Oxford University Press, Oxford ; New York, NY.

Price, T.D., 2000. Europes First Farmers. Cambridge Univerity Press, New York.

Prüfer, K., Racimo, F., Patterson, N., Jay, F., Sankararaman, S., Sawyer, S., Heinze, A., Renaud, G., Sudmant, P.H., de Filippo, C., Li, H., Mallick, S., Dannemann, M., Fu, Q., Kircher, M., Kuhlwilm, M., Lachmann, M., Meyer, M., Ongyerth, M., Siebauer, M., Theunert, C., Tandon, A., Moorjani, P., Pickrell, J., Mullikin, J.C., Vohr, S.H., Green, R.E., Hellmann, I., Johnson, P.L.F., Blanche, H., Cann, H., Kitzman, J.O., Shendure, J., Eichler, E.E., Lein, E.S., Bakken, T.E., Golovanova, L.V., Doronichev, V.B., Shunkov, M.V., Derevianko, A.P., Viola, B., Slatkin, M., Reich, D., Kelso, J., Pääbo, S., 2013. The complete genome sequence of a Neanderthal from the Altai Mountains. Nature 505, 43–49. https://doi.org/10.1038/nature12886

Reich, D., Thangaraj, K., Patterson, N., Price, A.L., Singh, L., 2009. Reconstructing Indian population history. Nature 461, 489. https://doi.org/10.1038/nature08365

Ringbauer, H., Novembre, J., Steinrücken, M., 2021. Parental relatedness through time revealed by runs of homozygosity in ancient DNA. Nat. Commun. 12, 5425. https://doi.org/10.1038/s41467-021-25289-w

Rundkvist, M., Lindqvist, C., Thorsberg, K., 2004. Barshalder. 3: Rojrhage in Grötlingbo: a multi-component Neolithic shore site on Gotland, Stockholm archaeological reports. Dep. of Archaeology, Univ. of Stockholm, Stockholm.

Sánchez-Quinto, F., Malmström, H., Fraser, M., Girdland-Flink, L., Svensson, E.M., Simões, L.G., George, R., Hollfelder, N., Burenhult, G., Noble, G., Britton, K., Talamo, S., Curtis, N., Brzobohata, H., Sumberova, R., Götherström, A., Storå, J., Jakobsson, M., 2019. Megalithic tombs in western and northern Neolithic Europe were linked to a kindred society. Proc. Natl. Acad. Sci. 201818037. https://doi.org/10.1073/pnas.1818037116
Schiffels, S., Durbin, R., 2014. Inferring human population size and separation history from multiple genome sequences. Nat. Genet. 46, 919–925. https://doi.org/10.1038/ng.3015
Schnittger, B., 1913. Grottan Stora Förvar på Stora Karlsö. Cederquists grafiska aktiebolag, Stockholm.
Schnittger, B., Rydh, H., 1940. Grottan Stora Förvar på Stora Karlsö. Wahlström & Widmark., Stockholm.
Schroeder, H., Margaryan, A., Szmyt, M., Theulot, B., Włodarczak, P., Rasmussen, S., Gopalakrishnan, S., Szczepanek, A., Konopka, T., Jensen, T.Z.T., Witkowska, B., Wilk, S., Przybyła, M.M., Pospieszny, Ł., Sjögren, K.-G., Belka, Z., Olsen, J., Kristiansen, K., Willerslev, E., Frei, K.M., Sikora, M., Johannsen, N.N., Allentoft, M.E., 2019. Unraveling ancestry, kinship, and violence in a Late Neolithic mass grave. Proc. Natl. Acad. Sci. 116, 10705–10710. https://doi.org/10.1073/pnas.1820210116
Schultz Paulsson, B., 2017. Time and stone: the emergence and development of megaliths and megalithic societies in Europe. Archaeopress Archaeology, Oxford.
Sjögren, K.-G., 2003. Mångfalldige uhrminnes grafar: Megalitgravar och samhälle i Västsverige. GOTARC. Series B, Gothenburg archaeological theses 27 (Göteborg 2003).
Skoglund, P., Malmström, H., Omrak, A., Raghavan, M., Valdiosera, C., Günther, T., Hall, P., Tambets, K., Parik, J., Sjögren, K.-G., Apel, J., Willerslev, E., Storå, J., Götherström, A., Jakobsson, M., 2014. Genomic diversity and admixture differs for Stone-Age Scandinavian foragers and farmers. Science 344, 747–50. https://doi.org/10.1126/science.1253448
Skoglund, P., Malmstrom, H., Raghavan, M., Stora, J., Hall, P., Willerslev, E., Gilbert, M.T.P., Gotherstrom, A., Jakobsson, M., 2012. Origins and Genetic Legacy of Neolithic Farmers and Hunter-Gatherers in Europe. Science 336, 466–469. https://doi.org/10.1126/science.1216304
Sørensen, L., 2014. From hunter to farmer in northern Europe. Migration and adaption during the Neolithic and Bronze Age. Copenhagen.
Sørensen, L., Karg, S., 2014. The expansion of agrarian societies towards the north – new evidence for agriculture during the Mesolithic/Neolithic transition in Southern Scandinavia. J. Archaeol. Sci. 51, 98–114. https://doi.org/10.1016/j.jas.2012.08.042
Storå, J., 2001. Reading bones : Stone Age hunters and seals in the Baltic, Stockholm studies in archaeology, 0349-4128 ; 21. Univ., Stockholm.
Susat, J., Bonczarowska, J.H., Pētersone-Gordina, E., Immel, A., Nebel, A., Gerhards, G., Krause-Kyora, B., 2020. Yersinia pestis strains from Latvia show depletion of the pla virulence gene at the end of the second plague pandemic. Sci. Rep. 10, 14628. https://doi.org/10.1038/s41598-020-71530-9
Tilley, C., 1999. The dolmen and passage graves of Sweden. An introduction and guide. Institute of Archaeology, ULC.
Vanhanen, S., Gustafsson, S., Ranheden, H., Björck, N., Kemell, M., Heyd, V., 2019. Maritime Hunter-Gatherers Adopt Cultivation at the Farming Extreme of Northern Europe 5000 Years Ago. Sci. Rep. 9, 4756. https://doi.org/10.1038/s41598-019-41293-z
von Hackwitz, K., 2009. Längs med Hjälmarens stränder och förbi : relationen mellan den gropkeramiska kulturen och båtyxekulturen. Institutionen för arkeologi och antikens kultur, Stockholms universitet, Stockholm.
Wallin, P., 2016. The Use and Organisation of a Middle-Neolithic Pitted Ware Coastal Site on the Island of Gotland in the Baltic Sea. In Catherine Dupont et Gregor Marchand (Eds.). Archaeology of maritime hunter-gatherers. From settlement function to the organization of the coastal zone Actes de la séance de la Société préhistorique française de Rennes, 10-11 avril 2014. Paris, Société préhistorique française.
Wallin, P., 2015. A perfect Death: Examples of Pitted Ware Ritualization of the Dead., in: In K von Hackwitz and R. Peyroteo Stjerna (Eds.). Ancient Death Ways: Proceedings of the Workshop on Archaeology

and Mortuary Practices, Uppsala, 16-17 May 2013, Occational Papers in Archaeology. Uppsala Universitet, Uppsala, pp. 47–64.

Wallin, P., 2010. Neolithic monuments on Gotland: Material expressions of the domestication process., in: In H. Martinsson-Wallin (Ed.), Baltic Prehistoric Interactions and Transformations: The Neolithic to the Bronze Age. Gotland University Press. No. 5. Visby, pp. 39–61.

Wallin, P., Martinsson-Wallin, H., 2016. Collective spaces and material expressions: ritual practice and island identities in Neolithic Gotland, in: In.Eds. George Nash & Andrew Townsend, Decoding the Neolithic and Mediterranean Island Ritual. Oxford Books, Philadelphia, pp. 1–15.

Wallin, P., Martinsson-Wallin, H., 1997. Osteological analysis of skelletal remains from a megalithic grave in Ansarve, Tofta Parish, Gotland, in: In G. Burenhult (Ed.), Remote Sensing, Vol. 1. Thesis and papers in North-European Archaeology 13:a. Stockholm University, Stockholm, pp. 23–28.

Wallin, P., Sjöstrand, A., 2018. Rapport från arkeologisk undersökning vid Licksarve 2:1, Raä Tofta 27:1, Gotland. (Arkeologisk fältrapport No. 20). Uppsala University.

Wallin, P., Wehlin, J., 2010. Räddad, reformerad och registrerad – en ”grävande” reviderande rapport rörande en fornlämning i Tofta, in: In. Eds. Ridder & Sandström, Gotlandsakademiker Tycker Om...2010. Gotland University Press, pp. 23–33.

Widerström, P., Norderäng, J., 2020. Rapport Hamra Långmyre- En tidigare okänd stenålderslokal (No. Länsstyrelsens Dnr 431-4155-19). Gotlands Museum, Visby.
