## Supplementary figures and images for "Ancestry, admixture, and pathogens in contemporaneous Neolithic farmers and foragers on the Island of Gotland"

### Extended Data Figure1

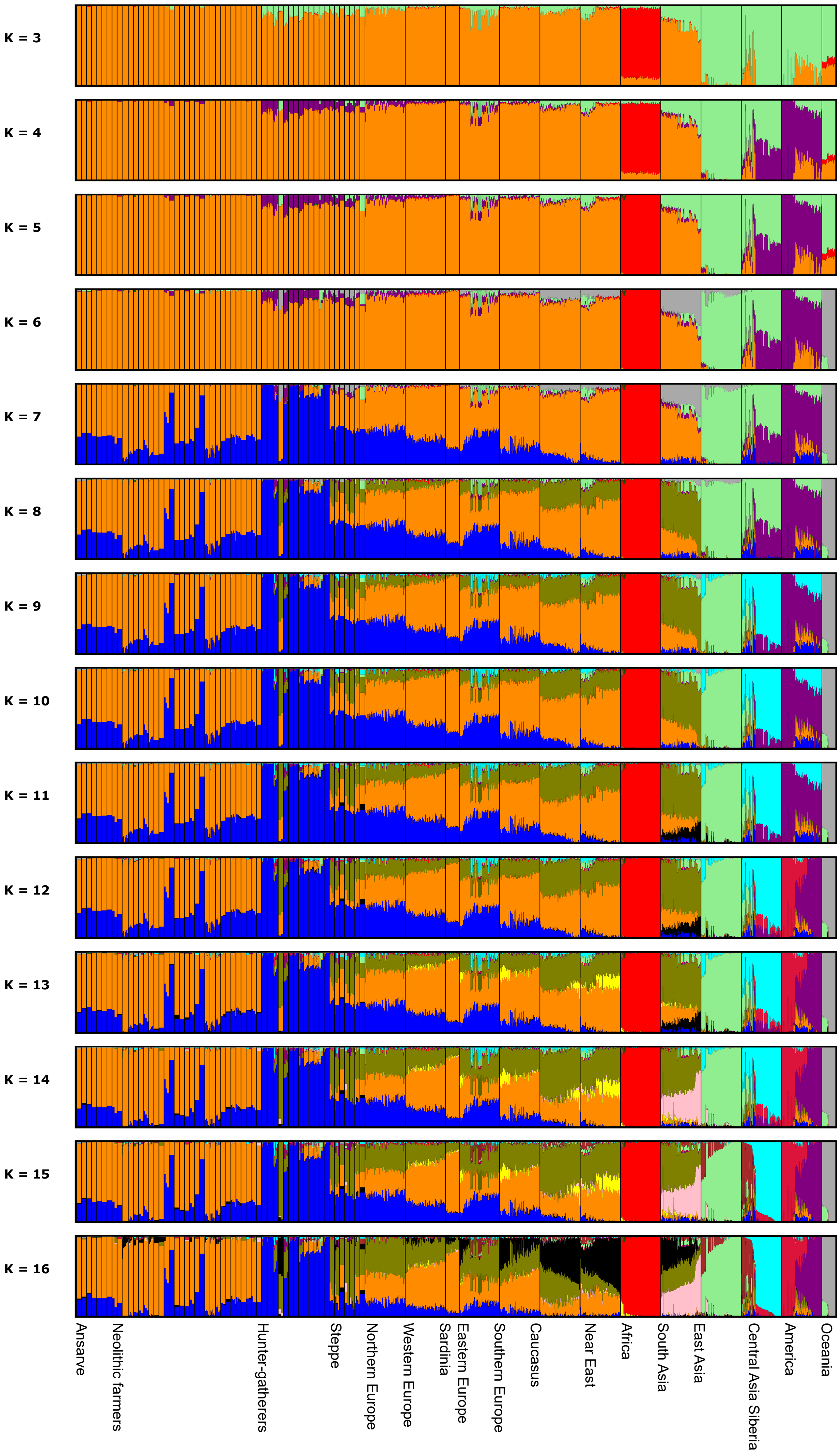


The ancient individuals are in the same order as Fig. 2A
